# The conquest and diversification of leafy spurges across the Holarctic and beyond: biogeography and evolution of life-history of *Euphorbia* subgenus *Esula*

**DOI:** 10.1101/2025.09.04.673735

**Authors:** Irene Masa-Iranzo, Isabel Sanmartín, Božo Frajman, Andrea S. Meseguer, Ricarda Riina

## Abstract

*Euphorbia* subgenus *Esula* is one of four main lineages within the megadiverse angiosperm genus *Euphorbia*. It comprises 490 species, including herbs, shrubs, dendroid shrubs, and succulents. The subgenus is a northern temperate lineage, most diverse in the Irano-Turanian and Mediterranean regions. We assembled the largest taxon sampling of subg. *Esula* to date (321 spp.), using DNA sequences from nuclear ITS and plastid *ndhF* regions, and updated previous phylogenetic analyses and sectional classification of the group. We used Bayesian methods to estimate divergence times and reconstruct ancestral ranges, characterising the tempo and mode of lineage diversification that shaped the present distribution and extant diversity of the subgenus. We tested whether changes in life-history (annual vs. perennial) played a role in the diversification and current elevational distribution of subg. *Esula*. Our results show that subg. *Esula* diverged from its sister clade in the Mid-Eocene and started to diversify in the Late Eocene. The Western Palearctic was inferred to be the ancestral area of the subgenus, where the first diversification events began 41 Mya. Expansion to other continents was dated as occurring in the last 10 million years. This coincided with an increase in the diversification rate and with clade-specific rate shifts coupled with changes from annual to perennial life-histories. We found a positive correlation between perenniality and high elevations. These results support that the recent rapid diversification of subg. *Esula* could be associated with an evolutionary shift to perenniality, allowing colonisation of montane habitats and global range expansion beyond the Western Palearctic.

## 1 Introduction

Disentangling the origins and evolution of the Northern Hemisphere biomes remains an important goal of modern botany (Donoghue & Edwards, 2014; Adhikari et al., 2015). This question is intricately linked to our understanding of the evolutionary history of lineages across space and time, and their common assembly into biogeographic regions or realms. Within the Holarctic realm, complex biogeographic narratives, involving ancient vicariance events and more recent dispersal between the Palearctic and Nearctic regions, have been hypothesised for many lineages. These hypotheses attribute species diversification to the interplay of plate tectonics and associated climatic shifts (Wen, 1999; Morley, 2011; Kleckova et al., 2015; Hansen et al., 2023).

The Western Palearctic region within the Holarctic realm, i.e. the Palearctic west of the Ural Mountains (Sanmartín et al., 2001), underwent the most dynamic tectonic and climatic history in the area (summarised in Table 1). In the Early Paleocene (∼65 Mya, Table 1: GI1), the African Plate on its northeastward drift collided with the Iberian Peninsula, closing the western side of the Mesozoic Tethys Seaway. The rise of the proto-Alps led to the formation of the Eastern Paratethys Sea at the Eocene-Oligocene boundary (∼35 Mya) and partially isolated it from the Central Tethys (Meulenkamp & Sissing, 2003). The climate of the Holarctic landmasses was warm and humid during most of the Paleogene (∼65–40 Mya) and climaxed during the Paleocene-Eocene Thermal Maximum (PETM, ∼55 Mya; Zachos et al., 2001). At this time, a broad-leaved, evergreen vegetation belt, the boreotropical forest, extended across the northern latitudes (Wolfe, 1978; Tiffney, 1985a; Morley, 2003). The southern Western Palearctic probably had drier, sclerophyllous Madrean-Tethyan vegetation (Axelrod, 1975). At the Eocene-Oligocene boundary (Terminal Eocene Event, TEE, ∼34 Mya; Tiffney, 1985a), with the onset of colder and more arid climates, the boreotropical forests evolved into more deciduous, mixed-mesophytic vegetation (Table 1: GI1). In the Early Miocene (Table 1: GI2), the Arabian Plate, an African promontory, collided with Eurasia on its eastern side, closing the connection of the Tethys Seaway with the Indian Ocean (Terminal Tethyan Event, TTE, ∼20–18 Mya; Liu et al., 2018). This, together with a prior increase in temperatures, the Late Oligocene Warming Event (LOWE, ∼28 Mya; Hansen et al., 2008), introduced drier continental climates in the Western Palearctic. A continuous landmass was formed, separating the Paratethys from the Central Tethys, allowing biotic exchange between the eastern and western proto-Mediterranean (Meulenkamp & Sissingh, 2003). Another consequence of the Arabian-Eurasian collision was the rise of the Anatolian and Iranian plateaus, which started ∼20 Mya and underwent geographically asynchronous periods of uplift (Table 1: GI2; Manafzadeh et al., 2017). Following the Mid-Miocene Climatic Optimum (MMCO, ∼16–14 Mya; Zachos et al., 2008), the Western Palearctic experienced a more arid and colder climate. This change was triggered by the uplift of the Iranian Plateau and the Alborz, Kopet Dagh, and Zagros mountain ranges, as well as the encroachment of the Paratethys and the proto-Mediterranean seas (Table 1). A continental steppe vegetation replaced the temperate mixed-mesophytic in Eastern Europe and Central Asia (Ivanov et al., 2011). The end of the Miocene was characterised by colder climates (Late Miocene Cooling Event, LMCE, ∼11–7 Mya; Zachos et al., 2008), driven partly by further uplift of the Iranian Plateau (Manafzadeh et al., 2017), the formation of the Sahara Desert (∼8 Mya; Sepulchre et al., 2006), and the almost complete desiccation of the Mediterranean Sea (Messinian Salinity Crisis, MSC, ∼6–5.3 Mya; Krijgsman et al., 2010). These conditions favoured the evolution of continental-adapted plants, which became dominant in the Iranian Plateau and the Mediterranean Basin (Manafzadeh et al., 2017). The opening of the Strait of Gibraltar (∼3.5 Mya) and a new period of climatic aridity (Mid-Pliocene Warming Event, MPWE, ∼3.5 Mya; Zachos et al., 2008) led to the onset of the Mediterranean biome (Thompson, 2005) and the expansion of xerophytic taxa across the southwestern Palearctic (Table 1). Other important events for the biogeography of the Western Palearctic lineages were the closing and opening of seaways. For example, the closure of the Turgai Strait at 30 Mya allowed biotic migration towards and from the Eastern Palearctic (Tiffney, 1985b). A widening North Atlantic Ocean isolated the Western Palearctic from the Nearctic at the start of the Cenozoic, but connections were possible through land bridges until 50 or 20 Mya, depending on the literature (Sanmartín et al., 2001; Brikiatis, 2014). Another event which had a profound influence on the climate and vegetation of the Western Palearctic was the uplift of the Qinghai-Tibetan Plateau (QTP) within the Himalayan range (Favre et al., 2015). The rise of the QTP was caused by the collision of the Indian Plate with the Eurasian Plate and involved four major stages: QTP1 (∼25–17 Mya), QTP2 (∼15–13 Mya), QTP3 (∼8–7 Mya), and QTP4 (∼3.5–1.7 Mya) (Guo et al., 2002).

**Table 1.**
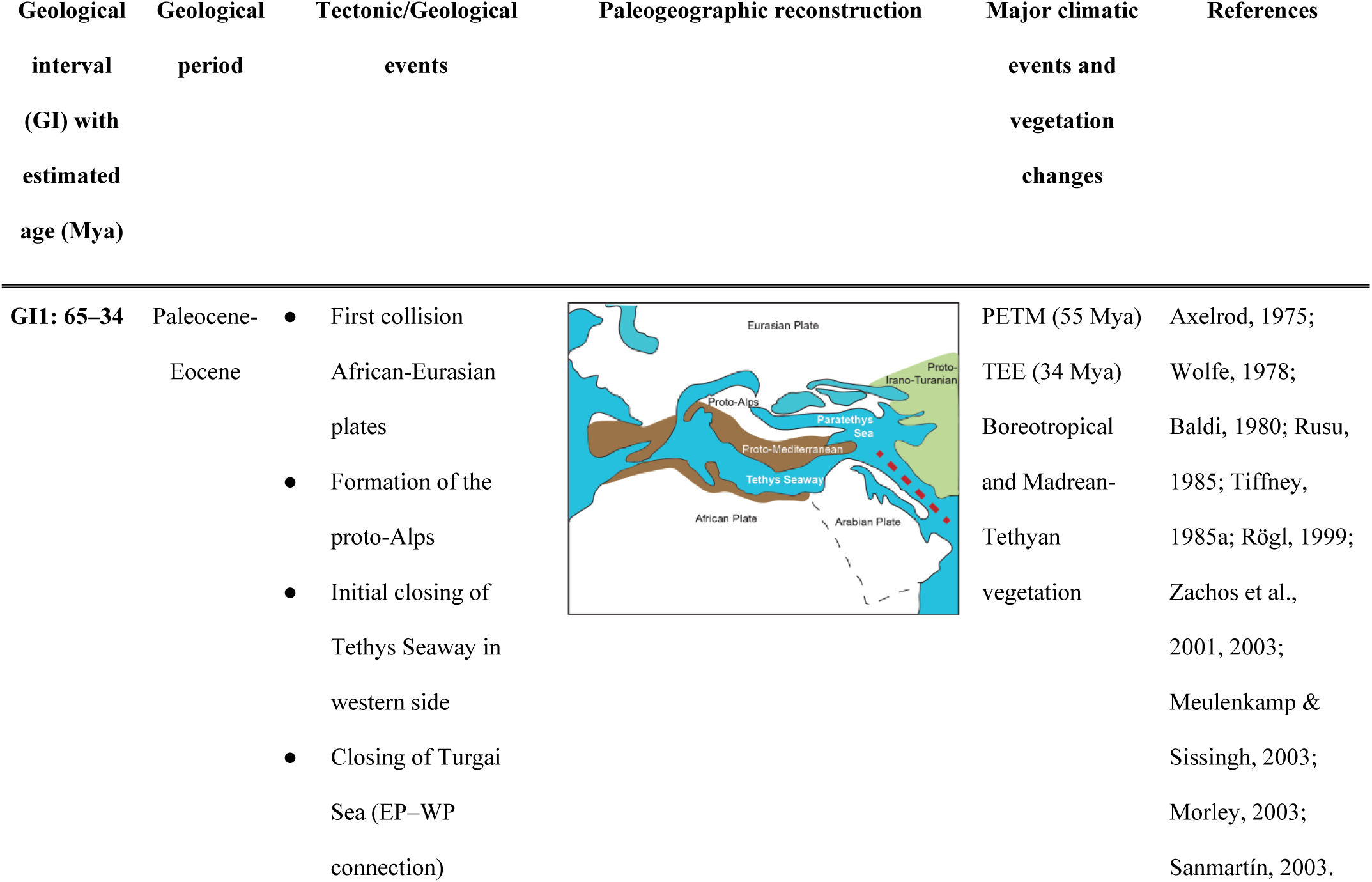
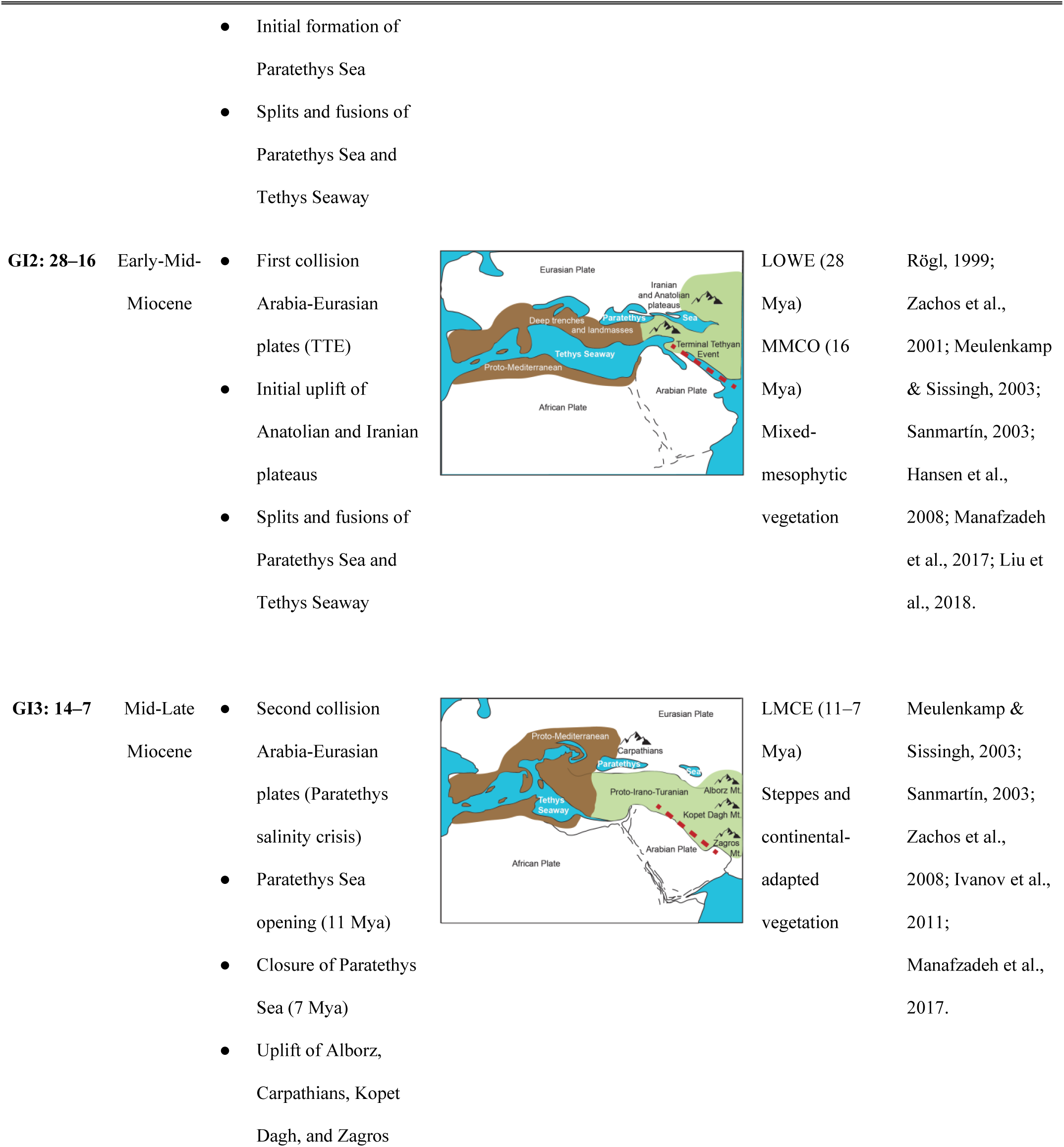
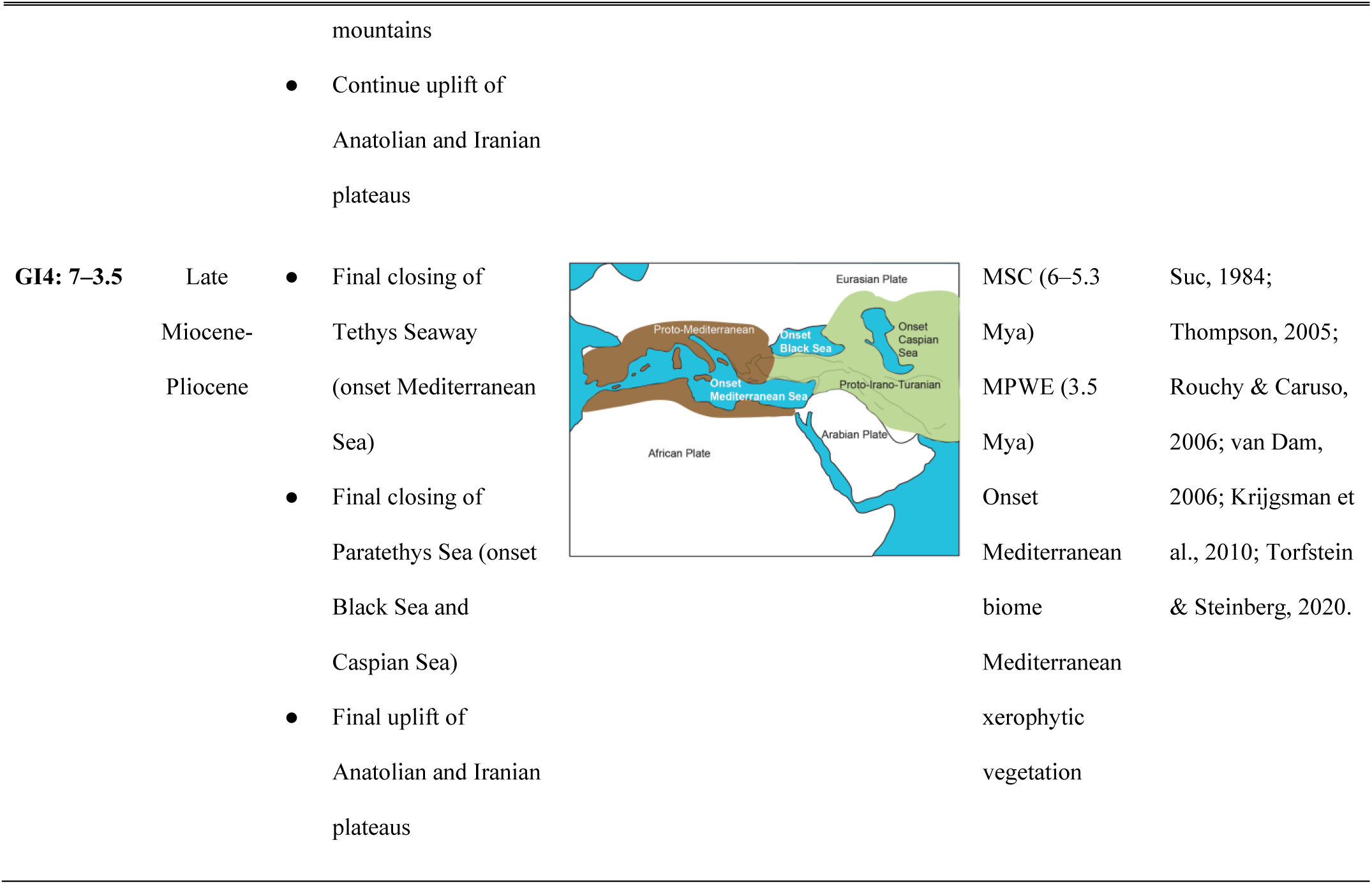
Main geological and climatic events ordered by their estimated age and associated changes in vegetation cover in the Western Palearctic during the Cenozoic. Depending on the reference, the dates of the events may vary. The maps show paleogeographic reconstructions of the Western Palearctic terranes and marine basins throughout the Cenozoic, reflecting the main collision stages of the African and Eurasian plates (adapted from Ree & Sanmartín, 2009). Acronyms: EP, Eastern Palearctic; LMCE, Late Miocene Cooling Event; LOWE, Late Oligocene Warming Event; MMCO, Mid-Miocene Climatic Optimum; MPWE, Mid-Pliocene Warming Event; MSC, Messinian Salinity Crisis; PETM, Paleocene-Eocene Thermal Maximum; TEE, Terminal Eocene Event; TTE, Terminal Tethyan Event; WP, Western Palearctic.

| Geological interval (GI) with estimated age (Mya) | Geological period | Tectonic/Geological events | Paleogeographic reconstruction | Major climatic events and vegetation changes | References |
| --- | --- | --- | --- | --- | --- |
| **GI1: 65–34** | Paleocene-Eocene | First collision African-Eurasian plates<br>Formation of the proto-Alps<br>Initial closing of Tethys Seaway in western side<br>Closing of Turgai Sea (EP–WP connection) |  | PETM (55 Mya)<br>TEE (34 Mya)<br>Boreotropical and Madrean-Tethyan vegetation | Axelrod, 1975;<br>Wolfe, 1978;<br>Baldi, 1980; Rusu, 1985; Tiffney, 1985a; Rögl, 1999;<br>Zachos et al., 2001, 2003;<br>Meulenkamp & Sissingh, 2003;<br>Morley, 2003;<br>Sanmartín, 2003. |
|  |  | Initial formation of Paratethys Sea<br>Splits and fusions of Paratethys Sea and Tethys Seaway |  |  |  |
| **GI2: 28–16** | Early-Mid-Miocene | First collision Arabia-Eurasian plates (TTE)<br>Initial uplift of Anatolian and Iranian plateaus<br>Splits and fusions of Paratethys Sea and Tethys Seaway |  | LOWE (28 Mya)<br>MMCO (16 Mya)<br>Mixed-mesophytic vegetation | Rögl, 1999;<br>Zachos et al., 2001; Meulenkamp & Sissingh, 2003;<br>Sanmartín, 2003;<br>Hansen et al., 2008; Manafzadeh et al., 2017; Liu et al., 2018. |
| **GI3: 14–7** | Mid-Late Miocene | Second collision Arabia-Eurasian plates (Paratethys salinity crisis)<br>Paratethys Sea opening (11 Mya)<br>Closure of Paratethys Sea (7 Mya)<br>Uplift of Alborz, Carpathians, Kopet Dag, and Zagros<br>mountains |  | LMCE (11–7 Mya)<br>Steppes and continental-adapted vegetation | Meulenkamp & Sissingh, 2003;<br>Sanmartín, 2003;<br>Zachos et al., 2008; Ivanov et al., 2011;<br>Manafzadeh et al., 2017. |
|  |  | Continue uplift of Anatolian and Iranian plateaus |  |  |  |
| **GI4: 7–3.5** | Late Miocene-Pliocene | Final closing of Tethys Seaway (onset Mediterranean Sea)<br>Final closing of Paratethys Sea (onset Black Sea and Caspian Sea)<br>Final uplift of Anatolian and Iranian plateaus |  | MSC (6–5.3 Mya)<br>MPWE (3.5 Mya)<br>Onset Mediterranean biome<br>Mediterranean xerophytic vegetation | Suc, 1984;<br>Thompson, 2005;<br>Rouchy & Caruso, 2006; van Dam, 2006; Krijgsman et al., 2010; Torfstein & Steinberg, 2020. |

Associated with the complex and dynamic geological history of the Northern Hemisphere, many Holarctic lineages evolved common strategies to cope with major climatic changes resulting from the aforementioned tectonic rearrangements. Several studies have shown that the transition from wet tropical forests to more xeric habitats (temperate forests, steppes, Mediterranean shrubland, etc.) was often accompanied by the evolution of complex physiological traits aimed to withstand freezing and dry conditions. These traits include shedding leaves during freezing periods or senescing above-ground tissues, as well as major physiological changes, such as the evolution of annual or perennial life-history strategies (Donoghue & Edwards, 2014; Lindberg et al., 2020). It is thought that the shift from a perennial to an annual life-cycle allows lineages to escape adverse seasons (e.g. periods of extreme heat and drought), while the shift to perenniality promotes colonisation of habitats at higher elevations and latitudes (Drummond et al., 2012a; Zanne et al., 2014; Boyko et al., 2023). Plant families containing annuals are distributed throughout the entire angiosperm tree of life, suggesting that this trait has evolved multiple times independently (Hjertaas et al., 2023). Historically, transitions between life-history strategies have been thought to occur from perennial ancestors to annual descendants (Jeffrey, 1916; Stebbins, 1982; see also Hjertaas et al., 2023). However, several well-documented phylogenetic studies have inferred the opposite pattern: the evolution of perennials from annual ancestors, for example, in the tribes Delphinieae (Jabbour & Renner, 2012) and Fabeae (Schaefer et al., 2012), or the genera *Androsace* L. (Schneeweiss et al., 2004), *Castilleja* Mutis ex L.f. (Tank & Olmstead, 2008), *Knautia* L. (Rešetnik et al., 2014), *Lupinus* L. (Drummond et al., 2012a), and *Medicago* L. (Bena et al., 1998).

*Euphorbia* subgenus *Esula* Pers. (Euphorbiaceae), commonly known as the leafy spurges, is one of the most emblematic elements of the Western Palearctic flora. It comprises 490 species that exhibit a wide range of growth forms, from herbs (annuals or perennials) to shrubs, dendroid shrubs, and even some succulent species (Riina et al., 2013; Heimer & Frajman, 2023). This subgenus belongs to one of the largest angiosperm genera, *Euphorbia* L., including more than 2,150 species distributed worldwide (Horn et al., 2012; WFO, 2024). The other three subgenera of *Euphorbia* are *E.* subg. *Athymalus* Neck. ex Rchb. (150 spp.), *E*. subg. *Chamaesyce* Raf. (600 spp.), and *E*. subg. *Euphorbia* (700 spp.) (Yang et al., 2012; Dorsey et al., 2013; Peirson et al., 2013). Subgenus *Esula* represents the primary northern temperate radiation of *Euphorbia*, while the other three subgenera consist of mostly tropical and subtropical lineages with major centres of diversity in Africa and the Americas. Though members of subg. *Esula* can be found in Africa (47 spp.), the New World (37 spp.), Macaronesia (24 spp.), and the Indo-Pacific region (3 spp.), the subgenus is most diverse in the Western Palearctic region (268 spp.), followed by the Eastern Palearctic (169 spp.) (Fig. 1). Within the Palearctic region, the Mediterranean Basin and the Irano-Turanian region harbour the highest number of species (Fig. 1; Geltman, 2015; Pahlevani et al., 2020). The subgenus includes also some species that are considered noxious weeds, growing outside their native ranges, such as *E. virgata* Waldst. & Kit. (commonly misidentified as *E. esula* L.; Berry et al., 2016), *E. oblongata* Griseb., and *E. terracina* L. (DiTomaso & Healy, 2007; Smith Jr, 2019). Similar to the other three subgenera of *Euphorbia*, subg. *Esula* displays variability in life-histories; while most species are perennials (413 spp., 84.29%), there is also a moderate number of annuals (77 spp., 15.71%). Horn et al. (2012) and Anest et al. (2021) reconstructed the most recent common ancestor (MRCA) of the subgenus as a herbaceous plant, but did not consider variation in life-history (annual or perennial). Frajman & Schönswetter (2011) inferred that a transition from perennial to annual life-history occurred independently in several lineages. However, sampling was relatively low (99 spp., 20%, and only 10 out of 21 sections) and the life-history state of the ancestor of subg. *Esula* could not be inferred with any certainty.

**Fig. 1.**
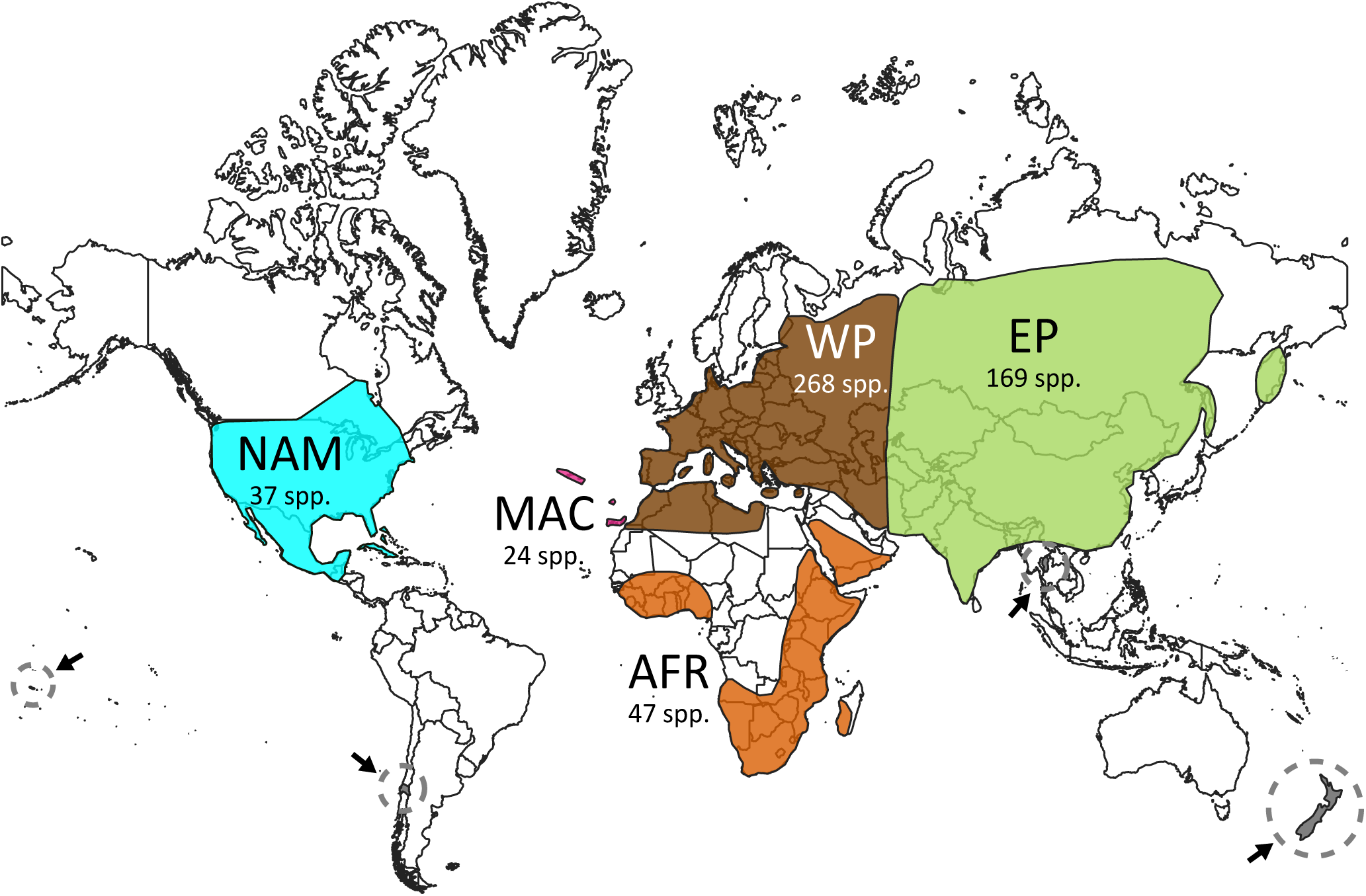
Schematic map showing the distribution of *Euphorbia* subg. *Esula* based on current knowledge (see Methods and Table S1). Colours indicate the six areas used in the biogeographic analyses: AFR, Africa; EP, Eastern Palearctic; MAC, Macaronesia; NAM, North America, Central America, and Caribbean region; WP, Western Palearctic. Grey-dashed circles with an arrow indicate the areas (Chile, New Zealand, Samoa, and Thailand) with one species each, not included in our biogeographic analyses.

In the last decades, molecular phylogenetic studies based on a limited number of DNA regions from the nuclear and plastid genomes have consolidated the monophyly of subg. *Esula*, as well as its phylogenetic structure and systematic classification (Steinmann & Porter, 2002; Frajman & Schönswetter, 2011; Horn et al., 2012; Riina et al., 2013). The most comprehensive phylogenetic analysis of the subgenus in terms of number of taxa is Riina et al. (2013), who sequenced the nuclear ITS marker and the plastid *ndhF* region for 273 species (55.71%). Horn et al. (2012), in their comprehensive study of the entire genus *Euphorbia*, included more loci (10 nuclear and plastid regions); however, their sampling within subg. *Esula* was considerably low (8%). Other studies have contributed to the taxonomic knowledge of the subgenus, through the description of new species or a better delimitation of taxonomic entities within species complexes or poorly known lineages (e.g. Mayfield, 2013; Pahlevani, 2017a; Bruyns, 2019; Stevanoski et al., 2020). Most of these recently established species, published after Riina et al. (2013), have not yet been framed within a global phylogenetic context. Regarding the spatial evolution of subg. *Esula*, Geltman (2015) provided an exhaustive narrative biogeographic account, but a dated phylogeny and a formal analysis to understand the biogeographic history of the subgenus are still lacking.

Our study aims to infer the spatio-temporal evolution and diversification history of *Euphorbia* subg. *Esula* through the analysis of a comprehensively species-level phylogeny. Specifically, we aim to test whether the remarkable species richness and widespread distribution of the subgenus were influenced by past geological and climatic events in the Western Palearctic region, as well as by subsequent migration to other continental landmasses. We also explored whether these events of geographic expansion and rapid diversification were linked to transitions between the annual and the perennial life-history. To achieve these goals, (1) we built a species-rich phylogeny of subg. *Esula* based on the DNA dataset of Riina et al. (2013) but expanded to increase taxon sampling to 321 species (66% of the subgenus diversity). This allowed us (2) to update the subgenus systematics, by placing new, revised, and previously unclassified taxa in their respective sections. (3) We then estimated species divergence times using Bayesian relaxed clocks and used the inferred time tree to estimate the biogeographic history of subg. *Esula*. Finally, (4) we inferred rates of speciation and extinction over time using Bayesian methods and explored potential correlations with life-history strategies; in particular, we tested whether the shift between annual and perennial life-histories was associated with increased diversification rates and these, in turn, were linked to habitat shifts associated with changes in elevation.

## 2 Material and Methods

### 2.1 Taxonomic updates of subg. *Esula*, taxon sampling, and alignment

We reviewed all the new literature concerning subg. *Esula* since the last global taxonomic treatment (Riina et al., 2013) of the subgenus (e.g. Lazkov & Sennikov, 2019; Yu et al., 2022; see full reference list in Table S1). We searched for newly described species, new synonyms and combinations, and all the taxonomic changes to generate the species list of the entire subgenus and each of its sections.

Based on this survey, we compiled a large data set using the multi-copy nuclear marker ITS and the plastid *ndhF* gene, including sequences of 341 species of subg. *Esula* and related lineages (outgroups) downloaded from GenBank. Our sampling covered 20 of 21 sections of this subgenus, as we excluded the monotypic sect. *Szovitsiae* because it was hypothesised to be an intersectional hybrid due to incongruent positions in the ITS and plastid tree (Riina et al., 2013). In addition to 321 species (out of the 490) ascribed to subg. *Esula* (Riina et al., 2013), we have included 17 species from the other subgenera in *Euphorbia*: seven species from subg. *Athymalus* (Peirson et al., 2013), five species from subg. *Chamaesyce* (Yang et al., 2012), five species from subg. *Euphorbia* (Dorsey et al., 2013); and two species from the sister clade of *Euphorbia* formed by *Neoguillauminia cleopatra* (Baill.) Croizat and *Calycopeplus casuarinoides* L.S.Sm. (Horn et al., 2012). Voucher information and GenBank accession numbers are listed in Supplementary File 1. The sequences were aligned and adjustments to the alignments were made in Geneious Prime v.2021.2.2 (Biomatters Ltd, Auckland, New Zealand). For the manual adjustment we applied a similarity criterion (Simmons, 2004). Summary statistics for each alignment according to the maximum parsimony criterion were estimated in PAUP v.4.0b10 (Swofford, 2003).

### 2.2 Phylogenetic analyses

Phylogenetic relationships were inferred using MrBayes v.3.2.7 (Ronquist et al., 2012) hosted on the CIPRES Science Gateway (Miller et al., 2010). Substitution models for each individual genetic marker were selected with the Akaike information criterion in jModelTest v.2.1 (Guindon & Gascuel, 2003; Darriba et al., 2012); these were SYM+I+G for ITS and GTR+G for *ndhF*. Because MrBayes does not implement the complexity of models evaluated in jModelTest, we used the nearest more complex model, which was GTR+I+G for ITS and GTR+G for *ndhF*. Two analyses of four chains each were run for 10^7^ generations, sampling every 1,000 generations. Convergence was assessed with the potential scale reduction factor (1) and by monitoring the standard deviation of split frequencies (< 0.01). A 50% majority rule consensus tree was constructed after discarding the first 25% samples as burn-in. Clade posterior probability values were considered as “high” support if ≥ 95% (Alfaro et al., 2003). No incongruent clades receiving high support were found among the individual gene trees. Therefore, we concatenated the two markers ITS and *ndhF* into a combined nuclear/plastid data set, which was used for further analyses. This matrix was partitioned by region, allowing each marker to have their own substitution model (above) and with the overall substitution rate unlinked between partitions.

### 2.3 Divergence time estimation

Lineage divergence times were estimated in BEAST v.1.8.4 (Drummond et al., 2012b). We used the concatenated matrix, with the same substitution models per marker as above, and the uncorrelated lognormal relaxed clock as the molecular clock prior. We compared two alternative models for the tree-growth prior: a pure-birth model and a birth-death model with incomplete taxon sampling (Stadler, 2009). The marginal likelihood of each model was estimated using parallel path sampling (PS) and stepping-stone sampling (SS) power posteriors (Baele et al., 2013), with 100 steps. Bayes Factors (BF) comparisons of log marginal likelihoods were computed as BF01=2(logM1-logM0), with values greater than two representing positive evidence for M1 and greater than six indicating strong evidence (Kass & Raftery, 1995). For each analysis, two Markov chain Monte Carlo (MCMC) chains of 10^7^ generations were run, sampling every 1,000th generation. Tracer v.1.7.1 (Rambaut et al., 2018) was used to monitor mixing and ensure that all parameters reached Effective Sample Size (ESS) values ≥ 200, indicating stationarity. TreeAnnotator v.1.8.4 (Drummond et al., 2012b) and FigTree v.1.4.4 (tree.bio.ed.ac.uk/software/figtree) were used, respectively, to generate and visualise the maximum clade credibility (MCC) tree, with a burn-in of 25% total length.

Given the lack of anatomical detail in the known fossils attributed to *Euphorbia* (Friis & Crepet, 1987; Anderson et al., 2009), absolute divergence times were estimated using six secondary calibration points obtained from the latest dated phylogenetic tree of the genus (Horn et al., 2014). Their calibration strategy employed one fossil calibration point (*Hippomanoidea warmanensis* Crepet & Daghlian, 1982) belonging to the outgroup tribe Hippomaneae within subfamily Euphorbioideae, and two secondary calibration points within Euphorbioideae, the latter obtained from a fossil-calibrated time tree of order Malpighiales (Xi et al., 2012). As calibration prior we used a normal distribution, with mean and standard deviation encompassing the confidence interval, the 95% highest posterior density credibility interval (HPD), and the mean of the posterior values estimated by Horn et al. (2014). This approach has been used in previous studies to perform divergence time estimation of clades within the subgenus (Frajman & Schönswetter, 2017; Stojilkovič et al., 2022; Faltner et al., 2023; Heimer & Frajman, 2023). The root node, the divergence between *Euphorbia* and the clade comprising *Calycopeplus* and *Neoguillauminia*, was assigned a normal distribution with mean (M) = 47.81 Mya and standard deviation (SD) = 4.20, 95% HPD = 40.90–54.72 Mya. The second calibration point, the divergence between subg. *Esula* and the clade comprising the other three subgenera (*Athymalus*, *Chamaesyce*, and *Euphorbia*) was assigned a normal distribution with M = 44.69 Mya and SD = 2.90, 95% HPD = 40.70–48.69 Mya. The third calibration point, the divergence within subg. *Esula* of the clade comprising *Euphorbia* sections *Lagascae* and *Lathyris*, was assigned a normal distribution with M = 40.49 Mya, SD = 4.00, 95% HPD = 33.20–47.70 Mya. The fourth calibration point, the divergence of the clade comprising all sections excluding the clade comprising sects. *Lagascae* and *Lathyris*, was assigned a prior distribution with M = 36.47 Mya, SD = 3.90, 95% HPD = 30.06–42.89 Mya. The fifth calibration point, the divergence of the clade comprising sects. *Myrsiniteae* and *Pithyusa*, was assigned a prior distribution with M = 32.53 Mya, SD = 4.00, 95% HPD = 39.00–26.06 Mya. Finally, the sixth calibration point, the divergence between sects. *Helioscopia* and *Holophyllum*, was assigned a normal distribution with M = 23.41 Mya, SD = 4.70, 95% HPD = 15.69–31.15 Mya. Divergence times estimations based on secondary calibrations from a higher-level taxonomic study are naturally associated with a larger degree of uncertainty than those employing fossil calibrations from the study group. To assess the impact of calibration strategies in our estimates, we performed a sensitivity analysis in which we used different calibration priors while also exploring alternative molecular clock models. Specifically, we used secondary calibration estimates from Janssens et al. (2020)’s study, who employed different fossils than Horn et al. (2014) and Xi et al. (2012) to calibrate an angiosperm-wide phylogenetic tree. The effect of this new calibration strategy was explored with two additional dating methods: treePL (Smith & O’Meara, 2012) applies the penalized likelihood algorithm (Sanderson, 2002) and implements a smoothed autocorrelation molecular clock; RelTime (MEGA v.11, Tamura et al., 2021) implements a rapid relaxed-clock method (Tao et al., 2020). We used as secondary calibrations the 95% highest posterior density (HPD) credibility intervals estimated by Janssens et al. (2020) for the following nodes: (1) the divergence between the *Calycopeplus* + *Neoguillauminia* clade and *Euphorbia*, (2) the crown node of *Euphorbia*, and (3) the crown node of subgenus *Esula*. We ran analyses with three calibration points or, alternatively, using only two calibrations, 2 and 3, temporally closer to the study group *Esula*. Unlike BEAST, MEGA and treePL require the user to provide a fixed tree topology. We used two different trees for dating: *(i)* the MrBayes tree estimated from the concatenated dataset (MEGA and treePL); *(ii)* a new tree estimated in IQ-TREE2 v.2.2.3 (Minh et al., 2020) under the maximum likelihood framework, only for treePL. Maurin (2020) recommended using treePL in a maximum likelihood environment. We were also unable to estimate confidence intervals in treePL when the MrBayes tree was used.

### 2.4 Biogeographic analyses

We inferred ancestral ranges and rates of range evolution using the Dispersal–Extinction– Cladogenesis (DEC) model (Ree & Smith, 2008), implemented within a Bayesian framework in RevBayes v.1.1.1 (Höhna et al., 2016a; Landis et al., 2018). Area definition was based on several criteria: the current distribution patterns of extant subg. *Esula* species (GBIF, 2023; POWO, 2023; Riina & Berry, 2023); the paleogeographic history of the African, American, and Eurasian continents (Sanmartín et al., 2001); and an effort to maximise congruence in the selected unit biogeographic areas with respect to other plant studies (Meseguer et al., 2015; Jin et al., 2020). We delimited five areas (Fig. 1): Africa (AFR), Eastern Palearctic (EP), Macaronesian region (MAC), North America, Central America, and Caribbean region (NAM), and Western Palearctic (WP). Widespread ancestral areas were limited to combinations of three areas, which is the maximum range of extant taxa. We use the native distribution of the extant species, i.e. discarding potential invasive ranges or human introductions (Meseguer et al., 2013). Four species of *Euphorbia* are distributed outside our predefined biogeographic areas: one in Chile and three in the Indo-Pacific region (Fig. 1, Table S1). Of these, only *E. glauca* G.Forst. (New Zealand) was included in our phylogeny. Since the number of ancestral states in the DEC model increases exponentially with the number of areas (Landis et al., 2018), we pruned *E. glauca* from the tree to avoid adding a sixth area to the analysis. It is unlikely that including this single species would have a major effect on the inference of biogeographic rates and ancestral ranges.

The DEC analysis was run on the MCC tree with one tip per species, assigning each tip the species’ entire native range. We used the *drop.tip* function of the package *ape* (Paradis & Schliep, 2019) in R (R Development Core Team, 2017) to prune the additional tips and the outgroup taxa from the tree; *E. glauca* was also excluded (see above). To inform the inference of ancestral regions, we retained one tip for each of the other three *Euphorbia* subgenera, coding them with the entire subgeneric range. This approach allowed for the consideration of biogeographic evolution before the MRCA of subg. *Esula*. Widespread ancestral areas were limited to combinations of three areas, which is the maximum range of extant taxa. Two independent analyses were run using the default priors for 10,000 generations, sampling every 10th generation, and results were summarised in the MCC tree. The first analysis (M0) was conducted with the rate of the baseline range evolution parameter (‘rate_bg’) modelled as constant through time and using a simplex prior, assigning equal probability for all cladogenetic events. The second analysis (M1) was run with cladogenetic events modelled with a Dirichlet distribution prior, in which the probability of allopatry “a” was equal to 1 minus the probability of sympatry “s”. Additionally, the M1 analysis included a time-stratified dispersal model, in which the baseline rate was rescaled according to geological connectivity scenarios, which reflected the shifting probability of migration over time. See a detailed explanation in the Extended Biogeographic Methods in Supplementary File 2.

We summarised nodal ancestral states as marginal PP onto the maximum a posteriori (MAP) tree. We also estimated the number and timing of dispersal and extinction events along the internal and terminal branches using a heuristic approximation to stochastic character mapping that does not require a rejection sampling step (Freyman & Höhna, 2019). Events of biogeographic change were summarised on the MAP tree using 500 time slices. We visualised the results using R scripts and the R packages *phytools* (Revell, 2012) and *RevGadgets* v.1.0 (Tribble et al., 2022). Finally, marginal likelihood values were estimated for the two alternative biogeographic models, M0 and M1, and compared via BF based on PS and SS power posteriors, with 100 steps each. The scripts to run these analyses can be found in Appendix S1.

### 2.5 Diversification analyses

All diversification analyses were run on the MCC tree from the BEAST analysis, pruned to remove the outgroup species and retain the 321 species of subg. *Esula*. We first plotted the lineage-through-time (LTT) plot to visually inspect the accumulation of lineages over time, using the *ltt.plot* function in the R package *ape* v.5.5 (Paradis & Schliep, 2019); a random sample of 1,000 chronograms from the Bayesian MCMC posterior distribution was also plotted to reflect uncertainty in divergence time estimates in the MCC tree. Next, we investigated whether speciation and extinction rates have been constant over the evolutionary history of subg. *Esula*, or if the subgenus has undergone episodic rate shifts, during which the diversification rate (speciation minus extinction) and the relative extinction rate (ratio of extinction to speciation) increased or decreased, simultaneously, across all lineages. We used the Compound Poisson Process of Mass Extinction model (CoMET; May et al., 2016) implemented in the R package *TESS* (Höhna et al., 2016b) to estimate shifts in speciation and extinction rates at discrete points in time, using a reversible jump MCMC algorithm and dynamic BF comparisons. We also used CoMET to estimate the magnitude and timing of mass extinction events (MEE), defined as discrete points in time when a significant percentage of the standing diversity (95–70%) is removed across all lineages. We ran two MCMC chains of 1 million iterations with a sampling frequency of 100 and ensured a minimum ESS of 1,000. We set the sampling fraction (ρ) at present to 0.66, i.e. the ratio between the number of sampled species in the phylogeny and the total extant species diversity in subg. *Esula* (321/490). Two different strategies were used to account for incomplete taxon sampling: uniform (random sampling) and diversified (maximise phylogenetic diversity).

We further explored time-varying birth-death scenarios using the Horseshoe Markov random field (HSMRF) episodic birth-death (EBD) model implemented in RevBayes (Magee et al., 2020). The use of a horseshoe estimator for sparse signals as the prior distribution in this EBD model allows modelling large jumps between prefixed time intervals, separated by relatively constant rates within each time interval. Incomplete taxon sampling was modelled with a global sampling fraction (ρ = 0.66), as in the CoMET analysis. We used ten time intervals as a compromise between inferring detailed temporal patterns and overfitting (i.e. a low ratio between parameters and data). The scripts to run the CoMET and HSMRF-EBD analyses can be found in Appendix S2.

The aforementioned EBD models assume that diversification rates are equal across all lineages at any given point in time. To implement a birth-death process where diversification rates vary across branches, we analysed the pruned MCC tree under the lineage-specific birth-death (LSBDS) model implemented in RevBayes (Höhna et al., 2019). LSBDS assumes a multistate state-dependent, speciation-extinction model (Maddison et al., 2007), in which each state has its own speciation and extinction rates, and where shifts in states (and hence in diversification rates) are modelled dynamically. The MCMC chain was run for 15,000 generations, using the same ρ value as above and an exponential prior. We then obtained the branch-specific diversification rate estimates for each branch of the tree using the rejection-free stochastic rate mapping algorithm (Freyman & Höhna, 2019). Appendix S3 provides the script to run the LSBDS analysis. The LSBDS model supported the existence of branch-specific changes in speciation rates in subg. *Esula* (see Results below). To test whether the observed shifts could be causally linked to changes in life-history, we reanalysed our phylogeny under the Binary State-dependent Speciation and Extinction model (BiSSE; Maddison et al., 2007; FitzJohn, 2012), implemented in a Bayesian framework in RevBayes. We coded life-history as two states: annual (0) or perennial (1) for all species of subg. *Esula*, including those not represented in our analysis (Table S1). We note that the definition of annuals used in our study includes those classified as “biennials” in the literature. We ran a single MCMC chain with 15,000 generations, sampling every 100th generation, and summarised ancestral states and state-shifts along branches in the MAP tree using stochastic free-rejection sampling with 500 time slices.

BiSSE assumes that all heterogeneity in diversification rates among clades in the phylogeny can be attributed to transitions in the focal character. However, rate heterogeneity could also be explained by unobserved ‘hidden’ characters that covary with the observed character across the phylogeny. To correct for the potential high Type I error rate in BiSSE (Rabosky & Goldberg, 2015), Beaulieu & O’Meara (2016) introduced the Hidden-State Speciation and Extinction model (HiSSE). In HiSSE, there are two ‘hidden’ states (A, B) within each state of the focal (“observed”) character states (0, 1), and each composite ‘observed-hidden state’ (e.g. 0A, 1B) is modelled as having its own speciation and extinction rate. We ran the HiSSE analysis in RevBayes, using lognormal priors for speciation and extinction rates centred in the extant diversity and exponential priors for asymmetric transition rates in the focal character. All other settings were similar to the BiSSE analysis. Since the RevBayes implementation of HiSSE does not apply state-specific ρ values, we additionally ran HiSSE in the R package *hisse* (Beaulieu et al., 2021), which allows the use of state-dependent sampling fractions. We used the function f = c(0.67, 0.66) to incorporate the percentages of annual and perennial species relative to the total diversity. Finally, we implemented in RevBayes a lineage-specific, character-independent rate-variable diversification model (CID; Caetano et al., 2018). Specifically, we modelled speciation and extinction rates as different between the two hidden states A and B, but equal for the two states within the focal character, annual and perennial. All other settings were identical to those in the BiSSE analysis. We used BF comparisons in RevBayes to compare the BiSSE and HiSSE models against the CID model, considered here as the null model, i.e. no causal relationship between the evolution of a trait and diversification rates. The marginal likelihood value for each model was estimated through PS and SS sampling with 50 parallel power posterior analyses, each with a chain length of 500 generations. See Supplementary File 3 for further details on the diversification analyses. Appendix S4 provides the scripts to run the BiSSE, HiSSE, and CID analyses.

### 2.6 Association between elevation and life-history

We used phylogenetic comparative analysis to test whether species elevational distributions were significantly different between annual and perennial life-histories. Specifically, we statistically incorporated the phylogenetic load in the estimation of the error component of the correlation elevation/life-history using three methods: Phylogenetic Generalized Least Squares (PGLS) analysis, Pagel’s Binary Correlation Test, and Phylogenetic Logistic Regression, implemented in the R packages *caper* (Orme et al., 2013), *phytools* with the fitPagel function, and *phylolm* (Ho et al., 2016) with the phyloglm function. Life-history was coded as a categorical binary variable: annual, perennial. Elevation – the response variable – was coded as a continuous trait for PGLS, represented by the mean elevation value of each species’ elevation range as shown in Table S1. For the fitPagel and phyloglm functions, elevation was coded as a discrete variable with two states. The 321 species coded for this analysis were assigned to either the “low” (distributed below 1,500 m) or “high” (above 1,500 m) elevation category (Table S1). If a species is predominantly distributed in one of the two categories but its range slightly overlaps with another category (e.g. 100–1,600 m), we coded the species for the range accounting for most of the species’ elevation distribution, with a maximum difference of 100 m (e.g., “low” in the previous example). Our division into “low” and “high” was intended to resemble those used in other studies of alpine floras, such as Carnicero et al. (2022, defining high as “alpine” and “subalpine” zones). In several cases, a species’ elevation range spans the entire elevational gradient, from “low” to “high” elevations. These species were assigned alternatively to the “low” or “high” categories in two independent analyses to account for the effect of the “binary” categorization enforced in the Pagel Correlation Test and the Phylogenetic Logistic Regression analysis, i.e., in one analysis, these “mixed” taxa were coded as “low” and in the second analysis, as “high”. In *phylolm*, we also used the mean elevation of each species’ elevation range as a third alternative analysis for the “mixed” species. The elevational range of each species was based on current distribution patterns and was estimated using the information available in the taxonomic literature (e.g. Lebrun & Stork, 2006; Pahlevani, 2017b; Caković & Frajman, 2020; Skubic et al., 2023), online floras and databases (e.g. Flora of China, 2008; FNA, 2023; GBIF, 2023). The scripts to run these analyses can be found in Appendix S5.

## 3 Results

### 3.1 Phylogenetic relationships and taxonomic updates

Phylogenetic relationships inferred in our study based on the individual gene topologies are shown in Figs. S1, S2, as well as the concatenated matrix in Figs. 2G, 2H, S3. The ITS alignment consisted of 328 accessions, whereas the *ndhF* alignment had 237 accessions (one per species). Summary statistics for the ITS and *ndhF* matrices and concatenated dataset are listed in Table S2. The majority of the sections were moderately to strongly supported as monophyletic both in the ITS (PP ≥ 0.90, 17 out of 20 sections, not sects. *Exiguae*, *Guyonianae*, and *Lathyris*, Fig. S1) and the *ndhF* tree (PP ≥ 0.90, 18 out of 20 sections, not sects. *Exiguae* and *Herpetorrhizae*, Fig. S2). All sections were strongly supported as monophyletic in the concatenated ITS+*ndhF* tree (PP = 1), except sects. *Exiguae* (PP = 0.54), *Guyonianae* (PP = 0.85), and *Lathyris* (PP = 0.95) (Fig. S3). The overall topology of the concatenated tree (Fig. S3) was congruent with the previous phylogenetic hypothesis of the subgenus (Riina et al., 2013), with the main exception being the clade formed by sects. *Lagascae* and *Lathyris*, which was recovered as the sister to the rest of subg. *Esula* (PP = 0.95).

**Fig. 2.**
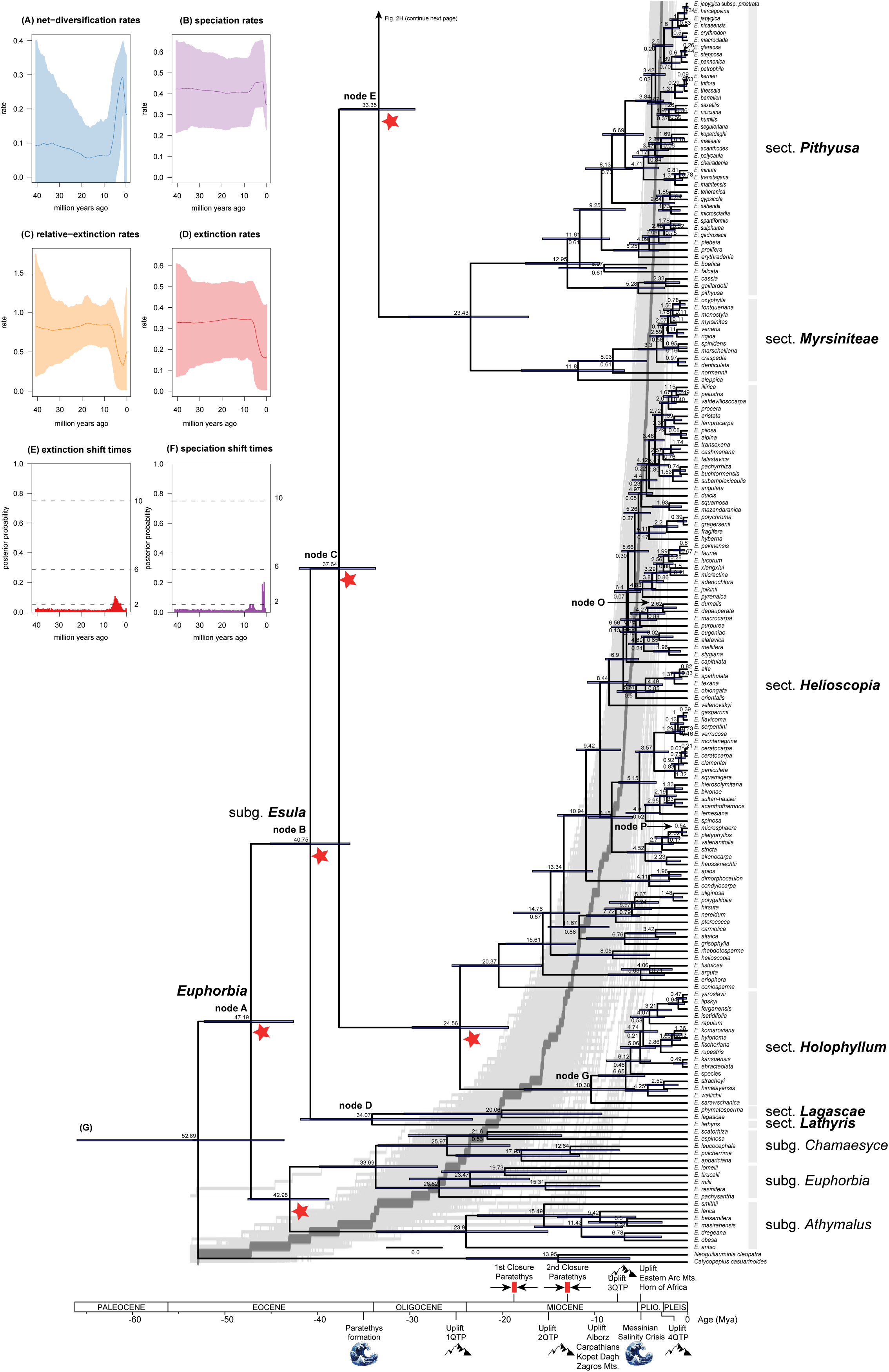
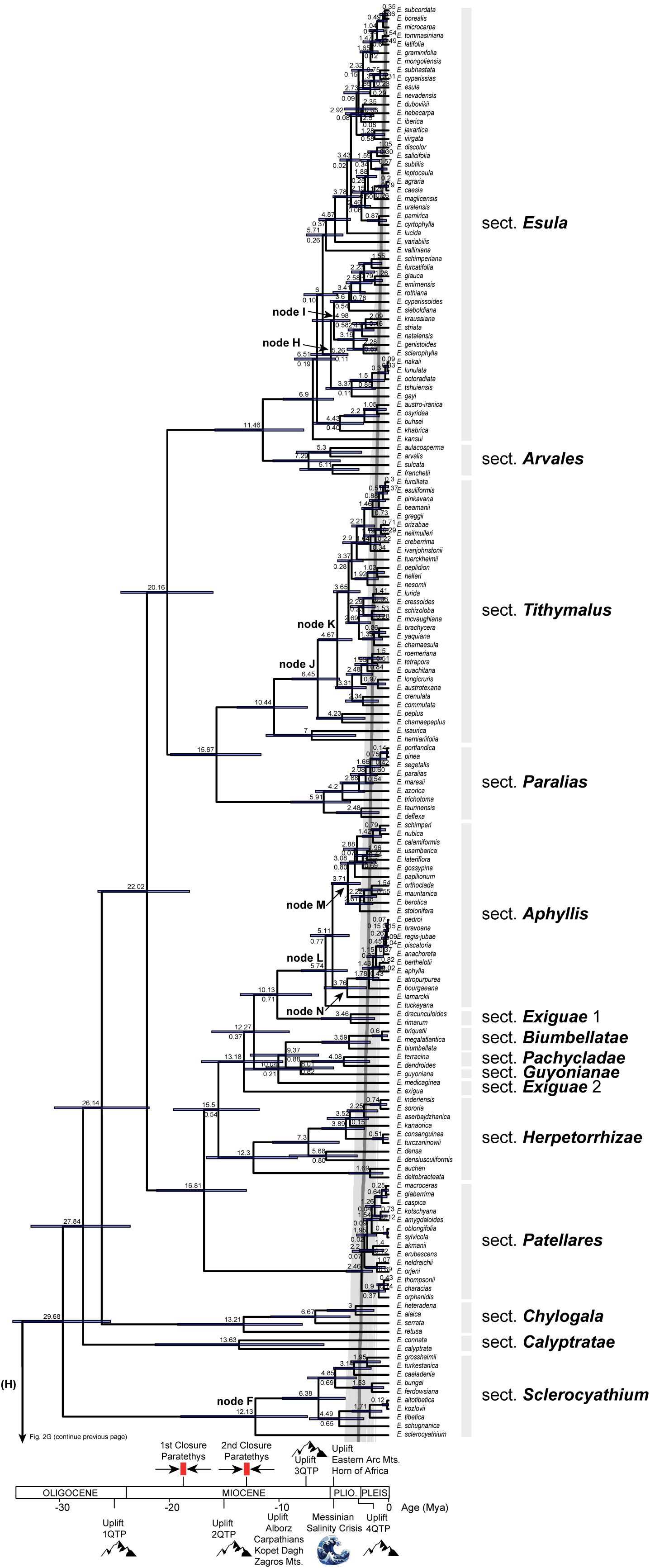
Lineage divergence times and diversification trajectories in *Euphorbia* subg. *Esula*. **(A–F)**: estimates of diversification rates over the evolutionary history of *E.* subg. *Esula* from the episodic birth-death model implemented in CoMET. Changes in **(A)** net diversification, **(B)** speciation, **(C)** relative extinction, **(D)** extinction rates, and **(E)** extinction and **(F)** speciation shift times; significance of the extinction and speciation shift times plots in (E) and (F) was assessed using Bayes Factor (BF) comparisons (BF > 2, positive support; Kass & Raftery, 1995). **(G–H)**: Maximum clade credibility (MCC) tree for *E.* subg. *Esula* and related taxa, showing phylogenetic relationships and lineage divergence times inferred by BEAST. The red stars indicate the position of six secondary calibrations points. Numbers above nodes indicate mean age estimates (Mya), those below nodes show posterior probability values < 0.90. Blue bars at nodes indicate the associated 95% highest posterior densities for all nodes. Lineage-through-time plots illustrate the cumulative number of lineages over time for *E*. subg. *Esula*. Dark grey curve corresponds to the combined MCC tree and light grey curves represent a random sample of 1,000 chronograms from the Bayesian MCMC posterior distribution to reflect uncertainty in divergence time estimates.

**Fig. 3.**
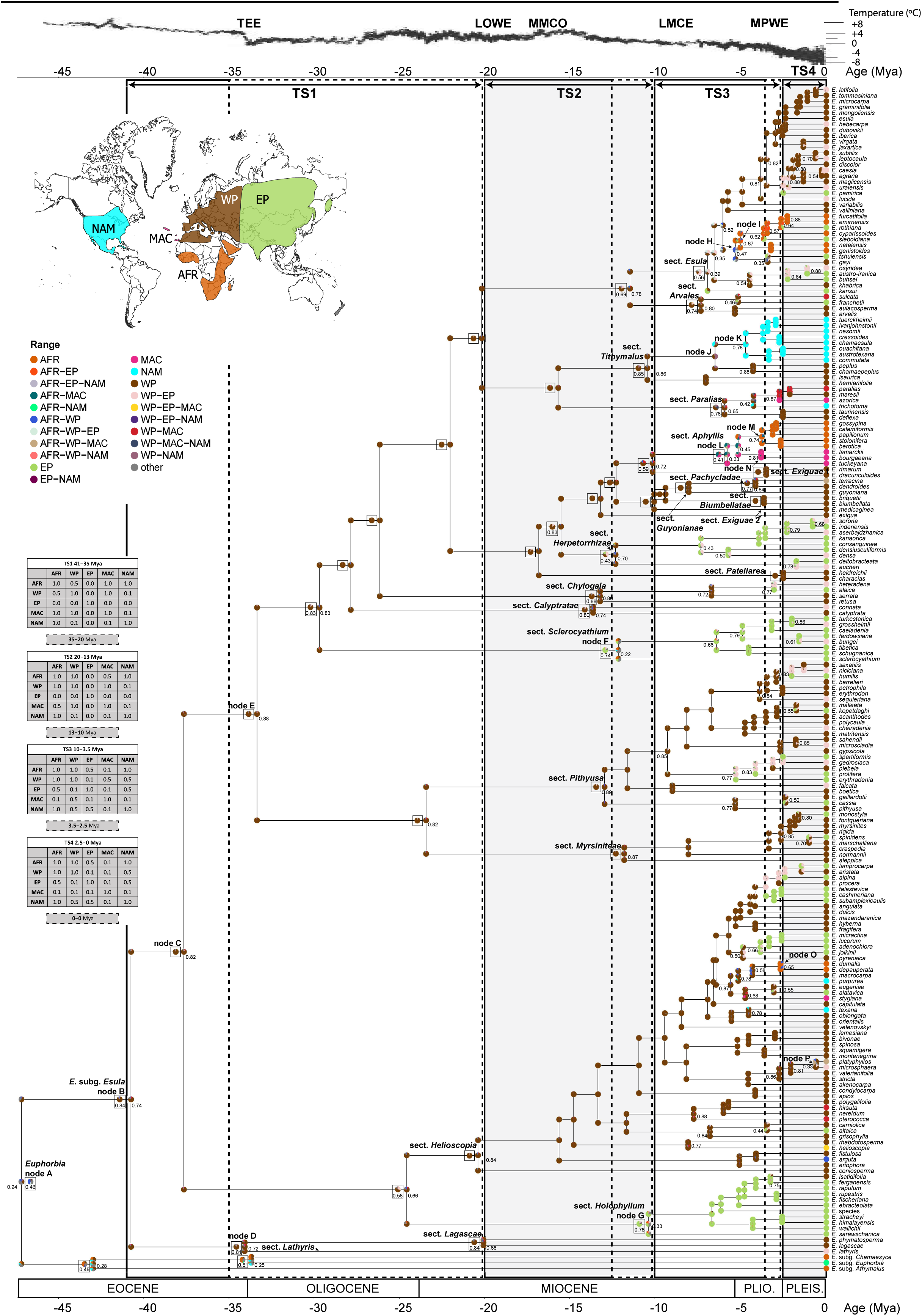
Reconstruction of the spatio-temporal evolution of *Euphorbia* subg. *Esula* and closest allies, using Bayesian dispersal extinction cladogenesis (DEC) and model M1 (see Methods). The phylogenetic tree is the maximum clade credibility (MCC) tree based on concatenated ITS and *ndhF* sequences, pruned from the BEAST tree shown in Figs. 2G, 2H. As representatives of the other three *Euphorbia* subgenera we included one taxon per subgenus, representing the main distributional area of the respective subgenus. To reduce the size of the tree, those clades within *E.* subg. *Esula* with ages less than 2.5 Mya that had the same distribution were eliminated. The maximum number of areas allowed at one time was three. A pie chart on the node indicates alternative ancestral geographical ranges, with pie slices indicating their posterior probabilities (PP). The reconstructions of the ancestral nodes of major diversification events of the M0 model are shown in black frames to the left of the M1 model reconstructions. Values below nodal pie charts indicate the PP for the ancestral range receiving the highest probability; only values < 0.90 are shown. Letters near nodal pie charts indicate the nodes mentioned in the text. Coloured nodes represent the geographical range state according to the legend. Colours of five major biogeographic areas correspond to the inset map. Coloured circles before the species names give present ranges. The smaller pie charts on the branches indicate the geographical range immediately after the speciation event; a pie on the node indicates how this range has changed as a result of range expansion and contraction events along the branch. Acronyms above the global mean temperature curve, obtained from Zachos et al. (2001), indicate climatic events which may have played a role in the evolution of *E*. subg. *Esula*: LMCE, Late Miocene Cooling Event; LOWE, Late Oligocene Warming Event; MMCO, Mid-Miocene climatic optimum; MPWE, Mid-Pliocene Warming Event; TEE, Terminal Eocene Event. TS1 to TS4 are time slices (TS) which divide the phylogenetic tree to capture important periods of tectonic rearrangements and/or climatic change (see Methods). The dashed lines within TS1 to TS3 symbolise the time interval in which each could have ended.

Our literature review revealed that subg. *Esula* comprises a total of 490 species: 457 species were listed in the taxonomic treatment of Riina et al. (2013); since 2013, nine names have been synonymized and 42 new species have been described. The phylogenetic hypothesis presented here allowed us to assign or confirm the assignation of these new species to the following sections: *Esula* (11 spp.), *Pithyusa* (11 spp.), *Helioscopia* (5 spp.), *Tithymalus* (5 spp.), *Holophyllum* (4 spp.), *Patellares* (3 spp.), *Paralias* (2 spp.), and *Sclerocyathium* (1 sp.) (Table S1). An updated list of all the currently accepted species names of subg. *Esula* by section is included in Table S1.

### 3.2 Divergence time estimation

BF comparison of the marginal likelihoods of the pure-birth and birth-death with incomplete sampling models in BEAST favoured the latter (PS = –40220.7990, Table 2A). The MCC chronogram (Figs. 2G, 2H) and the MrBayes tree based on the concatenated dataset (Fig. S3) showed very similar topologies and strong support values (PP ≥ 0.92) for all backbone nodes. Section *Exiguae* was not recovered as monophyletic in the BEAST tree, but as three separate clades in different positions (Fig. 2H): *Exiguae* 1 (*E. dracunculoides* Lam. and *E*. *rimarum* Coss. & Balansa.) sister to sect. *Aphyllis*, and *Exiguae* 2 as two separated branches (*E*. *exigua* L. and *E*. *medicaginea* Boiss.). This is compatible with the MrBayes topology (Fig. S3), where species of sect. *Exiguae* were grouped in a poorly supported clade (PP = 0.54, Fig. S3).

**Table 2.** Marginal likelihood values and Bayes Factor (BF) comparison for the different models applied to *Euphorbia* subg. *Esula*. **(A)** Marginal likelihood values for birth-death tree priors (pure-birth and birth-death with incomplete sampling) and uncorrelated relaxed molecular clock models, as estimated by path sampling (PS) and stepping-stone sampling (SS) in BEAST. **(B)** BF comparison of marginal likelihood values of the M0 and M1 biogeographic models. **(C)** BF comparison of marginal likelihood estimates using the Horseshoe Markov random field (HSMRF) episodic birth-death (EBD) model. BiSSE, HiSSE, and CID models under the species-level fraction. Marginal likelihood estimates are analysed here using the global phylogeny of *E.* subg. *Esula*, and parallel power posterior analyses with PS and SS.

| MODELS |  | Marginal likelihood |  |
| --- | --- | --- | --- |
| (A) | DIVERGENCE |  |  |
|  | TIME | ss | ps |
|  | ESTIMATION |  |  |
|  | Pure-birth | −40249.2082 | −40235.2450 |
|  | Birth-death with<br>incomplete<br>sampling | −40229.7963 | −40220.7990 |
| (B) | BIOGEOGRAPHI |  |  |
|  | C | ss | ps |
|  | Simple M0 | −349.415 | −349.4396 |
|  | Stratified M1 | −406.3398 | −406.3651 |
| (C) | DIVERSIFICATIO | ss | ps |
|  | N |  |  |
|  | HSMRF EBD | −2121.421 | −2121.095 |
|  | BiSSE | −2199.79 | −2199.606 |
|  | HiSSE | −2185.854 | −2185.675 |
|  | CID | −2184.551 | −2184.456 |

Divergence of subg. *Esula* from the other subgenera (node A, Fig. 2G) was dated in the Mid-Eocene (47.19 Mya, 95% HPD = 52.24–42.56 Mya) with strong support (PP = 1), while the MRCA of the subgenus (node B) was dated in the Mid-Eocene (40.75 Mya, 95% HPD = 45.1–36.47 Mya, PP = 1). The age of node C, grouping the majority of sections in subg. *Esula* was dated in the Late Eocene (37.64 Mya, 95% HPD 41.95–33.69 Mya, PP = 1), whereas node D (the MRCA of sects. *Lagascae* and *Lathyris*) was dated at the end of the Eocene epoch, close to the TEE: 34.07 Mya (95% HPD 41.08–23.21 Mya, PP = 0.94). The majority of subg. *Esula* sections (15 out of 20) are inferred to have originated during the Miocene, with MRCA ages ranging between 21.7 and 6.85 Mya. These estimates generally received high support (PP ≥ 0.99, Figs. 2G, 2H, Table S3). The exceptions were sects. *Biumbellatae*, *Pachycladae*, and *Patellares*, and clade *Exiguae* 1 within sect. *Exiguae*, whose MRCAs are dated around the Early-Mid Pliocene (4.76–3.14 Mya, PP = 1, Fig. 2H, Table S3). Credibility intervals around these age estimates were relatively small, except for the divergence of node D and the crown nodes of sects. *Calyptratae*, *Chylogala*, *Lagascae*, and *Sclerocyathium* (Figs. 2G, 2H). Detailed results from the divergence time analysis are shown in Table S3; the BEAST MCC tree is provided in Supplementary File 4.

The divergence time estimation sensitivity analysis, exploring alternative calibration, clock model and tree topology strategies, were highly congruent with the BEAST results, with mean ages differing in a few million years and with overlapping 95% confidence intervals (Table S4). These results are also congruent with those obtained by Horn et al. (2014) for *Euphorbia*, and Janssens et al. (2020) at the whole-angiosperm level, and closely align with divergence age estimates reported in other large-scale angiosperm studies (Ramírez-Barahona et al., 2020; Zuntini et al., 2024) (Table S4). This consistency across different calibration points and dating methods provides strong support for the robustness of our conclusions and indicates that they do not depend on a single dating approach.

### 3.3 Biogeographic inference

BF comparison of the marginal likelihoods of the M1 and M0 models favoured the M0 model (BF = 113.84, Table 2B). However, reconstructed nodal ancestral states and stochastic events of extinction and dispersal along branches were very similar in the two reconstructions. Fig. 2 depicts the M1 reconstruction to showcase the temporal geological stratification, but the M0 reconstruction is shown also for the backbone nodes. Fig. S4A shows the M0 reconstruction for all nodes in the tree. Marginal probabilities for the transition events of M0 and M1 models are shown in Figs. S4B and S5A, respectively. In the M1 model (Fig. 3), the ancestor of genus *Euphorbia* (node A) was reconstructed as having originated in the WP region, though support was low (PP = 0.24); other possible scenarios included AFR and NAM (Table S5). Model M0 reconstructed the ancestral range of node A as widespread in AFR–WP (PP = 0.46), with the next likely scenario adding EP with low support (AFR–EP–WP, PP = 0.10). In both M0 and M1 models, the origin of subg. *Esula* was inferred to be in the WP with high support (node B, Fig. 3, PP = 0.84, 0.74, Table S5). WP was also the ancestral range of many of the backbone nodes between 41 and 12 Mya (nodes C to G, Fig. 3), as well as the MRCA of many sections (e.g. *Lagascae*, *Helioscopia*, *Pithyusa*, *Myrsiniteae*, etc.), with moderate support (PP ≥ 0.64, Table S5). Only a few sections were inferred as originating outside WP (Table S5). Section *Aphyllis* (node L) originated in AFR−MAC in the M1 (PP = 0.33) and the M0 (PP = 0.41) model. This widespread range splitted into an AFR clade (node M, PP = 0.74) and a MAC clade (node N, PP = 0.81). The ancestral range of sect. *Holophyllum* (node G) was reconstructed as EP with high probability in M0 (PP = 0.78), but not in M1 (PP = 0.33). The MRCA of sect. *Sclerocyathium* (node F) was inferred as being ancestral to AFR in the M1 model (PP = 0.22) and to EP in the M0 model (PP = 0.74). Section *Herpetorrhizae* was reconstructed as WP (PP = 0.70) in M1 but EP−WP in M0 (PP = 0.43). The ancestral area of sect. *Esula* was inferred as widespread in EP−WP in both M1 (PP= 0.39) and M0 (PP = 0.56). At node H, AFR−WP was recovered by M1 (PP = 0.47) and EP−WP (PP = 0.31) by M0 (Fig. S4A). Node I was reconstructed as AFR in M1 (PP = 0.67) but AFR−EP (PP = 0.46) in M0 (Fig. S4A).

### 3.4 Diversification rates through time and in relation to life-history

The EBD models implemented in RevBayes and TESS supported a diversification trajectory with a slow initial accumulation in the number of lineages followed by an increase in the diversification rate from the Mid-Late Miocene to the present (apparent also in the LTT plot, Figs. 2G, 2H). The HSRMF model inferred a nearly flat net diversification rate up to ∼9−8 Mya, when it accelerates towards the present (Fig. S6). The CoMET model recovered a decrease in the net diversification rate between ∼35 Mya (r ∼0.10) and ∼16–14 Mya (r ∼0.07), followed by a slow increase that rapidly accelerated from 9 Mya onwards (r ∼0.3; Figs. 2A, 2C). Uncertainty in parameter estimation was high, i.e. 95% HPD credibility intervals were broad. Relative extinction rates showed the opposite pattern: an increase in the relative extinction rate between ∼30−9 Mya, followed by a plateau and a rapid decrease from 9 Mya towards the present (Fig. 2C); the latter is also captured by the HSRMF model (Fig. S6). BF comparisons in CoMET identified a downward shift in the extinction rate at ∼5 Mya (BF > 2, Fig. 2E). The upward shift in the speciation rate at ∼9–7 Mya is not significant (BF ∼ 2, Fig. 2F), but the increase at 2 Mya is strongly supported (BF > 4, Fig. 2F). CoMET detected several potential MEE between ∼40–12 Mya under the uniform sampling strategy (Fig. S7), but none were significant (BF < 2). No MEE were inferred under the diversified sampling strategy, which otherwise supported similar changes in the diversification trajectory (Fig. S8). Fig. S9 shows MCMC diagnostics in CoMET for some parameters under the uniform strategy. The LSBDS model in RevBayes supported differences in net diversification rates across lineages, with higher speciation rates in sect. *Patellares* and in several clades within sects. *Aphyllis, Esula, Helioscopia, Holophyllum, Myrsiniteae, Paralias, Pithyusa*, and *Tithymalus* (Fig. 4). The extinction rate was estimated as 0.0001 events per lineage/ million years (Fig. S10). All SSE analyses reached convergence, with trace plots showing adequate mixing and ESS values larger than 200 for all parameters (for many, > 1,000). BF comparisons based on PS and SS power posteriors supported HiSSE as the best model (PS = –2,185.675), though only the comparison with BiSSE received strong support (BF > 13, Table 2C). BiSSE recovered significant differences in the speciation rate and the net diversification rate between the two life-histories: these were higher for the perennial than for the annual state (Fig. 4); the relative extinction and the extinction rates did not differ between states (Fig. S11, Table 3). The rate of transition from annual to perennial was higher than in the other direction (Fig. S11, Table 3). The ancestral condition for the MRCA of subg. *Esula* was inferred to be annual, while the first nodes recovered as perennial date back to the last 13 million years (Fig. 4). Marginal probabilities for the MRCAs of sections were inferred with strong support (PP > 0.90), with the exception of sects. *Calyptratae*, *Chylogala*, *Exiguae*, *Paralias*, *Pithyusa*, and *Sclerocyathium*; within-section ancestral states were more uncertain (Fig. 4). Stochastic character mapping (Fig. S12A) recovered as many as 22 independent transition events from annual to perennial, starting ∼13–12 Mya and extending as late as ∼2.5 Mya (0.051 events Mya^-1^). Six reversal events to annual were also inferred, beginning at ∼6 Mya (0.032 events Mya^-1^, Fig. S12A). Marginal probabilities for these transition events are shown in Fig. S12B.

**Fig. 4.**
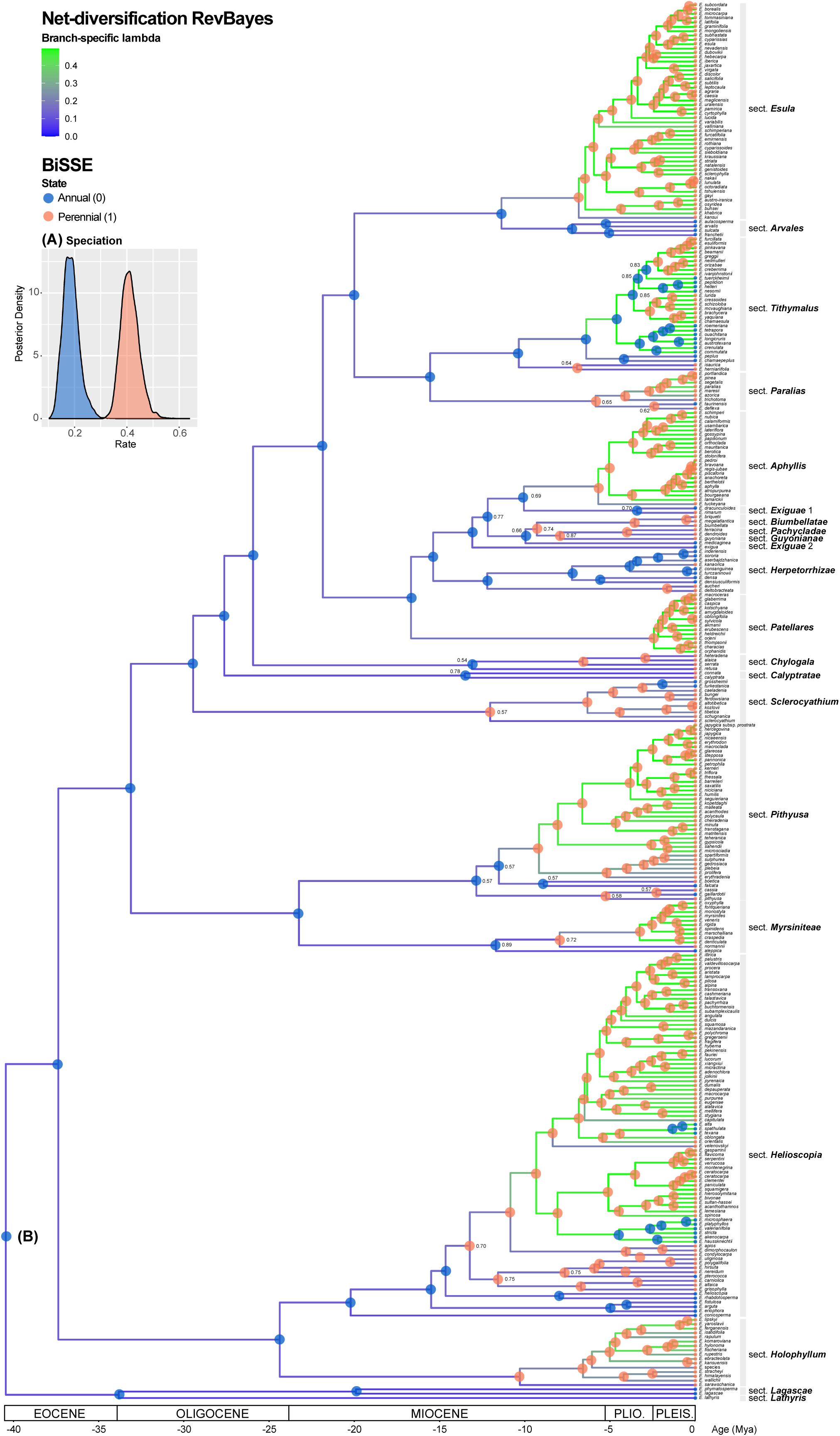
Diversification and life-history reconstruction within Euphorbia subg. Esula inferred with RevBayes. (A) Posterior distribution of speciation rates for annual (blue) and perennial (orange) lineages with BiSSE model. (B) Estimated branch-specific speciation rates in the pruned maximum clade credibility tree with the LBDS model; circles at nodes are marginal probabilities for ancestral character states inferred with BiSSE: blue, annual; orange, perennial. Branch colours present marginal stochastic character mapping shown in the heat-map diagram in the upper left corner.

**Table 3.**
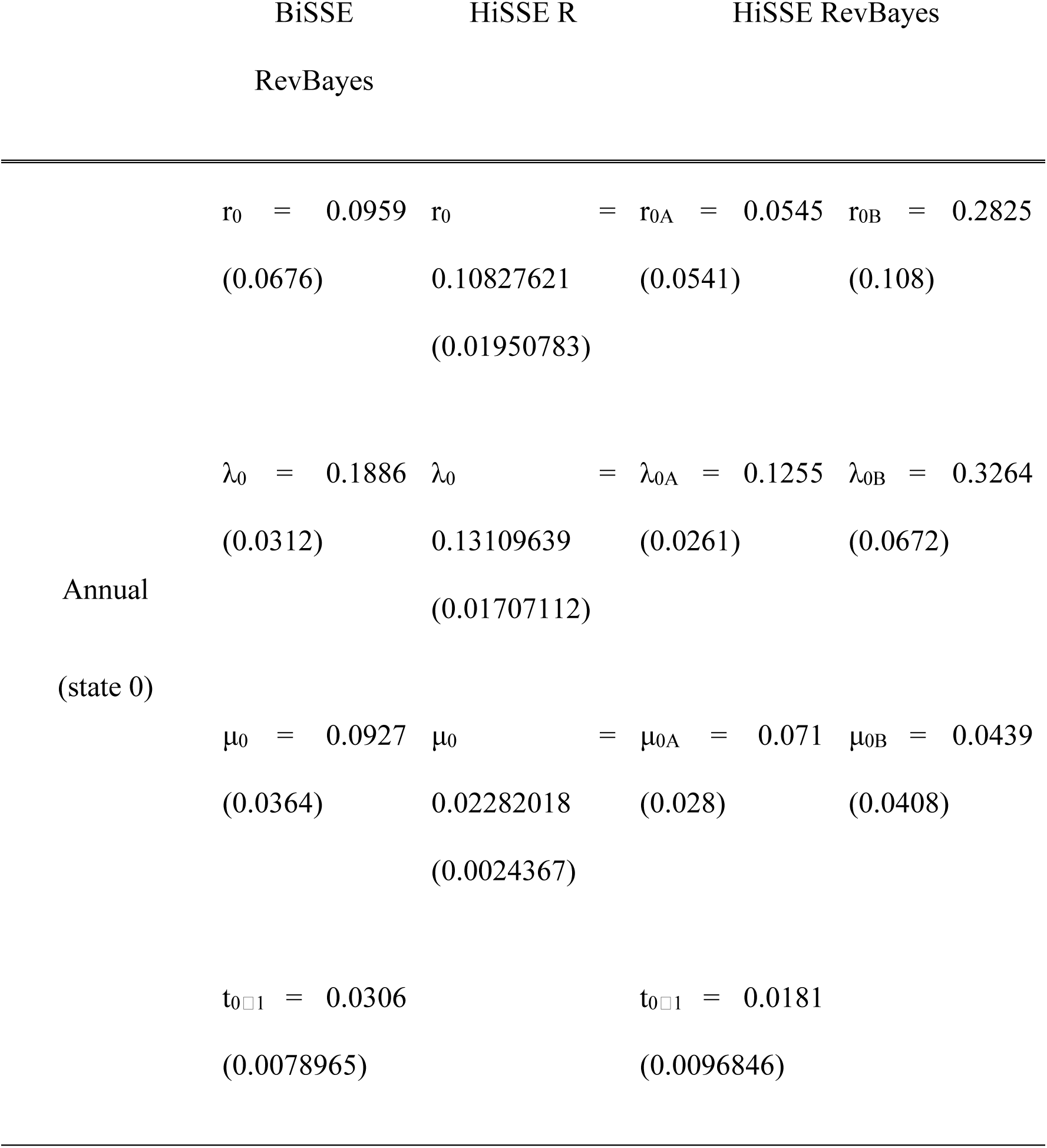
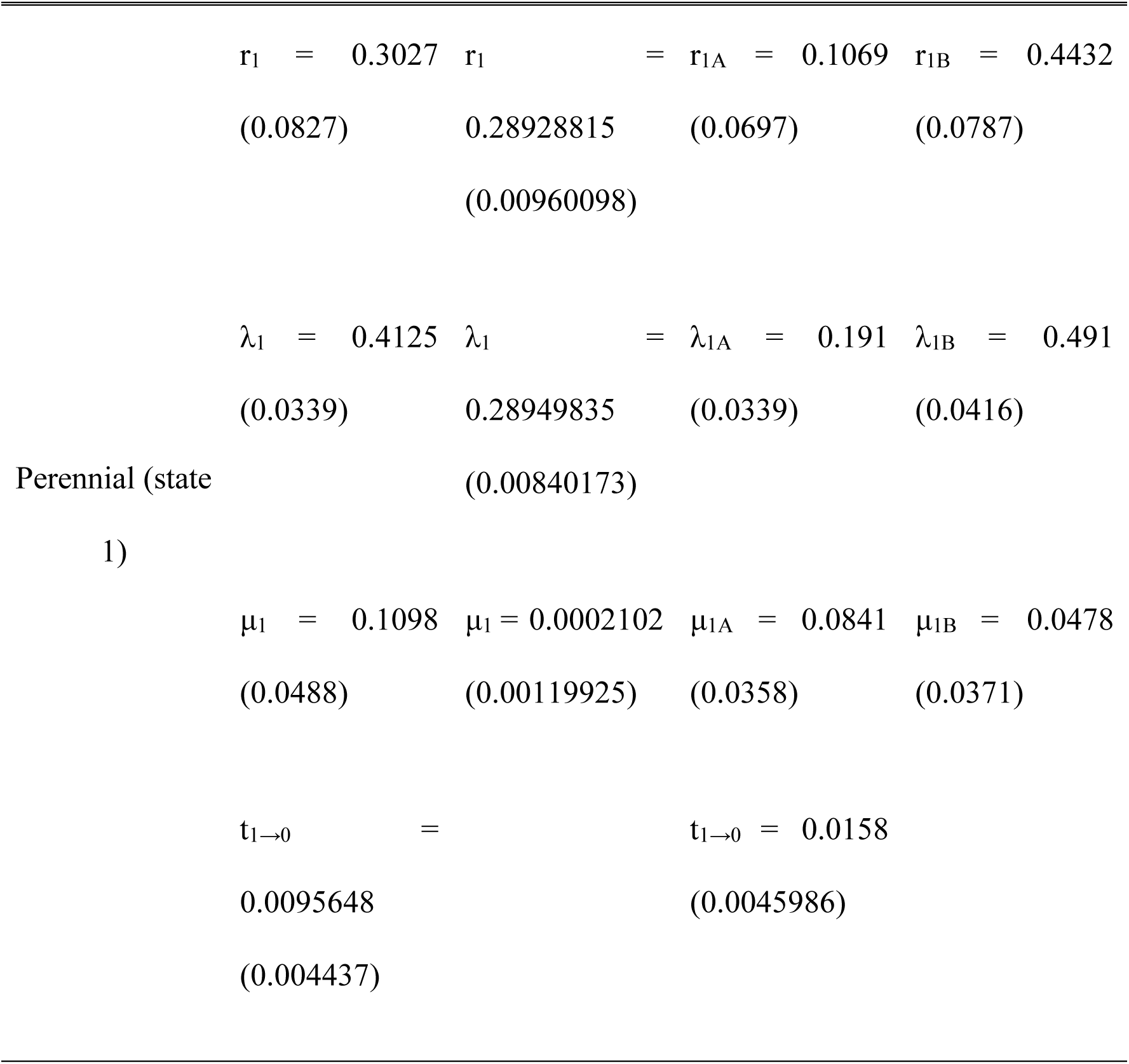
Diversification parameter estimates of *Euphorbia* subg. *Esula* (*r*, net diversification rate; *λ*, speciation rate; *μ*, extinction rate; *t*, transition rate; 0, annual; 1, perennial) obtained in RevBayes using the R packages *diversitree* and *hisse*. Reported values are means and standard deviations (in brackets) across tips and nodes obtained after model averaging.

HiSSE results were somewhat contradictory (Figs. S13–S17). RevBayes HiSSE (Fig. S13) inferred higher net diversification and speciation rates for the perennial state compared to the annual state, irrespective of the hidden state, i.e. significant differences in rate values were found between states 0A and 1A, and between 0B and 1B (Fig. S14, Table 3). Similar to BiSSE, no significant differences were observed for the relative extinction and extinction rates (Fig. S14), but unlike BiSSE, no significant differences were found between the two types of transition rates (Fig. S14, Table 3). However, the most important difference with BiSSE lies in the inference of ancestral life-histories (Fig. S15). The MRCA of subg. *Esula* was inferred to be a perennial plant (1A), with moderated probability (PP = 0.57, Fig. S15A). Stochastic character mapping recovered 18 transitions along the phylogeny from the perennial state to the annual state, mostly associated with hidden state A: 15 events from 1A to 0A and one event from 1A to 0B (Fig. S15B); conversely, only two events from 1B to 0B were inferred (within sect. *Helioscopia*, Fig. S14B). Transitions from annual to perennial were less frequent: three events from 0B to 1B within sects. *Herpetorrhizae* and *Tithymalus*. For the focal trait, most transitions occurred within the perennial state: eight events from 1A to 1B and only one transition from the annual state 0A to 0B (within sect. *Herpetorrhizae*, Figs. S14B, S15A). Marginal probabilities for these transition events are shown in Fig. S15C. There was a strong impact of the hidden state in the type and frequency of transition events: 0A and 1B were the least frequent source and the most frequent sink states, respectively (Figs. S14B, S15A). This can also be observed in the speciation rates (Fig. S13). Differences between hidden states corresponding to the same life-history (0A vs. 0B and 1A vs. 1B) were larger than differences between different life-histories associated with the same hidden state (0A vs. 1A, or 0B vs. 1B, Fig. S14). The perennial state with hidden state B (1B) was associated with the highest speciation and net diversification rates, while the opposite pattern is observed for the annual state with hidden state A (0A, Fig. S13). Unlike RevBayes, R HiSSE recovered results that were more similar to those from BiSSE (Fig. S16). The ancestral state of the MRCA of subg. *Esula* and many of the earliest speciation events were inferred to be annual plants, and up to 21 events of transition to perenniality and six reversals from perennial to annual life-history were inferred both at the sectional and within-section level (Fig. S16), in agreement with the BiSSE results (Figs. S11, S12A). Speciation and net diversification rates were higher for the perennial than for the annual state (Fig. S16), whereas the opposite pattern was found for the extinction and relative extinction rates (Fig. S17).

### 3.5 Association between life-history and elevation

Of the 321 species from subg. *Esula* analysed, 52 were annual (16%) and 269 perennial (84%) (Fig. S18, Table S1). Of them, 33 annual species were coded for low elevation, one for high elevation, and 18 species spanning the two ranges (“mixed”). Perennial species were distributed as follows: 117 at low elevation, 57 at high elevation, and 95 in both categories (Fig. S18, Table S1). All phylogenetic correlation analyses (fitPagel, PGLS, and phyloglm) supported the existence of a significant positive correlation (p < 0.05, Table 4) between life-history and elevation. Though only 2% of total variance was explained by life-history in the continuous PGLS analysis (R^2^, p = 0.011), the categorization analyses fitPagel and phyloglm support a significant correlation (p < 0.001, Table 4), between the high elevation type and the perennial life-history under all alternative codification schemes.

**Table 4.** Results of the phylogenetic comparative analysis using three methods: Phylogenetic Generalized Least Squares (PGLS) analysis, Pagel’s Binary Correlation Test with the fitPagel function, and Phylogenetic logistic regression with the phyloglm function.

| Analysis | Elevation<br>value | Value | p-value | Notes |
| --- | --- | --- | --- | --- |
| PGLS | Mean | $2.773 \times 10^{-5}$ | 0.011 | $R^2$: 0.017, Lambda: 1 |
| fitPagel | Mixed in low | 24.242 | $7.144 \times 10^{-5}$ | Method: fitDiscrete |
| fitPagel | Mixed in high | 12.070 | 0.001 | Method: fitDiscrete |
| phyloglm | Mean | 0.001 | 0.001 | Method: logistic_IG10, alpha:<br>0.044 |
| phyloglm | Mixed in low | 1.845 | $2.012 \times 10^{-5}$ | Method: logistic_IG10, alpha:<br>0.021 |
| phyloglm | Mixed in high | 0.480 | $1.087 \times 10^{-5}$ | Method: logistic_IG10, alpha:<br>0.039 |

## 4 Discussion

### 4.1 An updated phylogeny and a time tree for subg. *Esula*

In this study, we built a new, comprehensively phylogenetic tree for subg. *Esula*, representing 66% of the subgenus’s extant diversity: 321 species, including 60 species not present in the phylogenetic analysis of Riina et al. (2013), and covering all major geographic regions of species occurrence. No phylogenetic hypothesis of genus *Euphorbia* using large scale genomic data has been published yet, though there have been attempts to infer a genome-level phylogeny for less inclusive clades (Villaverde et al., 2018). Here, we chose to follow other studies focused on mega-diverse Holarctic lineages (Mansion et al., 2012; Betancur et al., 2019), which favoured taxon sampling over genetic sampling: our reduced two-gene phylogeny represents a 10% and 57% increase in taxon sampling compared to previous studies of subg. *Esula*, Riina et al. (2013) and Horn et al. (2014), respectively. Our backbone phylogeny for the subgenus is also identical to that presented in Horn et al. (2012) based on 10 genetic regions and focused on genus *Euphorbia*. A pattern of overall congruence between phylogenies based on Sanger-sequencing and those inferred using high-throughput sequencing has been found in other plant groups, including the persistence of recalcitrant regions of hard polytomies (Herrando-Moraira et al., 2019; Villaverde et al., 2020; Gallego-Narbón et al., 2022; Fonseca et al., 2023).

The expanded phylogeny presented here confirms the monophyly of subg. *Esula* and is consistent with major phylogenetic relationships recovered in previous studies, as well as the relationships of subg. *Esula* with its sister subgenera (Frajman & Schönswetter, 2011; Horn et al., 2012, 2014; Riina et al., 2013). The main exceptions are the lineage formed by sects. *Lagascae* and *Lathyris* (node D), which is recovered here as sister to the rest of subg. *Esula* (PP = 0.95, Fig. S3), and sect. *Exiguae*, which is here sister to the clade formed by sects. *Pachycladae* to *Guyonianae* (PP = 0.75, Fig. S3). Future phylogenomic studies in subg. *Esula* should address these incongruencies by ensuring that representatives of these conflicting clades are included. Our densely sampled phylogeny of subg. *Esula* also allowed us to update the current classification by placing recently described species in their respective sections (Table S1), e.g. *E. mongoliensis* M.H.Li & C.H.Zhang (Zang et al., 2021). It also enables the systematic placement of previously misclassified taxa that were assigned to sections based on morphology alone. For example, *E. hylonoma* Hand.-Mazz. was classified in sect. *Helioscopia* by Riina et al. (2013), but our analyses recovered this species within sect. *Holophyllum*. This phylogenetic position was first shown in Zang et al. (2021) and it is congruent with the geographic distribution of *E*. *hylonoma* (China), as Eastern Asia is an important centre of diversity for sect. *Holophyllum*. Besides the geographic evidence, a close examination of the type specimen of *E. hylonoma* reveals that the ovaries are smooth; capsules are described as smooth in Flora of China (2008). Smooth capsules are typical of sect. *Holophyllum*, whereas most of the species in sect. *Helioscopia* have verrucose capsules. A similar case is *E. normannii* Schmalh. ex Lipsky, which was placed in sect. *Arvales* by Riina et al. (2013). Our phylogenetic results placed this species within sect. *Myrsiniteae* (Figs. 2G, S1, S3), in agreement with Geltman (2015) and the nuclear phylogeny of Frajman & Geltman (2021). However, Frajman & Geltman (2021) assigned *E. normannii* to sect. *Pithyusa* based on the plastid phylogeny and morphology (similar to *E*. *falcata* L.). Lastly, as demonstrated by Faltner et al. (2023), *E. orphanidis* Boiss., previously classified in sect. *Pithyusa* (Riina et al., 2013), is now part of sect. *Patellares*.

### 4.2 A Madrean-Tethyan clade with boreotropical ancestors and continental-arid adapted descendants

Our spatio-temporal reconstructions place the stem ancestors of subg. *Esula* in the Early Eocene (node A, 47 Mya, Fig. 2G, Table S4), a period characterised by globally warm temperatures, during which a mixed evergreen-deciduous boreotropical forest reached the higher latitudes of the Holarctic (Morley, 2003). The first diversification event in the subgenus (node B, 41 Mya, Fig. 2G, Table S4) predates the global climate cooling brought about by the TEE (Fig. 3). This and the inferred WP ancestral distribution (Fig. 3) lend support to the hypothesis that the crown ancestors of subg. *Esula* formed part of the Madrean-Tethyan vegetation that covered the Tethyan shores during the Late Paleogene (Table 1: GI1; Axelrod, 1975). The TEE, a drop of 10°C in global temperatures, favoured the replacement of the warm-adapted boreotropical forest by a temperate, cold-adapted mixed-mesophytic vegetation (Tiffney, 1985a; Zachos et al., 2001). The cooling TEE has been associated with high extinction rates and migration to southern latitudes in angiosperms, based on evidence from the fossil record and molecular phylogenetic studies (Morley, 2003; Plana et al., 2004; Antonelli & Sanmartín, 2011). It is likely that the Paleogene boreotropical and Tethyan ancestors of subg. *Esula* underwent extinction during this period of early diversification. CoMET shows a drop in the net diversification rate and a concomitant increase in the relative extinction rate, that started at ∼35 Mya and lasted until the Mid-Miocene (Figs. 2A, 2C, S7, S8). The HSMRF model recovers a flat, nearly constant accumulation of lineages, congruent with high turnover rates, from the Late Eocene until the Mid-Miocene, followed by a slight increase in the speciation rate (Fig. S6). Interestingly, the origin of species-poor sects. *Lagascae* and *Lathyris* is inferred around the TEE (Fig. 2G), suggesting that their extant low diversity is due to extinctions in their early evolutionary history. Diversification rates in subg. *Esula* started to slightly increase at ∼18–16 Mya, while relative extinction rates show a plateau (Figs. 2A, 2C, S7, S8); although less remarkable, the increase in diversification rate is also captured by the HSMRF model (Fig. S6). The Early-Mid-Miocene was a period of major tectonic and climatic changes: the collision of the Arabian and Eurasian plates closed the eastern arm of the Tethys Seaway (TTE) and led to the enclosure of the Paratethys Sea (Table 1: GI2; Liu et al., 2018). There was also a major increase in global temperatures (MMCO) and a drop in sea level (Shevenell et al., 2008; Zachos et al., 2008). Global temperatures plummeted at the start of the Late Miocene (11.6 Mya; Zachos et al., 2008). This dramatic cooling event (LMCE, Table 1: GI3) is captured in the EBD models as a notable surge in the diversification rate and a major decline in the relative extinction rate (Figs. 2A, 2C, S6, S7, S8). Though most sections in subg. *Esula* originated during the Early to Mid-Miocene, a large percentage of speciation events within each section are dated between 10 and 2.6 Mya (Figs. 2G, 2H), a period that coincides with the onset of a global climate cooling trend that climaxed with the Pleistocene glaciations.

Two conclusions can be drawn from these analyses. *(i)* There is not a direct correlation in subg. *Esula* between shifts in diversification rates and global changes in temperature: climatic events such as the LOWE (Table 1: GI2) are not captured by our EBD models, and both warming (MMCO) and cooling (LMCE) events triggered increases in the net diversification rate (Figs. 2A, S6, S7, S8). *(ii)* The pattern of species accumulation in subg. *Esula* seems, instead, to have responded to events of geographic evolution (Fig. 3). In the Early-Mid-Miocene, land bridges connected the Tethys and Paratethys shores, allowing dispersal across the proto-Mediterranean (Table 1: GI2; Meulenkamp & Sissingh, 2003; Manafzadeh et al., 2017). Subsequent rises in global sea level in the Mid-Miocene disrupted these connections, isolating proto-Mediterranean lineages on the eastern and western sides and are hypothesised to have triggered allopatric speciation via east-west geographic disjunctions (Sanmartín, 2003). Our biogeographic reconstruction shows that the MRCAs of the majority of sections in subg. *Esula* and many Early Neogene diversification events within each section (∼23–12 Mya, Figs. 2G, 2H) are confined to the WP region (Figs. 3, S4A, S4C, S5B). We thus hypothesise that tectonic rearrangements along the western Tethyan shores during the Early-Mid-Miocene caused the minor increase in diversification and speciation rates captured by our EBD models (Figs. 2A, 2B, S6, S7, S8). A similar explanation might account for the dramatic increase in net diversification and the concomitant decrease in relative extinction rates recovered by our EBD models for subg. *Esula*’s most recent past (Figs. 2A, 2C, S6, S7, S8). From the Late Miocene to the Mid-Pliocene many sections, which were originally confined to the WP, expanded their range towards other continental regions via dispersal events (Figs. 3, S4A). For example, several independent colonisations of the EP are reconstructed within sects. *Helioscopia*, *Herpetorrhizae*, and *Pithyusa* at ∼7–5 Mya (Figs. 3, S4A, S4C, S5B); range expansion from WP to EP in sect. *Lathyris* also occurred around this period (Fig. S5B) or slightly earlier, at ∼9 Mya (Fig. S4C). Expansion to the EP in sects. *Holophyllum* and *Sclerocyathium* is also dated in the Late Miocene (∼12.6–10.3 Mya, Figs. 3, S5B), though M0 places this event earlier (∼20 Mya, Fig. S4C). Migration from the WP to the EP might have been facilitated by the uplift of the QTP and the Anatolian and Iranian plateaus, which, though they started in the Late Oligocene-Early Miocene (Table 1: GI2), experienced major uplifts in the Mid-Late Miocene (Table 1: GI3–GI4). The elevation of high mountain plateaus in Central and Western Asia (e.g. the QTP, Alborz, Kopet Dagh, and Zagros mountain ranges) triggered widespread aridification across Eurasia (Barbolini et al., 2020). Plant lineages adapted to the colder and more arid continental climates migrated westward to the proto-Mediterranean region, displacing the temperate-adapted, mixed-mesophytic lineages (Manafzadeh et al., 2014, 2017; Meseguer et al., 2018; Shamsabad et al., 2021). Conversely, several proto-Mediterranean lineages of subg. *Esula* expanded their range into the newly formed mountains in western Asia, as discussed above. Adaptation to the drier, colder conditions in the eastern proto-Mediterranean region, caused by the Mid-Late Miocene closure of the Paratethys Sea and the Paratethys salinity crisis, might have aided this migration (Table 1: GI3; Manafzadeh et al., 2014). The uplift of the QTP3 and the MSC introduced an even colder and drier climate, with an almost complete desiccation of the Mediterranean Basin, and might be responsible for the migration events in sects. *Lathyris* and *Pithyusa* (Figs. 3, S5B). Range expansions into the EP in the Plio-Pleistocene (3.5–1.7 Mya, e.g. *E. aucheri* Boiss., Figs. 3, S5B) could have been mediated by the onset of the Mediterranean climate, the uplift of the QTP4, and the final closing of the Paratethys Sea (Table 1: GI4), whereas climatic oscillations in the Quaternary might explain the Late Pleistocene dispersal in *E. microsphaera* Boiss (Figs. 3, S5B). Our results thus support Takhtajan’s (1986) hypothesis that historical range expansion events, facilitated by the uplift of the QTP, explain the shared elements between the current Himalayan and Mediterranean floras. Uplift of the QTP3 and increasing continentalization in the proto-Mediterranean Basin due to the closure of the connection with the Atlantic Ocean (MSC, Table 1: GI4), might have also facilitated migration from the WP into AFR in the Pliocene. Events of geographic expansion into AFR and MAC are inferred in several sections – *Aphyllis* (node M), *Esula* (node I), and *Helioscopia –* most of them dated in the Early-Mid-Pliocene (Figs. 3, S4A, S4C, S5B). Similar events of range expansion from WP to AFR have been observed in other plant lineages with adaptive syndromes to strongly seasonal climates (Horn et al., 2014; Manafzadeh et al., 2014). Within sect. *Esula*, two dispersal events from AFR to EP are inferred along the stem ancestors of *E. sieboldiana* C.Morren & Decne. (∼3.6 Mya) and *E. rothiana* Spreng. (∼2.6 Mya) (Figs. 3, S5B). These events could have been driven by monsoon wind systems, occasional storms, or in floating vegetation, allowing seeds to travel between Africa, the Chagos archipelago, the Comores, India, and the Seychelles, as suggested in other studies (Malcomber, 2002; Renner, 2004; Li et al., 2009).

Finally, our temporal and biogeographic findings align with other paleobotanical (Grímsson & Denk, 2007; Grímsson et al., 2008) and molecular phylogenetic studies (Milne, 2004; Denk et al., 2010; Gorospe et al., 2020) that have inferred post-Miocene dispersal events from the WP to the Nearctic region across the Atlantic Ocean. Section *Tithymalus* originated from a single dispersal event from WP to NAM between ∼6.45–4.67 Mya, (Figs. 3, S4A, S4C, S5B). Three additional dispersals around the Mid-Pliocene to Late Pleistocene (∼3.5–1.7 Mya) were inferred in the stem ancestors of *E. trichotoma* Kunth (sect. *Paralias*), *E. purpurea* (Raf.) Fernald, and *E. texana* Boiss. (sect. *Helioscopia*) (Figs. 3, S5B). Parts of the Greenland-Scotland Transverse Ridge may have persisted above sea levels until the Pliocene (Thiede & Eldholm, 1983; Poore et al., 2006) and could explain these recent dispersals, which postdates the age of the trans-Atlantic land bridges (Tiffney, 1985b; Sanmartín et al., 2001). Entrance of subg. *Esula* into the New World coincides with the establishment of the Mediterranean climate in southwestern North America (Millar, 2012; Rundel et al., 2016), so these migration events could also be explained by long-distance dispersal from the Mediterranean region to the Madrean region of California in more recent times (Vargas et al., 2013; Gorospe et al., 2020). Interestingly, no events of dispersal from the EP to NAM region were inferred within subg. *Esula*. This stands in contrast with the pattern observed in other plant and animal lineages, for which the Bering Land Bridge (Late Cretaceous to Mid-Pliocene) acted as a route of biotic exchange between Eastern Asia and North America (Sanmartín et al., 2001; Li et al., 2015; Wen et al., 2016; Landis et al., 2021; Rose et al., 2023).

### 4.3 Mountain colonisation and rapid diversification preceded or were concomitant with changes in life-history

Although heterogeneously distributed, annuality has emerged repeatedly along the evolutionary history of angiosperms, including more than 100 families and 30 orders, highlighting its importance as a reproductive trait in this plant lineage (Angiosperm Phylogeny Group, 2016). The percentage of annual plants in angiosperms has been estimated to be 6–13% (Angiosperm Phylogeny Group, 2016; Begon et al., 2021; Poppenwimer et al., 2023). Our findings are consistent with the general angiosperm trend, with a disproportion of the number of annual (16%) versus perennial (84%) species in subg. *Esula* (Fig. S18, Table S1; Poppenwimer et al., 2023); the highest proportion of annuals has been reported (Hjertaas et al., 2023) for Poaceae (17%), Asteraceae (7%), and Fabaceae (5%), which makes *Euphorbia* exceptionally rich in annuals. The classical perspective is that the evolutionary trend in flowering plants is one from woody perennials, followed by the evolution of herbaceousness, and later annuality (Jeffrey, 1916; Stebbins, 1982; Hjertaas et al., 2023). However, subg. *Esula* follows a different pattern. Both BiSSE and HiSSE R reconstructed the ancestor of subg. *Esula* as an annual plant (Figs. 4, S16), with frequent transitions to the perennial life-history (up to 21) starting ∼14 Mya (0.051 events Mya^-1^), coincident with the origin of the first steppes and open grassy biomes (Ivanov et al., 2011; Manafzadeh et al., 2017), and only seven reversals, from perennial back to annual, inferred over 6 Mya to the present (0.037 events Mya^-1^, Figs. 4, S12A, S12B). Our reconstruction is very similar to that of genus *Androsace*, with annual ancestors that first evolved into cold-tolerant perennials during the Mid-Miocene in Central Asia, which was followed by a few reversals to an annual life-history associated with the driest niches (Boucher et al., 2012). The highly asymmetric transition rate inferred in subg. *Esula*, with a low rate of transition towards annuality (Fig. S11), has been documented in other clades such as the cosmopolitan family Montiaceae (Ogburn & Edwards, 2015), the pantropical genus *Begonia* L. (Kidner et al., 2016), or the widespread genus *Crassula* L. (Lu et al., 2022). The opposite pattern is found in the North American *Castilleja*, with a single transition to perenniality from an annual ancestor and later reversions to annuality (Tank & Olmstead, 2008); in *Lupinus*, Drummond et al. (2012a) found annual ancestors associated with lowland areas, with several transitions to perenniality concomitant with rapid diversification in mountain ranges and posterior reversals to annuality. The evolutionary drivers underlying transitions between annual and perennial life-histories are unclear. Boyko et al. (2023) found that the highest temperature of the warmest month is the main climatic factor that shapes the evolution of annuals. Poppenwimer et al. (2023) show similar findings, with annuals favoured in hot and dry regions like the Mediterranean biome, in which the temperature and precipitation of the driest quarter is the most important factor. This pattern is consistent across families, suggesting convergent evolution (Poppenwimer et al., 2023). Reversal transitions to perenniality and secondary woodiness have typically been associated with the colonisation of islands and insular habitats (Dulin & Kirchoff, 2010; Whittaker et al., 2017), a long-recognized phenomenon known as insular woodiness (Darwin, 1859; Wallace, 1878; Carlquist, 1974), with climatic moderation suggested to prompt the evolution of longer plant life spans in island species compared with their mainland relatives. One example of this pattern within subg. *Esula* is the dendroid shrub species in sects. *Aphyllis* and *Tithymalus*, perennial species endemic to the Macaronesian and Caribbean regions (Fig. 3) that have evolved secondary woodiness, as an adaptation to the absence of environmental constraints in the peculiarly stable environment of oceanic islands (Nürk et al., 2019). However, a switch from annuality to perenniality is also considered a key adaptation that promotes the colonisation of montane habitats (Drummond et al., 2012a; Hughes & Atchison, 2015). Perenniality (with or without woodiness) can represent an advantage in high-elevation habitats, where aseasonality and lower temperatures can select against the survival of annual plants, via reduced relative growth rates and elevated rates of seed mortality (Ogburn & Edwards, 2015). Perenniality in mountain habitats also allows species to take advantage of year-round light availability for growth and survival (Crawley & Harral, 1997). Several studies have found that elevation is the main factor influencing life-history composition (Pavón et al., 2000; Matteodo et al., 2013; Irl et al., 2020). A strong link between perenniality and montane habitats has been documented in the Andean *Lupinus* (Drummond, 2008), the Himalayan Gentianinae (Favre et al., 2010), the Western Palearctic *Scorzoneroides* Moench (Cruz-Mazo et al., 2009), and the worldwide Arabideae (Karl & Koch, 2013). Anest et al. (2021) suggested that frost is a strong selective factor that shaped the above ground plant architecture of subg. *Esula* under low average temperatures, especially in the species distributed in the mountains, mostly perennials (Fig. 4, Table S1). Our results also support this view, the evolution of perenniality from annual ancestors is not restricted solely to oceanic islands, but also associated with the colonisation of high-elevation mountain areas (Fig. 4; Kostikova et al., 2013; Ogburn & Edwards, 2015; Gehrke et al., 2016). We found significant differences in the proportion of annuals and perennials with respect to elevational distribution, with the high-elevation habitats dominated by perennial species (Fig. S18, Tables 4, S1). Conversely, annual taxa are favoured under anthropogenic or soil disturbance and in climates with high rainfall seasonality (Boyko et al., 2023; Poppenwimer et al., 2023). The transition from an annual to a perennial life-history, linked to the colonisation of mountain habitats, has also been associated with an increase in diversification rates due to ecological opportunity, i.e. reduced interspecific competition (Hughes & Atchison, 2015). Many of the groups mentioned above, as examples of the link between perenniality and high-elevation habitats, also represent cases of species radiations (Arabideae, Gentianinae, and *Lupinus*; Favre et al., 2010; Drummond et al., 2012a; Karl & Koch, 2013). In subg. *Esula*, most transitions to perenniality are recovered within the last 9 Mya (Figs. 4, S12A, S16, S17), concomitant with the colonisation of mountain habitats in the Late Miocene (Fig. 3), the surge in diversification rates in several clades (LSBDS, Fig. 4), and the decrease in relative extinction rates inferred by EBD models around this period (Figs. 2A, S6). Both BiSSE and HiSSE (R and RevBayes) support higher speciation and net diversification rates in perennial species compared to annual species (Figs. 4, S11, S14, S16, S17, Table 3). Other examples of this association are shown in *Androsace* (Roquet et al., 2013), *Crassula* (Lu et al., 2022), *Lupinus* (Drummond et al., 2012a) and, at a higher level, Delphinieae (Jabbour & Renner, 2012) and Saxifragales (Soltis et al., 2013).

A puzzling result is the striking difference in ancestral state reconstruction between RevBayes HiSSE (Figs. S15A–S15C) and BiSSE and R HiSSE (Figs. 4, S12A, S12B, S16, S17). The MRCAs of subg. *Esula* and nearly every section were reconstructed as a perennial plant with associated hidden state A (1A) in RevBayes HiSSE (Fig. S15A), with most transitions to annuality (0A, 0B) delayed to the last 6 million years (Fig. S15B); this is opposite to the pattern found by BiSSE and R HiSSE (Figs. 4, S16). Interestingly, those sections or clades within sections reconstructed as hidden state B (0B, 1B) in RevBayes HiSSE (Fig. S15A) are associated with accelerated diversification rates in LSBDS (Fig. 4); examples are sects. *Helioscopia* and *Tithymalus* (0B, 1B), sects. *Aphyllis*, *Esula,* and *Patellares* (1B), and clades within sects. *Myrsiniteae*, *Holophyllum*, *Paralias*, and *Pithyusa* (1B, Fig. S15A). Conversely, not every clade reconstructed as ancestrally perennial in BiSSE is associated with accelerated speciation rates in LSBDS: e.g. clade *E. isaurica*-*E. herniariifolia* (sect. *Tithymalus*) is ancestrally perennial but shows no increase in speciation rates (Figs. 4, S12A); clade *E. microsphaera*-*E. haussknechtii* (sect. *Helioscopia*) comprises only annual plants and exhibits accelerated diversification rates. Moreover, we found there was a strong effect of the hidden state in the speciation and net diversification rates, in which state B, irrespective of the focal trait, was associated with the highest rates (i.e. annual 0B larger than perennial 1A, Figs. S13, S14). Thus, despite finding support for an effect of the focal trait, perennial vs. annual in subg. *Esula* diversification rates (Figs. 4, S11, S16, S17), our results suggest that there is a hidden trait that is also driving this correlation, with the strongest effect for hidden state B (Figs. S13, S15A).

An important goal in the study of organismal diversity is to characterise traits based on whether they positively or negatively affect diversification rates (Sauquet & Magallón, 2018). However, there is limited evidence to suggest that specific traits consistently influence diversification on their own (Helmstetter et al., 2023). The complexity of how phenotypic traits influence diversification can be attributed to at least two factors. First, a given trait may have variable effects on diversification, sometimes increasing and sometimes decreasing it. Second, a primary trait may influence diversification indirectly through its associations with other traits (“synnovarions”; Donoghue & Sanderson, 2015). Therefore, the effects of traits on diversification are likely to be driven more by the interactions between groups of traits than by individual traits alone (Anderson et al., 2023). This could apply to our study, with perenniality associated with, at least, another trait that promoted the diversification of subg. *Esula*, as evidenced by the HiSSE results. López-Estrada et al. (2022) found a similar pattern in their analysis of host preferences in a clade of insects. They attributed the incongruence between their BiSSE and HiSSE results to the existence of a hidden character, phoretic behaviour, which was not considered in the initial analysis but was covarying with the focal trait, host specificity. Identifying this second trait in subg. *Esula*, however, has proven to be more difficult: we could not find a common trait that was shared by all clades exhibiting an increase in the rate of diversification rate in the LSBDS analysis (Fig. 4).

Beyond life-history, changes in traits such as reproductive strategy (e.g. seed dispersal and viability) or physiological adaptation (e.g. development of cold tolerance or drought resistance) may facilitate the colonisation of mountain habitats, or species persistence under rapidly changing environmental conditions (e.g. Meseguer et al., 2018; Moharrek et al., 2019). Further studies are needed to determine whether these additional traits contributed significantly to the observed patterns of diversification and geographic expansion in subg. *Esula*, and to gain a more complete understanding of the factors driving the evolution of this clade.

## 5 Conclusions

This study provides a densely-sampled and the most complete phylogenetic hypothesis for *Euphorbia* subgenus *Esula*. Our spatio-temporal reconstruction supports a major role for geological, climatic, and biome formation events during the Cenozoic in the origin of current distribution patterns. Subgenus *Esula* likely originated in the Western Palearctic region from boreotropical and Tethyan ancestors. Starting in the Mid-Late Miocene, coincident with a decrease in global temperatures, the subgenus dispersed to other continents, reaching Africa, Asia, the Nearctic, and Macaronesian regions. These migration events seem to be linked to the colonisation of newly arised mountain ranges, such as the QTP and the Iranian plateaus, and the evolution of the perennial life-history across many sections/clades within subg. *Esula*. Our results support that the evolution of perenniality was associated with higher speciation and net diversification rates, which might have been correlated with the ecological opportunities provided by the new high-elevation habitats. However, other hidden factors or traits might have been involved in the upsurge in diversification rates that, together with life-history, explain and drive diversification dynamics through time in subg. *Esula*.

## Author contributions

Irene Masa-Iranzo: Conceptualization, Methodology, Validation, Formal analysis, Investigation, Data Curation, Writing-Original Draft, Writing-Review & Editing, Visualization. Isabel Sanmartín: Conceptualization, Methodology, Software, Resources, Writing-Original Draft, Writing-Review & Editing, Supervision. Božo Frajman: Data Curation, Resources, Writing-Review & Editing. Andrea S. Meseguer: Funding acquisition, Resources, Methodology, Writing-Review & Editing, Supervision. Ricarda Riina: Conceptualization, Data Curation, Methodology, Resources, Writing-Original Draft, Writing-Review & Editing, Supervision.

## Informed consent statement

Informed consent was obtained from all subjects involved in the study.

## Data availability

All R and RevBayes scripts needed to replicate the biogeographic, diversification, and the association between elevation and life-history analyses can be found in Appendix S1– S5 and stored in the GitHub account https://github.com/isabelsanmartin/RevBayes-analysis-Esula

## Acknowledgements

We would like to thank Paloma Ruiz, Mario Rincón-Barrado, E. Karen López-Estrada, Fernando Useros, Ángela Aguado-Lara, Rubén González-Miguéns, Yolanda Turégano, and Miguel Blázquez for helpful bioinformatic resources; to David Criado for useful discussions; to María Martínez-Ríos, David Criado, and Carmen Soler-Zamora for their help with image editing. IM-I and AS were supported by the Atracción de Talento CAM program (2019-T1/AMB-12648). IS and RR were supported by project PID2019-108109GB-I00, funded by MCIN/AEI/10.13039/501100011033/ and FEDER ‘A way to make Europe’. The authors acknowledge the CIPRES portal for computational resources. We acknowledge support of the publication fee by the CSIC Open Access Publication Support Initiative through its Unit of Information Resources for Research (URICI).

