## Supplementary Table 2 for "The conquest and diversification of leafy spurges across the Holarctic and beyond: biogeography and evolution of life-history of *Euphorbia* subgenus *Esula*"

**Table S2**. Summary statistics from ITS, *ndhF*, and combined ITS + *ndhF* datasets for *Euphorbia* subg. *Esula* and outgroup species analysed in this study.

| | **ITS** | ***ndhF*** | **Combined ITS + *ndhF*** |
|---|---|---|---|
| **Number of accessions** | 328 | 237 | 341 |
| **Ingroup species** | 308 | 219 | 321 |
| **Outgroup species** | 19 | 19 | 19 |
| **Aligned length** | 793 | 1500 | 2293 |
| **Number of variable characters (%)** | 85.6 | 44.3 | 58 |
| **Missing data (%)** | 20.71 | 5.11 | 30.70 |
| **Parsimony-informative sites** | 419 | 414 | 833 |
| **Model of nucleotide substitution** | GTR + I + G | GTR + G | GTR + I + G & GTR + G |
