## Supplementary Table 3 for "The conquest and diversification of leafy spurges across the Holarctic and beyond: biogeography and evolution of life-history of *Euphorbia* subgenus *Esula*"

**Table S3.** Clade posterior probabilities (PP), mean, and 95 % highest posterior density (HPD) credibility intervals values in million years ago (Mya) of node ages from the divergence times analysis run in BEAST of *Euphorbia* subg. *Esula* and other nodes marked in Figs. 2G and 2H.

| **Clades/Nodes** | **PP** | **Mean (Mya)** | **95% HPD** |
|---|---|---|---|
| Node A (*Euphorbia*) | 1 | 47.19 | 52.24–42.56 |
| Node B (*Euphorbia* subg. *Esula*) | 1 | 40.75 | 45.10–36.47 |
| Node C | 1 | 37.64 | 41.95–33.69 |
| Node D (*E.* sects. *Lagascae* + *Lathyris*) | 0.94 | 34.07 | 41.80–23.21 |
| Node E | 1 | 33.35 | 37.46–29.43 |
| Node F (*E.* sect. *Sclerocyathium*) | 1 | 12.13 | 18.93–7.39 |
| Node G (*E.* sect. *Holophyllum*) | 1 | 10.38 | 17.65–5.69 |
| Node L (*E.* sect. *Aphyllis*) | 0.77 | 5.74 | 7.12–3.59 |
| *E.* sects. *Aphyllis* + *Exiguae* 1 | 0.71 | 10.13 | 13.50–7.02 |
| *E.* sect. *Aphyllis* to sect. *Exiguae* 2 (*E. exigua*) | 1 | 13.18 | 17.07–9.64 |
| *E.* sect. *Aphyllis* to sect. *Exiguae* 2 (*E. medicaginea*) | 0.37 | 12.27 | 16.11–9.05 |
| *E.* sect. *Aphyllis* to sect. *Patellares* | 1 | 16.81 | 21.14–12.95 |
| *E.* sect. *Arvales* | 1 | 7.42 | 10.53–4.41 |
| *E.* sects. *Arvales* + *Esula* | 1 | 11.46 | 15.81–7.73 |
| *E.* sect. *Biumbellatae* | 1 | 3.59 | 6.67–1.67 |
| *E.* sect. *Biumbellatae* to sect. *Exiguae* 2 (*E. medicaginea*) | 0.21 | 10.04 | 13.17–6.99 |
| *E.* sect. *Biumbellatae* to sect. *Guyonianae* | 0.88 | 9.37 | 12.60–6.39 |
| *E.* sect. *Calyptratae* | 1 | 13.63 | 21.26–5.88 |
| *E.* sect. *Calyptratae* + remaining sections | 0.95 | 27.84 | 32.58–23.54 |
| *E.* sect. *Chylogala* | 1 | 13.21 | 19.20–7.87 |
| *E.* sect. *Chylogala* + remaining sections | 0.92 | 26.14 | 30.46–21.81 |
| *E.* sect. *Esula* | 1 | 6.90 | 9.60–5.03 |
| *E.* sect. *Esula* to sect. *Paralias* | 0.96 | 20.16 | 24.39–15.99 |
| *E.* sect. *Esula* to sect. *Patellares* | 1 | 22.02 | 26.49–18.13 |
| *E.* sect. *Exiguae* 1 | 1 | 3.46 | 6.14–1.27 |
| *E.* sects. *Guyonianae* + *Pachycladae* | 0.82 | 8.01 | 11.16–5 |
| *E.* sect. *Helioscopia* | 1 | 20.37 | 25.44–15.75 |
| *E.* sects. *Helioscopia* + *Holophyllum* | 1 | 24.56 | 29.75–19.33 |
| *E.* sect. *Herpetorrhizae* | 0.99 | 12.30 | 16.61–8.32 |

| **Clades/Nodes** | **PP** | **Mean (Mya)** | **95% HPD** |
|---|---|---|---|
| *E.* sect. *Lagascae* | 1 | 20.06 | 30.62–9.22 |
| *E.* sect. *Myrsiniteae* | 1 | 11.80 | 17.94–6.75 |
| *E.* sects. *Myrsiniteae* + *Pithyusa* | 1 | 23.43 | 30.12–17.13 |
| *E.* sect. *Pachycladae* | 1 | 4.08 | 6.98–1.72 |
| *E.* sect. *Paralias* | 1 | 5.91 | 8.92–3.46 |
| *E.* sects. *Paralias* + *Tithymalus* | 1 | 15.67 | 19.86–11.63 |
| *E.* sect. *Patellares* | 1 | 2.46 | 3.85–1.49 |
| *E.* sect. *Pithyusa* | 1 | 12.95 | 17.50–9.43 |
| *E.* sect. *Sclerocyathium* + remaining sections | 1 | 29.68 | 34.24–25.35 |
| *E.* sect. *Tithymalus* | 1 | 10.44 | 13.81–7.40 |
