## Supplementary Table 5 for "The conquest and diversification of leafy spurges across the Holarctic and beyond: biogeography and evolution of life-history of *Euphorbia* subgenus *Esula*"

**Table S5.** Ancestral geographic range of major clades of *Euphorbia* subg. *Esula* inferred with Bayesian dispersal extinction cladogenesis under the M1 model. Only the first three areas with the highest posterior probability (PP) value are given. Area codes and node letters followed those in Fig. 3.

| | Ancestral range reconstruction |
|---|---|
| **Nodes** | **Area, PP** |
| Node A (*Euphorbia*) | WP: 0.241, AFR−WP: 0.1505, WP−NAM: 0.1438. |
| Node B (*E.* subg. *Esula*) | WP: 0.739, WP−MAC: 0.0732, AFR−WP: 0.0373. |
| Node C | WP: 0.8216, WP−MAC: 0.0559, WP−EP: 0.0333. |
| Node D (*E.* sects. *Lagascae* + *Lathyris*) | WP: 0.719, WP−MAC: 0.0892, MAC: 0.0466. |
| Node E | WP: 0.8802, WP−MAC: 0.0386, WP−EP: 0.0213. |
| Node F (*E.* sect. *Sclerocyathium*) | AFR: 0.2157, WP: 0.1119, NAM: 0.0945. |
| Node G (*E.* sect. *Holophyllum*) | EP: 0.3302, AFR−EP: 0.1704, WP−EP: 0.1558. |
| Node L (*E.* sect. *Aphyllis*) | AFR−MAC: 0.3302, MAC: 0.225, AFR−WP−MAC: 0.225. |
| *E.* sects. *Aphyllis* + *Exiguae* 1 | WP: 0.715, WP−MAC: 0.1651, AFR−WP: 0.0559. |
| *E.* sect. *Aphyllis* to sect. *Exiguae* 2 (*E. exigua*) | WP: 0.9254, WP−MAC: 0.0479, AFR−WP: 0.0226. |
| *E.* sect. *Aphyllis* to sect. *Exiguae* 2 (*E. medicaginea*) | WP: 0.9174, WP−MAC: 0.0613, AFR−WP: 0.016. |
| *E.* sect. *Aphyllis* to sect. *Patellares* | WP: 0.9108, AFR−WP: 0.0426, WP−MAC: 0.0333. |
| *E.* sect. *Arvales* | WP: 0.7989, WP−MAC: 0.0746, WP−EP: 0.0439. |
| *E.* sects. *Arvales* + *Esula* | WP: 0.7803, AFR−WP: 0.0985, WP−MAC: 0.0506. |
| *E.* sect. *Biumbellatae* | WP: 0.9694, AFR−WP: 0.0093, WP−MAC: 0.0093. |
| *E.* sect. *Biumbellatae* to sect. *Pachycladae* | WP: 0.9534, WP−MAC: 0.02, AFR−WP: 0.0133. |
| *E.* sect. *Calyptratae* | WP: 0.743, AFR−WP: 0.0719, WP−MAC: 0.0652. |
| *E.* sect. *Calyptratae* + remaining sections | WP: 0.8961, WP−MAC: 0.0346, AFR−WP: 0.028. |
| *E.* sect. *Chylogala* | WP: 0.8003, AFR−WP: 0.0799, WP−MAC: 0.0573. |

**Ancestral range reconstruction**

| **Nodes** | **Area, PP** |
|---|---|
| *E.* sect. *Chylogala* + remaining sections | WP: 0.9427, WP−MAC: 0.0226, AFR−WP: 0.0146. |
| *E.* sect. *Esula* | WP−EP: 0.3915, WP: 0.2304, AFR−WP−EP: 0.2264. |
| *E.* sect. *Esula* to sect. *Paralias* | WP: 0.9281, AFR−WP: 0.032, WP−MAC: 0.0293. |
| *E.* sect. *Esula* to sect. *Patellares* | WP: 0.9374, WP−MAC: 0.032, AFR−WP: 0.0226. |
| *E.* sect. *Exiguae* 1 | WP: 0.9454, WP−MAC: 0.0213, AFR−WP: 0.0133. |
| *E.* sects. *Guyonianae* + *Pachycladae* | WP: 0.9521, AFR−WP: 0.02, WP−MAC: 0.0173. |
| *E.* sect. *Helioscopia* | WP: 0.8376, AFR−WP: 0.0546, WP−MAC: 0.0533. |
| *E.* sects. *Helioscopia* + *Holophyllum* | WP: 0.6618, AFR−WP: 0.0959, WP−MAC: 0.0945. |
| *E.* sect. *Herpetorrhizae* | WP: 0.6964, AFR−WP: 0.1052, WP−EP: 0.0599. |
| *E.* sect. *Lagascae* | WP: 0.6751, WP−MAC: 0.1079, AFR−WP: 0.0719. |
| *E.* sect. *Myrsiniteae* | WP: 0.8708, WP−MAC: 0.0586, AFR−WP: 0.0386. |
| *E.* sects. *Myrsiniteae* + *Pithyusa* | WP: 0.8242, WP−MAC: 0.0866, AFR−WP: 0.0453. |
| *E.* sect. *Pachycladae* | WP: 0.6405, WP−MAC: 0.1704, AFR−WP: 0.0945. |
| *E.* sect. *Pachycladae* to sect. *Exiguae* 2 (*E. medicaginea*) | WP: 0.9534, WP−MAC: 0.02, AFR−WP: 0.0133. |
| *E.* sect. *Paralias* | WP: 0.6511, WP−MAC: 0.1411, WP−NAM: 0.0985. |
| *E.* sects. *Paralias* + *Tithymalus* | WP: 0.9174, WP−MAC: 0.036, AFR−WP: 0.0293. |
| *E.* sect. *Patellares* | WP: 0.9534, WP−MAC: 0.0213, AFR−WP: 0.0173. |
| *E.* sect. *Pithyusa* | WP: 0.8868, AFR−WP: 0.0493, WP−MAC: 0.0439. |
| *E.* sect. *Sclerocyathium* + remaining sections | WP: 0.8296, WP−MAC: 0.0519, AFR−WP: 0.0426. |
| *E.* sect. *Tithymalus* | WP: 0.8602, WP−NAM: 0.0546, AFR−WP: 0.0426. |
