## Supplementary File 1 for "The conquest and diversification of leafy spurges across the Holarctic and beyond: biogeography and evolution of life-history of *Euphorbia* subgenus *Esula*"

**Supplementary File 1**. Taxon, origin, voucher information (herbarium acronym), and ITS and *ndhF* GenBank accession numbers for samples included in the phylogenetic analyses. Species names are up to date under the most recent taxonomic revisions.

***Calycopeplus casuarinoides*** L.S. Sm., cult., *V. Steinmann 1407* (RSA), AF537580, JN249080; ***Euphorbia acanthodes*** Akhani, Iran, *H. Akhani 14782* (IRAN), KC212160, –; ***Euphorbia acanthothamnos*** Heldr. & Sart. ex Boiss., Greece, *R. Riina 1563* (MICH), JQ750879, JQ750756; ***Euphorbia adenochlora*** C. Morren & Decne., Japan, *Oita 01393* (SAPS), KC212161, –; ***Euphorbia agraria*** M. Bieb., U.S.A., *R. McGregor 31711* (RM), KC212163, KC212434; ***Euphorbia akenocarpa*** Guss., Spain, *L. Barres & al. BCN53041* (BCN), HQ900574, KC212435; ***Euphorbia akmanii*** I. Genç & Kültür, Turkey, *F. Ehrendorfer & al. 15951* (IB), ON908612, –; ***Euphorbia alaica*** (Prokh.) Prokh., Kyrgyzstan, *G. Lazkov s.n.* (LE), KC212165, KC212436; ***Euphorbia alatavica*** Boiss., Kyrgyzstan, *G. Lazkov s.n.* (LE), GU953741, –; ***Euphorbia aleppica*** L. **1,** Cyprus, *R. Hand 5797* (B), MT957500, –; ***Euphorbia aleppica*** L. **2,** Iran, *Y. Salmaki & al. 39899* (TUH), –, KC212438; ***Euphorbia alpina*** Ledeb., Russia, *P. Berry 7983* (MICH), KC212168, –; ***Euphorbia alta*** Norton, U.S.A., *A. Sanders 5905* (RSA), AF537553, –; ***Euphorbia altaica*** Ledeb., Russia, *D. Geltman 27* (LE), GU979429, –; ***Euphorbia altotibetica*** Paulsen in S. Hedin, China, *H. Ting Meng & al. 400* (PE), KC212171, –; ***Euphorbia amygdaloides*** L., Bosnia, *M. Turjak & B. Frajman 11821* (IB), JN010024, KC212439; ***Euphorbia anachoreta*** Svent., Spain, *B. Dorsey 18* (MICH), KC212173, KC212440; ***Euphorbia angulata*** Jacq., Germany, *J. Esser 06–26* (M), KC212175, KC212441; ***Euphorbia antso*** Denis, cult., *V. Steinmann 1473* (RSA), JN250112, JN249102; ***Euphorbia aphylla*** Brouss. ex Willd., Spain, *B. Dorsey 4* (MICH), JN250113, JN249103; ***Euphorbia apios*** L., Greece, *W. Gutermann 25543* (Herb. Gutermann), JN010027, –; ***Euphorbia appariciana*** Rizzini, cult., *V. Steinmann 1442* (RSA), JN250114, JN249104; ***Euphorbia arguta*** Banks & Sol., Cyprus, *R. Meikle 2237* (LE), KC212176, –; ***Euphorbia aristata*** Schmalh., Russia, *D. Geltman s.n.* (LE), GU979434, –; ***Euphorbia arvalis*** subsp. ***arvalis*** Boiss. & Heldr., Iran, *V. Mozaffarian & M. Norouzi 34533* (TARI), KC212177, KC212442; ***Euphorbia aserbajdzhanica*** Bordz., Iran, *A. Pahlevani & Asef 47796* (IRAN), KC212180, KC212444; ***Euphorbia atropurpurea*** Brouss. ex Willd. **1,** Spain, *J. Molero 5/2007* (BCN), HQ900579, –; ***Euphorbia atropurpurea*** Brouss. ex Willd. **2,** Spain, *B. Dorsey 26* (MICH), –, KC212445; ***Euphorbia aucheri*** Boiss., Iran, *S. Zarre & al. 38189* (TUH), KC212181, KC212446; ***Euphorbia aulacosperma*** Boiss., Iran, *E. Eskandari 53764* (IRAN), LN680639, LN680648; ***Euphorbia austro-iranica*** Pahlevani, Iran, *Govanz Moussavi & Tehrani 47288* (IRAN), LN998089, LN998102; ***Euphorbia austrotexana*** *M. Mayfield*, U.S.A., *W. Carr 12504* (KSC, MICH), KJ149580, KJ149625; ***Euphorbia azorica*** Hochst. in M.A. Seubert, Portugal, *J. Molero & al. BCN 86828* (BCN), KC212182, KC212448; ***Euphorbia balsamifera*** Aiton, Spain, *B. Dorsey 3* (MICH), JN250117, JN249107; ***Euphorbia barrelieri*** Savi, Italy, *R. Vilatersana & al. 1235* (BC), KC212183, KC212450; ***Euphorbia beamanii*** M. C. Johnst., Mexico, *M. Mayfield 1903* (MICH), KJ149583, KJ149627; ***Euphorbia berotica*** N.E. Br., Angola, *P. Bruyns 10685* (BOL), KC212184, KC212451; ***Euphorbia berthelotii*** Bolle ex Boiss., Spain, *J. Molero 25/2007* (BCN), HQ900585, –; ***Euphorbia bivonae*** Steud., Italy, *R. Riina 1902* (IB), KT071809, –; ***Euphorbia biumbellata*** Poir., Spain, *J. Molero 26/2007* (BCN), KC212186, KC212452; ***Euphorbia boetica*** Boiss. **1,** Spain, *J. Molero 18/2007* (BCN), KC212187, –; ***Euphorbia***

***boetica*** Boiss. **2,** Spain, *J. Molero 22/2007* (BCN), –, KC212453; ***Euphorbia borealis*** Baikov, Russia, *P. Berry 7982* (MICH), KC212192, KC212455; ***Euphorbia bourgaeana*** J. Gay ex Boiss., Spain, *B. Dorsey 31* (MICH), KC212193, KC212456; ***Euphorbia brachycera*** Engelm., U.S.A., *B. van Ee 1010* (MICH), KC212194, KC212458; ***Euphorbia bravoana*** Svent., Spain, *A. Fernández-Lopez & J. Molero 08/2007* (BCN), HQ900588, –; ***Euphorbia briquetii*** Emb. & Maire, Morocco, *R. Riina 1801* (MICH), KC212195, KC212459; ***Euphorbia buchtormensis*** Ledeb., Kazakhstan, *S. Smirnov s.n.* (LE), KC212196, –; ***Euphorbia buhsei*** Boiss. **1,** Turkmenistan, *D. Kurbanov s.n.* (LE), KC212197, –; ***Euphorbia buhsei*** Boiss. **2,** Iran, *Pahlevani 53823* (IRAN), –, LN998105; ***Euphorbia bungei*** Boiss., Iran, *Y. Salmaki 39930* (TUH), KC212199, KC212460; ***Euphorbia caeladenia*** Boiss., Iran, *A. Pahlevani & Bahramishad 53777* (IRAN), KC212200, KC212461; ***Euphorbia caesia*** Kar. & Kir., Austria, *W. Till 12591* (IB), JN010031, KC212462; ***Euphorbia calamiformis*** P.R.O. Bally & S. Carter, Kenya, *S. Carter & B. Stannard 558* (K), HQ900589, –; ***Euphorbia calyptrata*** Coss. & Durieu, Morocco, *R. Riina 1810* (MICH), JN250123, JN249113; ***Euphorbia capitulata*** Rchb., Bosnia, *M. Turjak & B. Frajman 11838* (IB), JN010032, KC212465; ***Euphorbia carniolica*** Jacq., Bosnia, *M. Turjak & B. Frajman 11836* (IB), JN010033, –; ***Euphorbia cashmeriana*** Royle, Afghanistan, *L. Edelberg 781* (W), HQ900592, –; ***Euphorbia caspica*** Frajman & Pahlevani, Iran, *A. Pahlevani & Torabi 76626* (IRAN), ON908621, –; ***Euphorbia cassia*** Boiss., Cyprus, *M. Galbany & al. 2035 (duplicate)* (BC), KC212202, KC212467; ***Euphorbia ceratocarpa*** Ten., Italy, *R. Vilatersana & al. 1141* (BC), KC212203, KC212468; ***Euphorbia ceratocarpa*** Ten., Italy, *R. Vilatersana & al. 1161* (BC), HQ900597, KC212476; ***Euphorbia chamaepeplus*** Boiss. & Gaill., Jordan, *M. Staudinger 209a18 (W),* LN680640, LN680649; ***Euphorbia chamaesula*** Boiss., U.S.A., *J. Peirson 865* (MICH), KJ149589, KJ149634; ***Euphorbia characias*** subsp. ***characias*** L., France, *P. Berry 7917* (MICH), KC212205, KC212470; ***Euphorbia cheiradenia*** Boiss. & Hohen. **1,** Iran, *B. Frajman, M. Falch & Ch. Gilli 14278* (IB), MT957502, –; ***Euphorbia cheiradenia*** Boiss. & Hohen. **2,** Iran, *Y. Salmaki & S. Zarre 39920* (TUH), –, KC212471; ***Euphorbia clementei*** Boiss., Morocco, *R. Riina 1789* (MICH), KC212206, –; ***Euphorbia commutata*** Engelm. ex A. Gray, Canada, *M. Oldham 14724* (MICH), KC212207, KC212472; ***Euphorbia condylocarpa*** M. Bieb., Iran, *Y. Salmaki & al. 39955* (TUH), KC212208, KC212473; ***Euphorbia coniosperma*** Boiss. & Buhse**,** Armenia, *L. Vakhtina s.n.* (LE), KC212209, –; ***Euphorbia connata*** Boiss., Iran, *Y. Salmaki & al. 39937* (TUH), KC212211, KC212475; ***Euphorbia consanguinea*** Schrenk, Turkmenistan, *V. Botschantsev 410* (LE), KC212212, –; ***Euphorbia craspedia*** Boiss., Iran, *Y. Salmaki & S. Zarre 14327* (TUH), KC212213, KC212477; ***Euphorbia creberrima*** McVaugh, Mexico, *J. Rzedowski 17965* (MICH), –, KJ149638; ***Euphorbia crenulata*** Engelm., U.S.A., *F. Brunett 313* (MICH), KJ149593, KJ149640; ***Euphorbia cressoides*** M. C. Johnst., Mexico, *M. Johnston & al. 9551i* (TEX), –, KJ149642; ***Euphorbia cyparissias*** L., France, *P. Berry 7916* (MICH), KC212215, KC212480; ***Euphorbia cyparissioides*** Pax, Tanzania, *J. Morawetz 463* (MICH), KC212216, KC212481; ***Euphorbia cyrtophylla*** (Prokh.) Prokh., Tadjikistan, *I. Shibkova 2944* (LE), KC212217, –; ***Euphorbia deflexa*** Sibth. & Sm., Greece, *B. Frajman & P. Schönswetter 11676* (IB), JN010038, –; ***Euphorbia deltobracteata*** (Prokh.) Prokh., Iran, *M. Eskandari 53773* (IRAN), KC212218, KC212482; ***Euphorbia dendroides*** L., Greece, *R. Riina 1555* (MICH), JN250135, JN249125; ***Euphorbia densa*** Schrenk, Iran, *M. Eskandari s.n.* (MICH), KC212220, KC212483; ***Euphorbia densiusculiformis*** Popov, Uzbekistan, *R. Kamelin 223* (LE), KC212224, KC212487; ***Euphorbia denticulata*** Lam. **1,** Iran, *K. Rechinger 47845* (W), HQ900601, –; ***Euphorbia denticulata*** Lam. **2,** Iran, *Y. Salmaki & al. 41008* (TUH), –, KC212488; ***Euphorbia depauperata*** Hochst. ex A. Rich., Tanzania,

*J. Morawetz 466* (MICH), KC212225, KC212489; ***Euphorbia dimorphocaulon*** P.H. Davis, Turkey, *R. Riina 1673* (MA), KC212228, KC212492; ***Euphorbia discolor*** Ledeb., Russia, *P. Schönswetter & A. Tribsch TG-12* (WU), JN010040, KC212493; ***Euphorbia dracunculoides*** subsp. ***inconspicua*** (Ball) Maire, Algeria, *J. Aldasoro 9811* (BCN), KC212229, KC212494; ***Euphorbia dregeana*** E. Mey. ex Boiss., South Africa, *R. Becker 897* (PRE), JN250140, JN249130; ***Euphorbia dubovikii*** Oudejans, Russia, *D. Geltman 23* (LE), GU984308, –; ***Euphorbia dulcis*** L., Austria, *F. Schuhwerk 06–236* (M), KC212231, KC212496; ***Euphorbia dumalis*** S. Carter, Ethiopia, *J. Aldasoro 10386* (BCN), KC212232, KC212497; ***Euphorbia ebracteolata*** Hayata **1,** South Korea, *Park 1010*, EU659768, –; ***Euphorbia ebracteolata*** Hayata, no country, *no collector,* –, MT830860; ***Euphorbia emirnensis*** Baker, Madagascar, *T. Haevermans 95* (P), AJ508964, –; ***Euphorbia eriophora*** Boiss. **1,** Iran, *M. Iranshahr 46720* (IRAN), KC212238, –; ***Euphorbia eriophora*** Boiss. **2,** Iran, *V. Mozaffarian 97361* (TARI), –, LN680650; ***Euphorbia erubescens*** Boiss., Iran, *Y. Salmaki & al. 39954* (TUH), KC212240, KC212501; ***Euphorbia erythradenia*** Boiss., Iran, *A. Pahlevani & Bahramishad 53336* (IRAN), KC212241, –; ***Euphorbia erythrodon*** Boiss. & Heldr., Turkey, *M. Niketić & M. Jovanović 13875* (IB), OL640130, –; ***Euphorbia espinosa*** Pax, cult., *V. Steinmann 1494* (RSA), AF537416, AF538190; ***Euphorbia esula*** L., U.S.A., *P. Berry 7974* (MICH), KC212244, KC212504; ***Euphorbia esuliformis*** S.Schauer, Mexico, *M. Mayfield 1892 (Redo)* (MICH), KJ149598, KJ149645; ***Euphorbia eugeniae*** Prokh., Georgia, *D. Geltman 57a* (LE), GU979428, –; ***Euphorbia exigua*** L., Portugal, *R. Riina 1550* (MICH), KC212247, KC212506; ***Euphorbia falcata*** L., Bulgaria, *B. Frajman & P. Schönswetter 11347* (IB), JN010046, KC212507; ***Euphorbia fauriei*** H. Lév. & Vaniot, South Korea, *Tho & J.H. Kim 2001–0002*, EU659767, –; ***Euphorbia ferdowsiana*** Pahlevani, Iran, *Ayatollahi & Zangoei 22585* (IRAN), LN680642, LN680651; ***Euphorbia ferganensis*** B. Fedtsch., Kyrgyzstan, *R. Kamelin 1311* (LE), KC212248, –; ***Euphorbia fischeriana*** Steudel, China, *C. L. Zhang 150782180530034LY* (PE), MT635184, –; ***Euphorbia fistulosa*** M.S. Khan, Turkey, *P. Davis 28230* (LE), KC212249, KC212509; ***Euphorbia flavicoma*** DC., Spain, *J. Molero & al. BCN53617* (MICH), JN250152, JN249142; ***Euphorbia fontqueriana*** Greuter, Spain, *B. Frajman 12546* (IB), JN010047, –; ***Euphorbia fragifera*** Jan, cult., *R. Riina 1841* (MA), KC212255, KC212515; ***Euphorbia franchetii*** B. Fedtsch., Iran, *Y. Salmaki & al. 38179* (TUH), KC212256, KC212516; ***Euphorbia furcatifolia*** M.G. Gilbert, Ethiopia, *J. Aldasoro 10266* (BCN), KC212257, KC212517; ***Euphorbia furcillata*** Kunth, Mexico, *M. Salinas & al. 800* (MEXU), –, KC212518; ***Euphorbia gaillardotii*** Boiss. & Blanche, Turkey, *H. Haussknecht s.n* (LE), KC212258, KC212519; ***Euphorbia gasparrinii*** Boiss., Italy, *B. Frajman & S. Bogdanović 13186* (IB), MK088031, –; ***Euphorbia gayi*** Salis, Italy, *B. Frajman 16799* (IB), ON511349, –; ***Euphorbia gedrosiaca*** Rech.f., Aellen & Esfand., Iran, *A. Pahlevani & Bahramishad 53344* (IRAN), KC212259, –; ***Euphorbia genistoides*** P. J. Bergius, South Africa, *J. Morawetz 301* (MICH), KC212260, KC212520; ***Euphorbia glaberrima*** (K. Koch) K. Koch, Russia, *G. Stohr 23b* (B), ON908637, –; ***Euphorbia glareosa*** Pall. ex M. Bieb., Austria, *B. Frajman & P. Schönswetter 11100* (IB), JN010050, KC212522; ***Euphorbia glauca*** G. Forst., New Zealand, *P. Garnock-Jones 2844*, KC212261, KC212523; ***Euphorbia gossypina*** Pax, Tanzania, *J. Morawetz 439* (MICH), KC212264, KC212526; ***Euphorbia graminifolia*** Vill., France, *C. Voisin 17098* (IB), ON511353, –; ***Euphorbia gregersenii*** K. Malý ex Beck, Bosnia, *B. Frajman & P. Schönswetter 12554* (IB), JN010051, KC212528; ***Euphorbia greggii*** Engelm. ex Boiss., Mexico, *M. Mayfield 2250* (KSC, MICH), KJ149600, KJ149647; ***Euphorbia grisophylla*** M. S. Khan, Iran, *Maroofi 55797* (IRAN), LN680644, LN680653; ***Euphorbia grossheimii*** (Prokh.) Prokh., Iran, *Y. Salmaki & S.*

*Zarre 41012* (TUH), KC212266, KC212529; ***Euphorbia guyoniana*** Boiss. & Reut., Morocco, *R. Riina 1797* (MICH), JN250164, JN249155; ***Euphorbia gypsicola*** Rech.f. & Aellen, Iran, *Y. Salmaki & S. Zarre 41011* (TUH), KC212268, KC212530; ***Euphorbia haussknechtii*** Boiss., Syria, *H. Haussknecht s.n* (LE), KC212269, KC212531; ***Euphorbia hebecarpa*** Boiss., Iran, *Y. Salmaki & S. Zarre 31841* (TUH), KC212270, KC212532; ***Euphorbia heldreichii*** Orph., Greece, *B. Frajman 15894* (IB), MN954390, –; ***Euphorbia helioscopia*** L., Austria, *W. Till 4529* (WU), JN010052, KC212533; ***Euphorbia helleri*** Millsp., U.S.A., *M. Mayfield 2142* (MICH), KJ149602, KJ149650; ***Euphorbia hercegovina*** Beck, Bosnia, *B. Frajman & P. Schönswetter 12135* (IB), JN010053, KC212535; ***Euphorbia herniariifolia*** Willd., Greece, *B. Frajman & P. Schönswetter 11668* (IB), JN010054, KC212536; ***Euphorbia heteradena*** Jaub. & Spach, Iran, *S. Zarre & Y. Salmaki 39893* (TUH), KC212273, KC212537; ***Euphorbia hierosolymitana*** Boiss., Israel, *A. Danin & al., 2nd Iter Medit. 12019* (B), KT071810, –; ***Euphorbia himalayensis*** (Klotzsch) Boiss., China, *W. Jin 020* (MICH), KC212393, KC212648; ***Euphorbia hirsuta*** L., Spain, *R. Riina 1769 (duplicate)* (MICH), JN250171, JN249161; ***Euphorbia humilis*** Ledeb., Kyrgyzstan, *G. Lazkov s.n.* (LE), GU984329, –; ***Euphorbia hyberna*** L., Spain/France, *B. Frajman & P. Schönswetter 11436* (IB), JN010056, KC212538; ***Euphorbia hylonoma*** Hand.-Mazz., no country, *no collector,* EU659770, –; ***Euphorbia iberica*** Boiss., Iran, *A. Pahlevani & Asef 54569* (IRAN), KC212275, KC212540; ***Euphorbia illirica*** Lam. Slovakia, *B. Frajman & P. Schönswetter 12416* (IB), JN010100, KC212634; ***Euphorbia inderiensis*** Less. ex Kar. & Kir., Iran, *A. Pahlevani 53762* (IRAN), KC212277, KC212544; ***Euphorbia isatidifolia*** Lam., Spain, *R. Riina 1867* (MA), KC212279, KC212546; ***Euphorbia isaurica*** M.S. Khan, Turkey, *P. Davis 16189* (E), –, KC212547; ***Euphorbia ivanjohnstonii* f. *longifolia*** B.L. Turner, Mexico, *J. Henrickson 15643* (TEX), –, KJ149653; ***Euphorbia japygica*** Ten., Italy, *D. Regele 15814* (IB), OL640146, –; ***Euphorbia japygica* subsp. *prostrata*** (Fiori) Del Guacchio & Frajman, Italy, *B. Frajman & S. Bogdanović* 13191 (IB), MK091300, –; ***Euphorbia jaxartica*** (Prokh.) Krylov, Kyrgyzstan, *G. Lazkov s.n.* (LE), KC212280, –; ***Euphorbia jolkinii*** Boiss., China, *N. Yang 0935* (KUN), KC212282, KC212548; ***Euphorbia kanaorica*** Boiss., Tadjikistan, *R. Kamelin s.n.* (LE), KC212285, –; ***Euphorbia kansuensis*** Prokhanov, China, *L. Zhang 150221140516127LY* (PE), MT635178, –; ***Euphorbia kansui*** S. L. Liou, no country, *no collector,* –, MH392274; ***Euphorbia kerneri*** Huter ex A. Kern., Italy, *B. Frajman & P. Schönswetter 12076* (IB), JN010058, KC212549; ***Euphorbia khabrica*** Pahlevani, Iran, *A. Pahlevani & Bahramishad 55152* (IRAN), KC212382, KC212635; ***Euphorbia komaroviana*** Prokh., Russia, *P. Efimov & T. Moskalyuk 4* (LE), GU979439, –; ***Euphorbia kopetdaghi*** (Prokh.) Prokh., Iran, *Y. Salmaki & al. 38185* (TUH), KC212287, KC212550; ***Euphorbia kotschyana*** Fenzl, Turkey, *Classen-Bockhoff 2001-01442* (W), ON908639, –; ***Euphorbia kozlovii*** Prokh., Mongolia, *V. Grubov & al. 43* (LE), KC212288, –; ***Euphorbia kraussiana*** Bernh. ex Krauss, South Africa, *P. Bruyns 11289* (BOL), KC212290, –; ***Euphorbia lagascae*** Spreng., Spain, *J. Molero & al. 18/2008 (duplicate)* (BCN), KC212291, KC212551; ***Euphorbia lamarckii*** Sweet **1,** Spain, *B. Dorsey 23* (MICH), –, KC212553; ***Euphorbia lamarckii*** Sweet **2,** Spain, *J. Molero 6/2007* (BCN), HQ900619, –; ***Euphorbia lamprocarpa*** (Prokh.) Prokh., Kazakhstan, *D. Geltman 199* (LE), KC212294, KC212554; ***Euphorbia larica*** Boiss., Oman, *J. Morawetz 350* (MICH), HQ900620, KC212430; ***Euphorbia lateriflora*** Schumach., Ghana, *C. Jongkind & al. 1720* (MO), JN250179, JN249169; ***Euphorbia lathyris*** L., U.S.A., *K. Wurdack 5558* (US), JN250180, JN249170; ***Euphorbia latifolia*** Ledeb., Russia, *D. Geltman 42* (LE), GU984316, –; ***Euphorbia lemesiana*** Hadijk *& al.*, Cyprus, *B. Frajman & P. Schönswetter 12707* (IB), KT071814, –; ***Euphorbia leptocaula*** Boiss., Russia, *D.*

*Matveev s.n.* (LE), GU984320, –; ***Euphorbia leucocephala*** Lotsy, cult., *P. Berry 7841* (MICH), JN250182, JN249172; ***Euphorbia lipskyi*** (Prokh.) Prokh., Tadjikistan, *I. Shibkova & al. 1392* (LE), KC212295, –; ***Euphorbia lomelii*** V.W. Steinm., Mexico, *B. van Ee 703* (MICH), JN250184, JN249174; ***Euphorbia longicruris*** Scheele, U.S.A., *C. Ferguson 459* (LSU), KJ149604, KJ149654; ***Euphorbia lucida*** Waldst. & Kit., Slovenia, *B. Frajman 12023* (IB), JN010060, KC212557; ***Euphorbia lucorum*** Rupr., China, *B.U. Oh s.n.*, EU659771, –; ***Euphorbia lunulata*** Bunge, South Korea, *Oh & Kim 2001–0002*, EU659752, –; ***Euphorbia lurida*** Engelm., U.S.A., *M. Mayfield 3357* (KSC, MICH), KJ149606, KJ149656; ***Euphorbia macrocarpa*** Boiss. & Buhse, Iran, *Y. Salmaki & al. 39561* (TUH), KC212298, KC212559; ***Euphorbia macroceras*** Fisch. & C.A. Mey., Georgia, *A. Tribsch & al. 10916* (WU), JN010062, KC212560; ***Euphorbia macroclada*** Boiss., Iran, *Y. Salmaki & S. Zarre 39896* (TUH), –, KC212562; ***Euphorbia mcvaughiana*** M.C.Johnst., Mexico, *M. Mayfield 2253* (KSC, MICH), KJ149609, KJ149658; ***Euphorbia maglicensis*** Rohlena, Bosnia, *B. Frajman & P. Schönswetter 12393* (IB), JN010074, KC212592; ***Euphorbia malleata*** Boiss., Iran, *Y. Salmaki & S. Zarre 39894* (TUH), –, KC212563; ***Euphorbia maresii*** subsp. ***balearica*** (Willk.) Malag., Spain, *B. Frajman 12545* (IB), JN010063, –; ***Euphorbia marschalliana*** Boiss., Iran, *Y. Salmaki 39929* (TUH), JF732971, –; ***Euphorbia masirahensis*** Ghaz., Oman, *J. Morawetz 348* (MICH), HQ900623, KC212431; ***Euphorbia matritensis*** Boiss., Spain, *J. Molero & al. 11/2008* (BCN), KC212299, KC212564; ***Euphorbia mauritanica*** L., South Africa, *J. Morawetz 277* (MICH), JN250189, JN249178; ***Euphorbia mazandaranica*** Pahlevani, Iran, *A. Pahlevani & M. Eskandari 55150* (IRAN), KC212304, KC212567; ***Euphorbia medicaginea*** Boiss., Spain, *J. Molero & A. Rovira 22/2007* (BCN), KC212305, –; ***Euphorbia megalatlantica*** Ball, Morocco, *R. Romo & R. Vilatersana 12511 (13949)* (IB), JN010066, KC212569; ***Euphorbia mellifera*** Aiton, Spain, *J. Molero 12/2007* (BCN), KC212306, KC212570; ***Euphorbia micractina*** Boiss., China, *W. Jin 008* (MICH), KC212308, KC212571; ***Euphorbia microcarpa*** (Prokh.) Krylov, Russia, *D. Geltman 144* (LE), KC212310, –; ***Euphorbia microsciadia*** Boiss., Iran, *Y. Salmaki & al. 39935* (TUH), KC212312, KC212572; ***Euphorbia microsphaera*** Boiss., Iran, *S. Safavi & Dezfulian 17840* (IRAN), KC212313, –; ***Euphorbia milii*** Des Moul., cult., *P. Berry 7826* (MICH), JN250191, JN249180; ***Euphorbia minuta*** Loscos & Pardo, Spain, *R. Riina 1872* (MA), KC212314, KC212573; ***Euphorbia mongoliensis*** M. H. Li & C. H. Zhang, China, *M. H. Li & C. H. Zhang 150602200520001LY* (PE), MW004167, –; ***Euphorbia monostyla*** Prokh., Iran, *Pahlevani 53763* (IRAN), LN680645, LN680654; ***Euphorbia montenegrina*** (Bald.) K. Malý, Montenegro, *M. Niketic & al. 11898* (IB), JN010068, KC212574; ***Euphorbia myrsinites*** L., Bosnia, *B. Frajman 12040* (IB), JN010069, KC212576; ***Euphorbia nakaii*** Hurus., South Korea, *Chung 120*, EU659751, –; ***Euphorbia natalensis*** Bernh. ex Krauss, South Africa, *G. Germishuizen 866* (PRE), KC212316, KC212577; ***Euphorbia neilmulleri*** M.C. Johnst., Mexico, *J. Henrickson 22475b* (IEB), –, KC212578; ***Euphorbia nereidum*** Jahand. & Maire, Morocco, *R. Riina 1778* (MICH), JN250198, JN249186; ***Euphorbia nesomii*** M. Mayfield, Mexico, *M. Mayfield 1905* (KSC, MICH), KJ149610, KJ149662; ***Euphorbia nevadensis*** subsp. ***bolosii*** Molero & Rovira, Spain, *J. Molero BCN53349* (MICH), KC212319, KC212581; ***Euphorbia nicaeensis*** All., Spain, *R. Riina 1767* (MICH), JN250200, JN249188; ***Euphorbia niciciana*** Borbás ex Novák, Serbia, *B. Frajman & P. Schönswetter 11242* (IB), JN010071, KC212584; ***Euphorbia normannii*** Schmalh. Ex Lipsky, Russia, *D. Geltman 235* (LE), MT957497, –; ***Euphorbia nubica*** N.E. Br., Kenya, *J. Morawetz 390* (MICH), KC212322, KC212585; ***Euphorbia obesa*** Hook.f., cult., *K. Wurdack D539* (US), JN250202, JN249189; ***Euphorbia oblongata*** Griseb., U.S.A. (introduced), *R. Riina 1839* (MICH), KC212325, KC212587; ***Euphorbia oblongifolia*** (K. Koch) K. Koch,

Azerbaijan, *G. Schneeweiß & A. Tribsch 6744* (WU), JN010072, KC212588; ***Euphorbia octoradiata*** H. Lév. & Vaniot, South Korea, *Chung 200*, EU659750, –; ***Euphorbia orientalis*** L., cult., *BRUS 1981–12301* (K), EU659764, –; ***Euphorbia orizabae*** Boiss., Mexico, *W. Graham & M. Frohlich 1020* (MICH), KJ149612, KJ149664; ***Euphorbia orphanidis*** Boiss., Greece, *B. Frajman & F. Faltner 16425* (IB), ON908644, –; ***Euphorbia orthoclada*** Baker, cult., *R. Riina 1739* (MA), JN250204, JN249191; ***Euphorbia orjeni*** Beck, Montenegro, *B. Frajman 15789* (IB), MN954394, –; ***Euphorbia osyridea*** Boiss. **1,** Iran, *Iranshahr 18136* (IRAN), –, LN998111; ***Euphorbia osyridea*** Boiss. **2,** Iran, *Pahlevani 1*[a], LN998101, –; ***Euphorbia ouachitana*** M. Mayfield, U.S.A., M. Mayfield 3549 (KSC, MICH), KJ149618, KJ149672; ***Euphorbia oxyphylla*** Boiss., Spain, *J. Molero & al. BCN53036* (BCN), KC212327, KC212590; ***Euphorbia pachyrrhiza*** Kar. & Kir., Kazakhstan, *S. Smirnov & al. s.n.* (LE), KC212328, –; ***Euphorbia pachysantha*** Baill., cult., *W. Rauh 73350* (HEID), JN250206, JN249193; ***Euphorbia palustris*** L., Austria, *B. Frajman & P. Schönswetter 11099* (IB), JN010073, KC212591; ***Euphorbia pamirica*** (Prokh.) Prokh., Tajikistan, *B. Frajman & P. Schoenswetter 14867* (IB), ON511366, –; ***Euphorbia paniculata*** subsp. ***paniculata*** Desf., Spain, *J. Molero & al. 16/2008 (duplicate)* (BCN), KC212330, KC212594; ***Euphorbia pannonica*** Host, Czech Republic, *P. Efimov & M. Heida 18* (LE), GU984331, –; ***Euphorbia papilionum*** S. Carter, Somalia, *M. Thulin & al. 9415* (K), HQ900640, –; ***Euphorbia paralias*** L., Greece, *R. Riina 1565* (MICH), JN250207, JN249194; ***Euphorbia pedroi*** Molero & Rovira, Portugal, *R. Riina 1585* (MICH), KC212332, KC212596; ***Euphorbia pekinensis*** Rupr., China, *no collector*, AY594263, –; ***Euphorbia peplidion*** Engelm., U.S.A., *M. Mayfield 2119* (TEX), KJ149619, KJ149673; ***Euphorbia peplus*** **var.** ***peplus*** L. **1,** Spain, *L. Barres 16* (BC), HQ900643, –; ***Euphorbia peplus*** **var.** ***peplus*** L. **2,** France, *P. Berry 7915* (MICH), –, KC212598; ***Euphorbia petrophila*** C.A. Mey., Russia, *D. Geltman s.n.* (LE), GU984332, –; ***Euphorbia phymatosperma*** Boiss. & Gaill., Iran, *S. Zarre & al. 41004* (TUH), KC212336, KC212599; ***Euphorbia pilosa*** L., Russia, *P. Berry 8007* (MICH), KC212339, –; ***Euphorbia pinea*** L., France, *B. Frajman & P. Schönswetter 12231* (IB), JN010078, KC212600; ***Euphorbia pinkavana*** M.C. Johnst., Mexico, *J. Henrickson 16008* (IEB), –, KC212601; ***Euphorbia piscatoria*** Aiton, cult., *no collector* (BC), HQ900644, –; ***Euphorbia pithyusa*** L., Spain, *P. Benedi & N. Montes s.n.* (BC), KC212340, KC212602; ***Euphorbia platyphyllos*** L., Spain, *J. Molero s.n.* (BCN), KC212341, KC212603; ***Euphorbia plebeia*** Boiss., Iran, *S. Zarre & al. 39982* (TUH), JF732978, –; ***Euphorbia polycaula*** Boiss. & Hohen., Iran, *S. Zarre & al. 39933* (TUH), KC212342, KC212604; ***Euphorbia polychroma*** A. Kern., Serbia, *B. Frajman & P. Schönswetter 11362* (IB), JN010083, KC212605; ***Euphorbia polygalifolia*** Boiss. & Reut., Spain, *J. Molero & A. Rovira BCN53606* (MICH), KC212343, KC212606; ***Euphorbia portlandica*** L., France, *P. Berry 7914* (MICH), KC212346, KC212610; ***Euphorbia procera*** M. Bieb., Russia, *D. Geltman 12* (LE), GU979433, –; ***Euphorbia prolifera*** Buch.-Ham. ex D. Don, China, *W. Jin 022* (MICH), KC212347, –; ***Euphorbia pterococca*** Brot., Spain, *J. Molero & A. Rovira 20/2007 (duplicate)* (MICH), KC212348, KC212611; ***Euphorbia pulcherrima*** Willd. ex Klotzsch, cult., *K. Wurdack D084* (US), JN250220, JN249207; ***Euphorbia purpurea*** (Raf.) Fernald, U.S.A., *J. Morawetz 315* (MICH), KC212349, KC212612; ***Euphorbia pyrenaica*** Jord., Spain, *J. Molero & A. Rovira BCN53619* (BCN), KC212356, KC212614; ***Euphorbia rapulum*** Kar. & Kir., Kazakhstan, *S. Smirnov s.n.* (LE), KC212357, KC212616; ***Euphorbia regis-jubae*** Webb & Berthel., Morocco, *R. Riina 1804* (MICH), JN250226, JN249213; ***Euphorbia resinifera*** O. Berg, cult., *P. Berry 7817* (MICH), JN250227, JN249214; ***Euphorbia retusa*** Forssk., Egypt, *K. Khalik 2672* (WAG), JN250228, JN249215; ***Euphorbia rhabdotosperma*** Radcl.-Sm., Georgia, *Y. Roskov 793* (LE), KC212360, KC212618;

***Euphorbia rigida*** M. Bieb., Greece, *W. Gutermann 28856* (Herb. Gutermann), JN010087, KC212619; ***Euphorbia rimarum*** Coss. & Balansa, Morocco, *R. Riina 1774* (MICH), JN250230, JN249217; ***Euphorbia roemeriana*** Scheele, U.S.A., *M. Mayfield 2158* (TEX), KJ149620, KJ149675; ***Euphorbia rothiana*** Spreng., India, *P. Bruyns 11474* (BOL), KC212363, –; ***Euphorbia rupestris*** Ledeb., Russia, *P. Berry 7996* (MICH), KC212364, KC212620; ***Euphorbia sahendi*** Bornm., Iran, *Y. Salmaki & al. 39891* (TUH), KC212365, KC212621; ***Euphorbia salicifolia*** Host, Germany, *F. Schuhwerk 06* (M), KC212366, KC212623; ***Euphorbia sarawschanica*** Regel, Kyrgyzstan, *G. Lazkov s.n.* (LE), GU979438, –; ***Euphorbia saxatilis*** Jacq., Austria, *B. Frajman & P. Schönswetter 12074* (IB), JN010092, –; ***Euphorbia scatorhiza*** S. Carter, cult., *V. Steinmann 1441* (RSA), AF537420, AF538181; ***Euphorbia schimperi*** C. Presl, Oman, *J. Morawetz 332* (MICH), KC212367, KC212624; ***Euphorbia schimperiana*** Scheele, Ethiopia, *J. Aldasoro 10398* (BCN), KC212373, KC212629; ***Euphorbia schizoloba*** Engelm., U.S.A., *M. Mayfield 3567* (KSC, MICH), KJ149621, KJ149677; ***Euphorbia schugnanica*** B. Fedtsch., Tajikistan, *A. Konnov 266* (MO), JN250235, –; ***Euphorbia sclerocyathium*** Korovin & Popov, Turkmenistan, *D. Kurbanov 705* (MO), JN250237, JN249224; ***Euphorbia sclerophylla*** Boiss., South Africa, *H. Furness 61* (PRE), KC212375, KC212630; ***Euphorbia segetalis*** var. ***segetalis*** L., Portugal, *R. Riina 1547* (duplicate) (MICH), JN250238, JN249225; ***Euphorbia seguieriana*** Neck., France, *J. Molero 237/09* (MICH), KC212377, KC212631; ***Euphorbia serpentini*** Novák, Serbia, *N. Kuzmanović 15157* (IB), MT957520, –; ***Euphorbia serrata*** L., Spain, *R. Riina 1608* (MA), KC212378, KC212633; ***Euphorbia sieboldiana*** C. Morren & Decne., South Korea, *Tho & J.H. Kim 2001–0001*, EU659753, –; ***Euphorbia smithii*** S. Carter, Oman, *J. Morawetz 336* (MICH), JN250240, JN249227; ***Euphorbia sororia*** Schrenk, Iran, Mirtadzadini St 331-2 (IRAN), OQ230321, –; ***Euphorbia* sp.**, China, *W. Jin 015* (MICH), KC212384, KC212637; ***Euphorbia spartiformis*** Mobayen, Iran, *A. Pahlevani, M. Eskandari & Bahramishad 55156* (IRAN), KC212379, –; ***Euphorbia spathulata*** Lam., U.S.A., *G. Rink 5861* (NY), JN250242, JN249229; ***Euphorbia spinidens*** Bornm. ex Prokh., Iran, *Y. Salmaki & S. Zarre 38156* (TUH), –, KC212639; ***Euphorbia spinosa*** L., Switzerland, *R. Riina 1842* (MA), KC212386, KC212641; ***Euphorbia squamigera*** Loisel., Spain, *R. Riina 1760* (MA), KC212387, KC212642; ***Euphorbia squamosa*** Willd., Russia, *D. Geltman 33* (LE), GU937804, –; ***Euphorbia stepposa*** Zoz, Russia, *M. Wernisch 12477* (WHB), JN010103, KC212643; ***Euphorbia stolonifera*** Marloth ex A.C. White, R.A. Dyer & B. Sloane, South Africa, *R. Becker 1109* (MICH), KC212388, KC212644; ***Euphorbia stracheyi*** Boiss., China, *N. Yang 0937* (KUN), KC212391, KC212647; ***Euphorbia striata*** Thunb., South Africa, *N. Kroon 11703* (PRE), KC212394, –; ***Euphorbia stricta*** L., Bosnia, *B. Frajman & P. Schönswetter 12398* (IB), JN010104, KC212649; ***Euphorbia stygiana*** H.C. Watson, Portugal, *J. Molero & al. BCN 86911* (BCN), KC212397, KC212651; ***Euphorbia subamplexicaulis*** Kar. & Kir., Kazakhstan, *S. Smirnov & al. s.n* (LE), KC212398, –; ***Euphorbia subcordata*** Ledeb., Russia, *Shmakov & al. s.n.* (LE), KC212399, –; ***Euphorbia subhastata*** Vis. & Pancic, Serbia, *S. Kurovi. & S. Vukojiki 15831* (IB), ON511373, –; ***Euphorbia subtilis*** (Prokh.) Prokh., Russia, *N. Tzvelev 76* (LE), GU984321, –; ***Euphorbia sulcata*** Lens ex Loisel., Spain, *R. Riina 1861* (MA), KC212400, KC212652; ***Euphorbia sulphurea*** Pahlevani, Iran, *Pahlevani & Bahramishad 55151 (IRAN)*, LN680646, LN680655; ***Euphorbia sultan-hassei*** Å. Strid & al., Greece, *R. Riina 1568* (MICH), KC212401, KC212653; ***Euphorbia sylvicola*** Pahlevani & Frajman, Iran, *A. Pahlevani & Torabi 76622* (IRAN), ON908669, –; ***Euphorbia talastavica*** (Prokh.) Prokh., Kazakhstan, *D. Geltman 206* (LE), KC212405, KC212657; ***Euphorbia taurinensis*** All., Ukraine, *V. Byalt & al. 364* (LE), GU984337, –; ***Euphorbia teheranica*** Boiss., Iran, *Y. Salmaki & S. Zarre 39917* (TUH), KC212406,

KC212658; ***Euphorbia terracina*** L., South Africa, *R. Becker 1528* (MICH), KC212408, KC212660; ***Euphorbia tetrapora*** Engelm., U.S.A., *M. Mayfield 2684* (LSU), KJ149622, KJ149678; ***Euphorbia texana*** Boiss., U.S.A., *B. Tharp s.n.* (MICH), KC212409, KC212661; ***Euphorbia thessala*** (Formánek) Degen & Dörfl., Serbia, *B. Frajman & M. Turjak* 11981 (IB), MF521402, –; ***Euphorbia thompsonii*** Holmboe, Cyprus, *B. Frajman & P. Schönswetter 12736* (IB), ON908675, –; ***Euphorbia tibetica*** Boiss., Kyrgyzstan, *G. Lazkov s.n.* (LE), KC212411, –; ***Euphorbia tirucalli*** L., cult., *B. van Ee 741* (US), JN250247, JN249233; ***Euphorbia tommasiniana*** Bertol., Italy, *B. Frajman 16945* (IB), ON511375, –; ***Euphorbia transoxana*** (Prokh.) Prokh., Tajikistan, *R. Kamelin 1717* (LE), GU979425, –; ***Euphorbia transtagana*** Boiss., Portugal, *J. Molero & al. 13/2008* (BCN), KC212412, KC212662; ***Euphorbia trichotoma*** Kunth, Belize, *S. Hill 20357* (MO), AF537534, –; ***Euphorbia triflora*** Schott, Nyman & Kotschy, Slovenia, *B. Frajman & P. Schönswetter 11131* (IB), JN010107, KC212663; ***Euphorbia tshuiensis*** (Prokh.) Serg. ex Krylov, Russia, *A. Tribsch & F. Essl 10284* (WU), JN010108, KC212664; ***Euphorbia tuckeyana*** Steud. ex Webb, Cape Verde, *J. Molero & A. Rovira BCN58758* (BCN), KC212415, KC212667; ***Euphorbia tuerckheimii*** Urb., The Dominican Republic, *T. Clase & al. 7332* (MICH), –, KJ149680; ***Euphorbia turczaninowii*** Kar. & Kir., Iran, *Y. Salmaki & al. 38203* (TUH), KC212416, KC212668; ***Euphorbia turkestanica*** Regel, Turkmenistan, *G. Semidel & L. Kuzamina s.n.* (LE), KC212417, –; ***Euphorbia uliginosa*** Welw. ex Boiss., Spain, *J. Molero & A. Rovira BCN53605* (BCN), KC212418, KC212669; ***Euphorbia uralensis*** Fisch. ex Link, Russia, *D. Geltman 28* (LE), GU984315, –; ***Euphorbia usambarica*** Pax, Kenya, *J. Morawetz 401* (MICH), KC212419, KC212670; ***Euphorbia valdevillosocarpa*** Arvat & Nyár., Moldova, *I. Zhilkina s.n.* (LE), GU979431, –; ***Euphorbia valerianifolia*** Lam., Greece, *W. Gutermann 34627* (Herb. Gutermann), JN010109, KC212671; ***Euphorbia valliniana*** Belli, Italy, *B. Frajman & P. Schönswetter 12224* (IB), JN010110, KC212672; ***Euphorbia variabilis*** Ces., Italy, *G. Schneeweiß, P. Schönswetter & A. Tribsch 3933* (WU), JN010111, KC212673; ***Euphorbia velenovskyi*** Bornm., Bulgaria, *B. Frajman & P. Schönswetter 11345* (IB), JN010113, KC212674; ***Euphorbia veneris*** M.S. Khan, Cyprus, *M. Galbany & al. 2039* (BC), HQ900667, KC212675; ***Euphorbia verrucosa*** L., Slovenia, *B. Frajman & P. Schönswetter 11771* (IB), JN010115, KC212676; ***Euphorbia virgata*** Waldst. & Kit., Austria, *F. Schuhwerk 6/237* (M), KC212421, KC212679; ***Euphorbia wallichii*** Hook.f., China, *W. Jin 009* (MICH), KC212426, KC212683; ***Euphorbia yaquiana*** (Cockerell) Tidestr., U.S.A., *M. Mayfield 3364* (KSC, MICH), KJ149624, KJ149681; ***Euphorbia yaroslavii*** Poljakov, Kazakhstan, *A. Urazova s.n.* (AA), KC212429, –; ***Euphorbia xiangxiui*** N. Wei, Q. Yu, G. X. Chen & Q. F. Wang, China, *Q. Yu LX20051201* (HIB!), –, MZ494740; ***Neoguillauminia cleopatra*** (Baill.) Croizat, New Caledonia, *K. Cameron 2015* (NY), JN250099, JN249090.
