## Supplementary File 2 for "The conquest and diversification of leafy spurges across the Holarctic and beyond: biogeography and evolution of life-history of *Euphorbia* subgenus *Esula*"

**Supplementary File 2.** Detailed explanation of the biogeographic analyses carried out in the study.

Area definition was based on several criteria, including the current distribution of species of subg. *Esula* (GBIF, 2023; POWO, 2023; Riina & Berry, 2023), the paleogeographic history of Africa, America, and Eurasia (Sanmartín et al., 2001), and maximising congruence with previous studies (Meseguer et al., 2015; Jin et al., 2020). The five delimited areas were (Fig. 1): Africa (AFR), Eastern Palearctic (EP), Macaronesian region (MAC), North America, Central America, and Caribbean region (NAM), and Western Palearctic (WP). Widespread ancestral areas were limited to combinations of three areas, which is the maximum range of extant taxa. We use the native distribution of the extant species, i.e. discarding potential invasive ranges or human introductions (Meseguer et al., 2013). Four species of *Euphorbia* are distributed outside our predefined biogeographic areas: one in Chile and three in the Indo-Pacific region (Fig. 1, Table S1). Of these, only *E. glauca* G.Forst. (New Zealand) was included in our phylogeny. Since the number of ancestral states in the Dispersal–Extinction–Cladogenesis (DEC) model increases exponentially with the number of areas (Landis et al., 2018), we pruned *E. glauca* from the tree to avoid adding a sixth area to the analysis. It is unlikely that including this single species would have a major effect on the inference of biogeographic rates and ancestral ranges.

We inferred ancestral ranges and rates of range evolution under the DEC model (Ree & Smith, 2008) and using Landis et al.'s (2018) Bayesian implementation in RevBayes v.1.1.1 (Höhna et al., 2016a). We modelled anagenetic events as two free parameters, range expansion (dispersal) and range contraction (local extinction), using default priors. Cladogenetic events were modelled as a simplex, with equal probability for all range inheritance scenarios involving sympatric ('s') or allopatric ('a') events. We did not incorporate the cladogenetic founder speciation scenario modelled by the 'j' parameter (Matzke, 2016) due to observed biases in biogeographic inference associated with this model (Ree & Sanmartín, 2018; Sanmartín, 2021). All DEC-derived models assume that the observed phylogeny represents all speciation events that ever existed: i.e. inference is conditioned on the reconstructed tree (Cornuault & Sanmartín, 2022). Therefore, "hidden" cladogenetic events such as unsampled or extinct lineages are not accounted for in the estimation of biogeographic rates and ancestral ranges. The bias introduced by these missing cladogenetic events is larger under DEC models with the 'j' parameter (DEC+J, DIVA+J) because these models allow instantaneous dispersal to a new area to occur at the speciation node; conversely, the original DEC model treats dispersal as a branch-evolving process that is only dependent on time. The DEC analysis was run on the MCC tree with one tip per species, assigning each tip the species' entire native range. We used the *drop.tip* function of the package *ape* (Paradis & Schliep, 2019) in R (R Development Core Team, 2017) to prune the additional tips and the outgroup taxa from the tree; *E. glauca* was also excluded (see above). To inform the inference of ancestral regions, we retained one tip for each of the other three *Euphorbia* subgenera, coding them with the entire subgeneric range. This approach allowed for the consideration of biogeographic evolution before the MRCA of subg. *Esula*. We ran two independent analyses using the default priors described in the RevBayes online tutorial (revbayes.github.io/tutorials/biogeo/biogeo_simple.html). Two independent analyses were run using the default priors for 10,000 generations, sampling every 10th generation, and results were summarised in the MCC tree. The first analysis (M0) was conducted with the rate of the baseline range evolution parameter ('rate_bg') modelled as constant through time and using the simplex prior for cladogenetic events. The second analysis (M1) was run with cladogenetic events modelled with a Dirichlet distribution prior, in which the probability of allopatry "a" was equal to 1 minus the probability of sympatry "s". Additionally, the M1 analysis included a time-stratified dispersal model, in which the baseline rate was rescaled according to geological connectivity scenarios, which in turn reflected the shifting probability of migration over time. We used the following criteria to rescale the baseline dispersal rate: migration between areas sharing an edge or those connected by land corridors was not downscaled, i.e. rates were assigned a scalar of 1; migration between neighbouring areas not connected by land were downscaled by a factor of 0.5; migration between areas separated by large ocean barriers or between non-adjacent plates through an intermediate area (e.g., between AFR and EP through WP) was downscaled by a factor of 0.1. We divided the phylogenetic tree into four time slices (TS1–TS4) to capture important periods of tectonic rearrangements and/or climatic change (Table 1; Buerki et al., 2011). TS1: Late Paleocene-Late Oligocene within the Paleogene period, characterised by warm-tropical or temperate climates, including the TEE and LOWE; TS2: Early Miocene-Late Miocene within the Neogene period, including the MMCO warming event and ending with the start of a global cooling trend; TS3: the Late Neogene period, when global climates cooled significantly following the LMCE, but including the MSC and ending on a global warming event, the MPWE; TS4: the period comprising the Plio-Pleistocene and Quaternary periods, dominated by the glaciations and interglacial cycles of the Pleistocene and ending in the Holocene. Instead of using hard boundaries between the four TS, we implemented an approach similar to that used by Landis et al. (2018) for islands and by Masa-Iranzo et al. (2021) for continental scenarios. We applied a uniform distribution with upper and lower bounds, representing a time interval when significant climatic and tectonic events might have influenced migration rates between biogeographic regions. During the analysis, the MCMC algorithm samples a time point within this uniform distribution at which the model switches between the dispersal rate matrices of two different TS. The rationale is to include uncertainty regarding the specific climatic or tectonic event that influenced the dispersal rate in the lineage. For example, the transition between TS1 and TS2 was bounded as the time interval between the TEE and TTE, since these two events had a profound influence on the evolution of the temperate mixed-mesophytic and Madrean-Tethyan vegetation (Table 1). The interval between TS2 and TS3 was bounded by the LMCE and the MPWE, a period during which important changes in vegetation occurred (Schneck et al., 2012). The transition between TS3 and TS4 spans the period from the opening of the Bering Strait and the start of the glacial cycles of the Plio-Pleistocene (Table 1). We summarised nodal ancestral states as marginal PP onto the maximum a posteriori (MAP) tree using the code provided in the RevBayes website (revbayes.github.io/tutorials/morph_ase/scm_hrm.html). We also estimated the number and timing of dispersal and extinction events along the internal and terminal branches using a heuristic approximation to stochastic character mapping that does not require a rejection sampling step (Freyman & Höhna, 2019). Events of biogeographic change were summarised on the MAP tree using 500 TS. We visualised the results using R scripts and the R packages *phytools* (Revell, 2012) and *RevGadgets* v.1.0 (Tribble et al., 2022). Finally, marginal likelihood values were estimated for the two alternative biogeographic models, M0 and M1, and compared via BF based on PS and SS power posteriors, with 100 steps each. Appendix S1 provides the scripts to carry out these analyses.
