## Supplementary File 3 for "The conquest and diversification of leafy spurges across the Holarctic and beyond: biogeography and evolution of life-history of *Euphorbia* subgenus *Esula*"

**Supplementary File 3.** Detailed explanation of the diversification analyses carried out in the study.

All diversification analyses were run on the MCC tree from the BEAST analysis, pruned to remove the outgroup species and retain the 332 species of subg. *Esula*. We first plotted the lineage-through-time (LTT) plot to visually inspect the accumulation of lineages over time, using the *ltt.plot* function in the R package *ape* v.5.5 (Paradis & Schliep, 2019); a random sample of 1,000 chronograms from the Bayesian MCMC posterior distribution was also plotted to reflect uncertainty in divergence time estimates in the MCC tree. Next, we investigated whether speciation and extinction rates have been constant over the evolutionary history of subg. *Esula*, or if the subgenus has undergone episodic rate shifts during which the diversification rate (speciation minus extinction) and the relative extinction rate (ratio of extinction to speciation) increased or decreased, simultaneously, across all lineages. We used the Compound Poisson Process of Mass Extinction model (CoMET; May et al., 2016) implemented in the R package *TESS* (Höhna et al., 2016b) to estimate shifts in speciation and extinction rates at discrete points in time using a reversible jump MCMC algorithm and BF comparisons. We also used CoMET to estimate the magnitude and timing of mass extinction events (MEE), defined as discrete points in time when a significant percentage of the standing diversity (95–70%) is removed across all lineages. The joint estimation of shifts in speciation and extinction rates and the magnitude of the MEE, measured as the probability of survival for all lineages (ρ), is considered as statistically non-identifiable (Stadler, 2011). To solve this, CoMET uses a hierarchical Bayesian approach in which MEE are estimated after accounting for the probability of shifts in the speciation and extinction rates, considered as nuisance parameters (May et al., 2016). We ran two MCMC chains of 1 million iterations with a sampling frequency of 100 and ensured a minimum ESS of 1,000. We determined the shape of the prior distributions for speciation and extinction rates (normal) through empirical estimation from the data using the 'empiricalHyperPriors=TRUE' argument. We relied on default values in *TESS* for the initial speciation rate (2.0), initial extinction rate (1.0), and the expected number of rate changes and MEE (2). We set the sampling fraction (ρ) at present to 0.66, i.e. the ratio between the number of sampled species in the phylogeny and the total extant species diversity in subg. *Esula* (321/490). Two different strategies were implemented to model incomplete taxon sampling in our phylogeny: uniform sampling, which assumes that every species at present has the same probability ρ of being included in the sample; diversified sampling, which assumes that species in the phylogeny were selected to maximise systematic diversity and that the most recent speciation events were discarded (Höhna et al., 2016b). To assess the mixing and convergence of the two chains, we employed MCMC diagnostics in *TESS* (Höhna, 2013), including the Rubin-Gelman statistic and ensuring that ESS values reached a value > 500.

We further explored time-varying birth-death scenarios using the Horseshoe Markov random field (HSMRF) episodic birth-death (EBD) model implemented in RevBayes (revbayes.github.io/tutorials/divrate/ebd.html). In this model, rates of speciation and extinction are autocorrelated across time intervals, so that the mean of these rates is centred around the rate of the previous time interval, thus avoiding "abrupt" changes. The use of a horseshoe estimator for sparse signals as the prior distribution allows modelling large jumps between prefixed time intervals separated by relatively constant rates within time intervals (Magee et al., 2020). We followed the tutorial (revbayes.github.io/tutorials/divrate/ebd.html), dividing the phylogeny into 10 time intervals and using a uniform taxon sampling strategy with a ρ value of 0.66. An MCMC Bayesian analysis was run for $10^7$ generations, and the results visualised in *RevGadgets* as the marginal probabilities for speciation and extinction rates and the timing and magnitude of shift events. Appendix S2 in Data availability provides the script for the CoMET and HSMRF-EBD analyses.

The aforementioned EBD models assume that diversification rates are equal across all lineages at any given time point. To implement a birth-death process where diversification rates vary across branches, we analysed the pruned MCC tree under the lineage-specific birth-death (LSBDS) model implemented in RevBayes (Höhna et al., 2019). LSBDS assumes a multistate state-dependent speciation extinction model (Maddison et al., 2007), in which each state has its own speciation and extinction rates, and where shifts in states (and hence in diversification rates) are modelled dynamically. Numerical integration is used to compute the likelihood of multiple events in an instant of time, including the probability of no shifts and shifts occurring at unsampled or extinct lineages. We followed the tutorial (revbayes.github.io/tutorials/divrate/branch_specific.html), using four rate categories and three rate shifts in the speciation rate, while the extinction rate was kept constant to avoid problems with non-identifiability (Höhna et al., 2019). The MCMC chain was run for 15,000 generations, using the same ρ value as above and an exponential prior with a mean of 1/0.587405 for the SD (meaning the distributions' 95% intervals will cover one order of magnitude). We then obtained the branch-specific diversification rate estimates for each branch of the tree using the rejection-free stochastic rate mapping algorithm (Freyman & Höhna, 2019). Appendix S3 in Data availability provides the script to run the LSBDS analysis.

The LSBDS model supported the existence of branch-specific changes in speciation rates in subg. *Esula* (see Results below). To test whether the observed shifts could be causally linked to changes in life-history, we reanalysed our phylogeny under the Binary State-dependent Speciation and Extinction model (BiSSE; Maddison et al., 2007; FitzJohn, 2012), implemented in a Bayesian framework in RevBayes (revbayes.github.io/tutorials/sse/bisse.html). We coded life-history as two states: annual (0) or perennial (1) for all species of subg. *Esula*, including those not represented in our analysis (Table S1). We note that the definition of annuals used in our study includes those classified as “biennials” in the literature. We ran a single MCMC chain with 15,000 generations, sampling every 100th generation, and summarised ancestral states and state-shifts along branches in the MAP tree using stochastic free-rejection sampling with 500 time slices.

BiSSE assumes that all heterogeneity in diversification rates among clades in the phylogeny can be attributed to transitions in the focal character. However, rate heterogeneity could also be explained by unobserved 'hidden' characters that covary with the observed character across the phylogeny. To correct for the potential high Type I error rate in BiSSE (Rabosky & Goldberg, 2015), Beaulieu & O'Meara (2016) introduced the Hidden-State Speciation and Extinction model (HiSSE). In HiSSE, there are two 'hidden' states (A, B) within each state of the focal ("observed") character states (0, 1), and each composite 'observed-hidden state' (e.g. 0A, 1B) is modelled as having its own speciation and extinction rate. Transition rates between the hidden states are assumed to be equal, while transition rates between the two focal states can be asymmetric. Significant differences between the diversification rates of the focal states, regardless of the hidden state, are interpreted as support for a state-dependent diversification process. We ran the HiSSE analysis in RevBayes (revbayes.github.io/tutorials/sse/hisse.html), using lognormal priors for speciation and extinction rates centred in the extant diversity and exponential priors for asymmetric transition rates in the focal character. All other settings were similar to the BiSSE analysis. Since the RevBayes implementation of HiSSE does not apply state-specific ρ values, we additionally ran HiSSE in the R package *hisse* (Beaulieu et al., 2021), which allows the use of state-dependent sampling fractions. We used the function f = c(0.67, 0.66) to incorporate the percentages of annual and perennial species relative to the total diversity. Finally, we implemented in RevBayes a lineage-specific, character-independent rate-variable diversification model (CID; Caetano et al., 2018). This model allows birth-death rates to vary across branches or lineages in the tree, but unlike BiSSE and HiSSE, rate heterogeneity is not structured around an evolving focal trait. Specifically, we modelled speciation and extinction rates as different between the two hidden states A and B, but equal for the two states within the focal character, annual and perennial. All other settings were similar to those in the BiSSE analysis. We used BF comparisons in RevBayes to compare the BiSSE and HiSSE models against the CID model, considered here as the null model, i.e. no causal relationship between the evolution of a trait and diversification rates. The marginal likelihood value for each model was estimated through PS and SS sampling with 50 parallel power posterior analyses, each with a chain length of 500 generations. Appendix S4 in Data availability provides the scripts to run the BiSSE, HiSSE, and CID analyses.
