## Supplementary File 4 for "The conquest and diversification of leafy spurges across the Holarctic and beyond: biogeography and evolution of life-history of *Euphorbia* subgenus *Esula*"

**Supplementary File 4**. Maximum clade credibility tree for *Euphorbia* subg. *Esula* and related taxa, showing phylogenetic relationships and lineage divergence times inferred by BEAST.

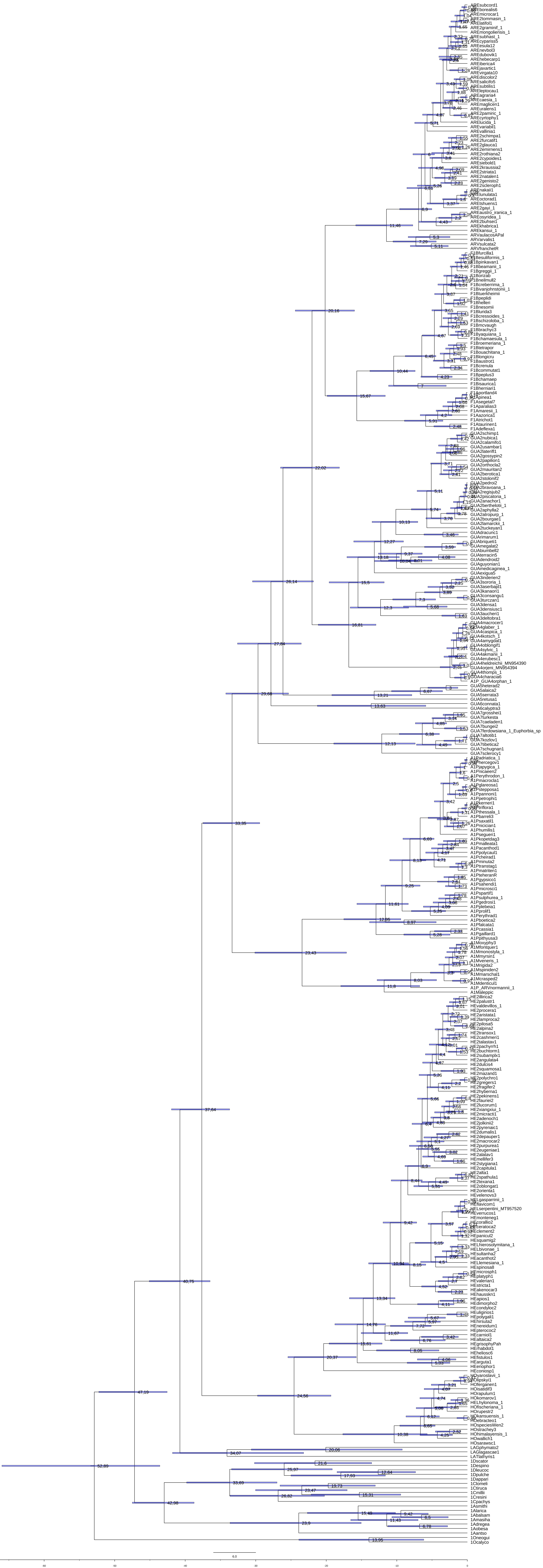

6.0
-60
-50
-40
-30
-20
-10
0
