## Supplementary Figure 1 for "The conquest and diversification of leafy spurges across the Holarctic and beyond: biogeography and evolution of life-history of *Euphorbia* subgenus *Esula*"

**Fig. S1**. Majority-rule consensus tree of the MrBayes analysis based on the complete concatenated ITS and *ndhF* sequences of *Euphorbia* subg. *Esula* and closest allies. The numbers represent support values with Bayesian inference posterior probability.

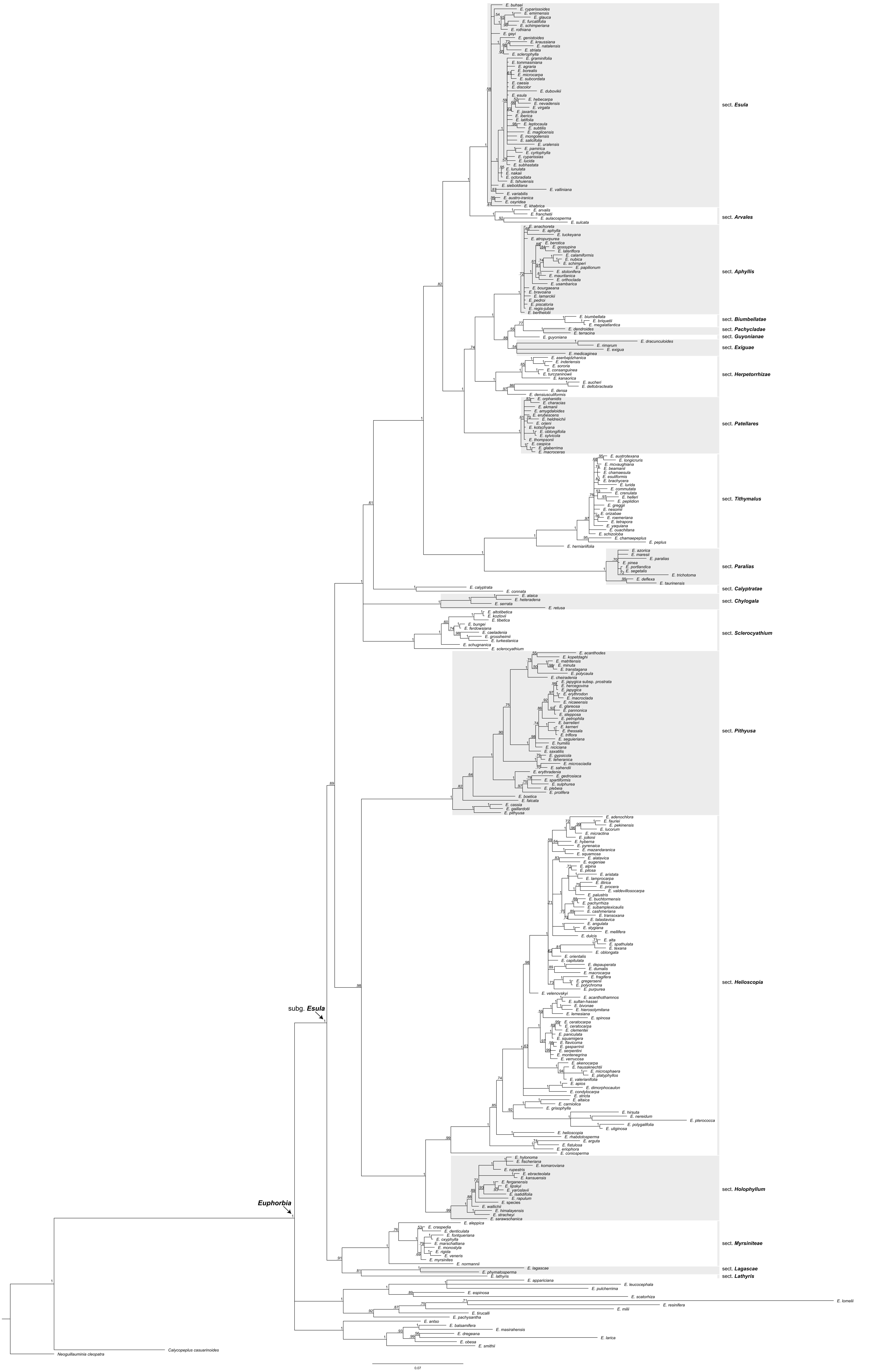

sect. Esula
sect. Arvales
sect. Aphyllis
sect. Biumbellatae
sect. Pachycladae
sect. Guyonianae
sect. Exiguae
sect. Herpetorrhizae
sect. Patellares
sect. Tithymalus
sect. Paralias
sect. Calyptratae
sect. Chylogala
sect. Sclerocyathium
sect. Pithyusa
sect. Helioscopia
sect. Holophyllum
sect. Myrsiniteae
sect. Lagascae
sect. Lathyris
subg. Esula
Euphorbia
Calycopeplus casuarinoides
Neoguillauminia cleopatra
0.07
