## Supplementary Figure 2 for "The conquest and diversification of leafy spurges across the Holarctic and beyond: biogeography and evolution of life-history of *Euphorbia* subgenus *Esula*"

**Fig. S2**. Majority-rule consensus tree of the MrBayes analysis based on the complete ITS sequences of *Euphorbia* subg. *Esula* and closest allies. The numbers represent support values with Bayesian inference posterior probability.

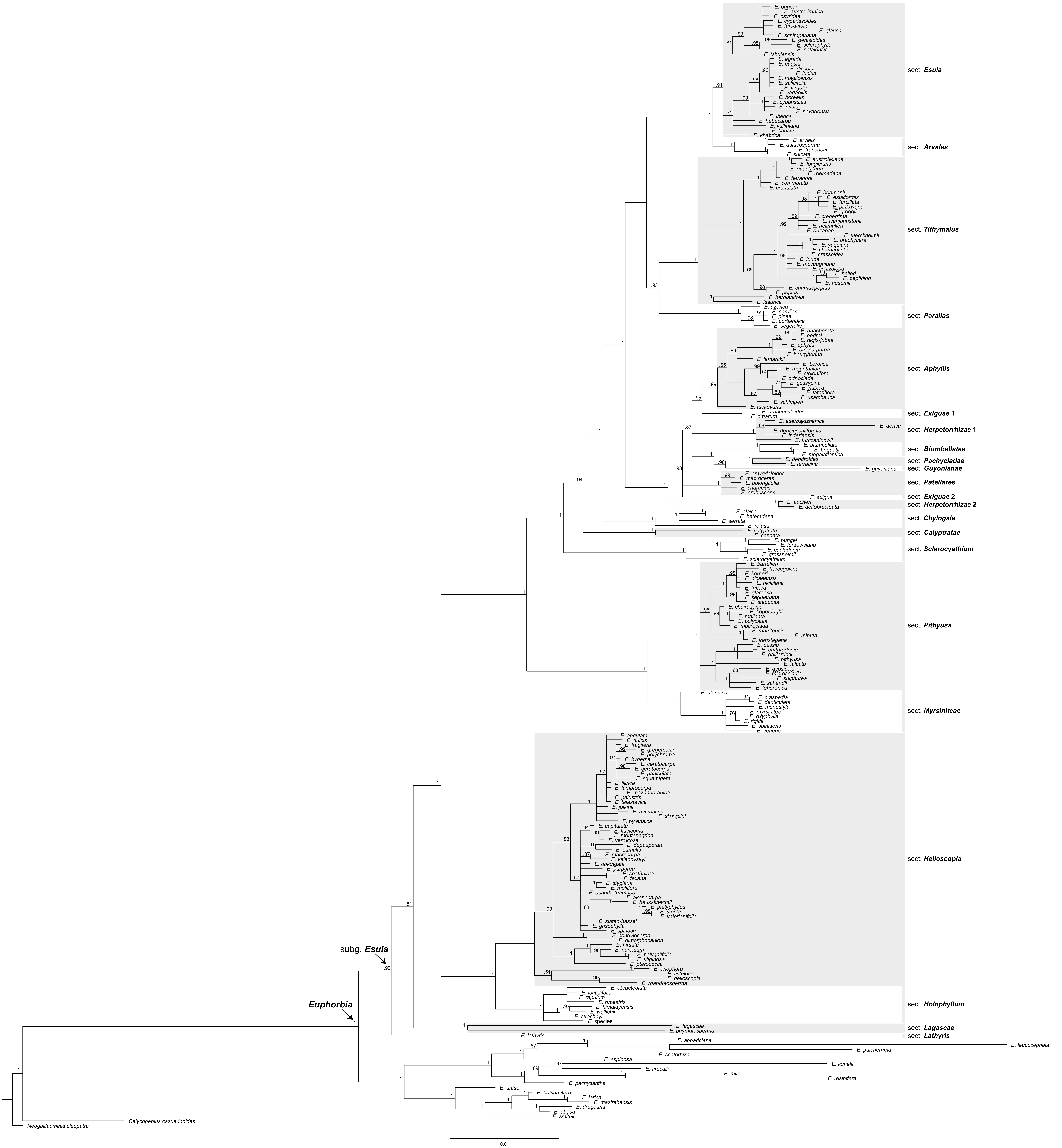
E. buhsei
E. austro-iranica
E. osyridea
E. cyparissoides
E. furcatifolia
E. glauca
E. schimperiana
E. genistoides
E. sclerophylla
E. natalensis
E. tshuiensis
E. agraria
E. caesia
E. discolor
E. lucida
E. maglicensis
E. salicifolia
E. virgata
E. variabilis
E. borealis
E. cyparissias
E. esula
E. nevadensis
E. iberica
E. hebecarpa
E. valliniana
E. kansui
E. khabrica
sect. Esula
E. arvalis
E. aulacosperma
E. franchetii
E. sulcata
sect. Arvales
E. austrotexana
E. longicruris
E. ouachitana
E. roemeriana
E. tetrapora
E. commutata
E. crenulata
E. beamanii
E. esuliformis
E. furcillata
E. pinkavana
E. greggii
E. creberrima
E. ivanjohnstonii
E. neilmulleri
E. orizabae
E. tuerckheimii
E. brachycera
E. yaquiana
E. chamaesula
E. cressoides
E. lurida
E. mcvaughiana
E. schizoloba
E. helleri
E. peplidion
E. nesomii
E. chamaepeplus
E. peplus
E. herniariifolia
E. isaurica
sect. Tithymalus
E. azorica
E. paralias
E. pinea
E. portlandica
E. segetalis
sect. Paralias
E. anachoreta
E. pedroi
E. regis-jubae
E. aphylla
E. atropurpurea
E. bourgaeana
E. lamarckii
E. berotica
E. mauritanica
E. stolonifera
E. orthoclada
E. gossypina
E. nubica
E. lateriflora
E. usambarica
E. schimperi
E. tuckeyana
sect. Aphyllis
E. dracunculoides
E. rimarum
sect. Exiguae 1
E. aserbajdzhanica
E. densa
E. densiusculiformis
E. inderiensis
E. turczaninowii
sect. Herpetorrhizae 1
E. biumbellata
E. briquetii
E. megalatlantica
sect. Biumbellatae
E. dendroides
E. terracina
sect. Pachycladae
E. guyoniana
sect. Guyonianae
E. amygdaloides
E. macroceras
E. oblongifolia
E. characias
E. erubescens
sect. Patellares
E. exigua
sect. Exiguae 2
E. aucheri
E. deltobracteata
sect. Herpetorrhizae 2
E. alaica
E. heteradena
E. serrata
sect. Chylogala
E. retusa
E. calyptrata
E. connata
sect. Calyptratae
E. bungei
E. ferdowsiana
E. caeladenia
E. grossheimii
E. sclerocyathium
sect. Sclerocyathium
E. barrelieri
E. hercegovina
E. kerneri
E. nicaeensis
E. niciciana
E. triflora
E. glareosa
E. seguieriana
E. stepposa
E. cheiradenia
E. kopetdaghi
E. malleata
E. polycaula
E. macroclada
E. matritensis
E. minuta
E. transtagana
E. cassia
E. erythradenia
E. gaillardotii
E. pithyusa
E. falcata
E. gypsicola
E. microsciadia
E. sulphurea
E. sahendii
E. teheranica
sect. Pithyusa
E. aleppica
E. craspedia
E. denticulata
E. monostyla
E. myrsinites
E. oxyphylla
E. rigida
E. spinidens
E. veneris
sect. Myrsiniteae
E. angulata
E. dulcis
E. fragifera
E. gregersenii
E. polychroma
E. hyberna
E. ceratocarpa
E. ceratocarpa
E. paniculata
E. squamigera
E. illirica
E. lamprocarpa
E. mazandaranica
E. palustris
E. talastavica
E. jolkinii
E. micractina
E. xiangxiui
E. pyrenaica
E. capitulata
E. flavicoma
E. montenegrina
E. verrucosa
E. depauperata
E. dumalis
E. macrocarpa
E. velenovskyi
E. oblongata
E. purpurea
E. spathulata
E. texana
E. stygiana
E. mellifera
E. acanthothamnos
E. akenocarpa
E. haussknechtii
E. platyphyllos
E. stricta
E. valerianifolia
E. sultan-hassei
E. grisophylla
E. spinosa
E. condylocarpa
E. dimorphocaulon
E. hirsuta
E. nereidum
E. polygalifolia
E. uliginosa
E. pterococca
E. eriophora
E. fistulosa
E. helioscopia
E. rhabdotosperma
sect. Helioscopia
E. ebracteolata
E. isatidifolia
E. rapulum
E. rupestris
E. himalayensis
E. wallichii
E. stracheyi
E. species
sect. Holophyllum
E. lagascae
E. phymatosperma
sect. Lagascae
E. lathyris
sect. Lathyris
subg. Esula
Euphorbia
E. appariciana
E. leucocephala
E. pulcherrima
E. scatorhiza
E. espinosa
E. lomelii
E. tirucalli
E. milii
E. resinifera
E. pachysantha
E. antso
E. balsamifera
E. larica
E. masirahensis
E. dregeana
E. obesa
E. smithii
Calycopeplus casuarinoides
Neoguillauminia cleopatra
0.01
