## Supplementary Figure 4 for "The conquest and diversification of leafy spurges across the Holarctic and beyond: biogeography and evolution of life-history of *Euphorbia* subgenus *Esula*"

**Fig. S4**. Reconstruction of the spatio-temporal evolution of *Euphorbia* subg. *Esula* and closest allies, using Bayesian dispersal extinction cladogenesis and model M0 (see Methods). The phylogenetic tree is the maximum clade credibility (MCC) tree based on concatenated ITS and *ndhF* sequences, pruned from the BEAST tree shown in Figs. 2G, 2H. As representatives of the other three *Euphorbia* subgenera we included one taxon per subgenus, representing the main distributional area of the respective subgenus. To reduce the size of the tree, those clades within *E.* subg. *Esula* with ages less than 2.5 Mya that had the same distribution were eliminated. **(A)** Biogeographic reconstruction of *E.* subg. *Esula*. The maximum number of areas allowed at one time was three. A pie chart on the node indicates alternative ancestral geographical ranges, with pie slices indicating their posterior probabilities (PP). Values below nodal pie charts indicate the PP for the ancestral range receiving the highest probability; only values < 0.90 are shown. Letters near nodal pie charts indicate the nodes mentioned in the text. Coloured nodes represent the geographical range state according to the legend. Colours of five major biogeographic areas correspond to the inset map. Coloured circles before the species names give present ranges. The smaller pie charts on the branches indicate the geographical range immediately after the speciation event; a pie on the node indicates how this range has changed as a result of range expansion and contraction events along the branch. The dashed vertical lines illustrate periods divided into decades. Acronyms above the global mean temperature curve, obtained from Zachos et al. (2001), indicate climatic events which may have played a role in the evolution of *E.* subg. *Esula*: LMCE, Late Miocene Cooling Event; LOWE, Late Oligocene Warming Event; MMCO, Mid-Miocene climatic optimum; MPWE, Mid-Pliocene Warming Event; TEE, Terminal Eocene Event. **(B)** MAP tree showing support values, in the form of marginal posterior probabilities, for the transition events between ancestral ranges reconstructed along branches in (C) using stochastic mapping; boxes on the left indicate marginal posterior probabilities for the inferences depicted as colours. **(C)** Maximum a posteriori (MAP) M0 reconstruction of the spatio-temporal evolution in *E.* subg. *Esula* simulated under Bayesian stochastic character mapping; transitions between biogeographic areas are indicated by changes in colour along the branches.

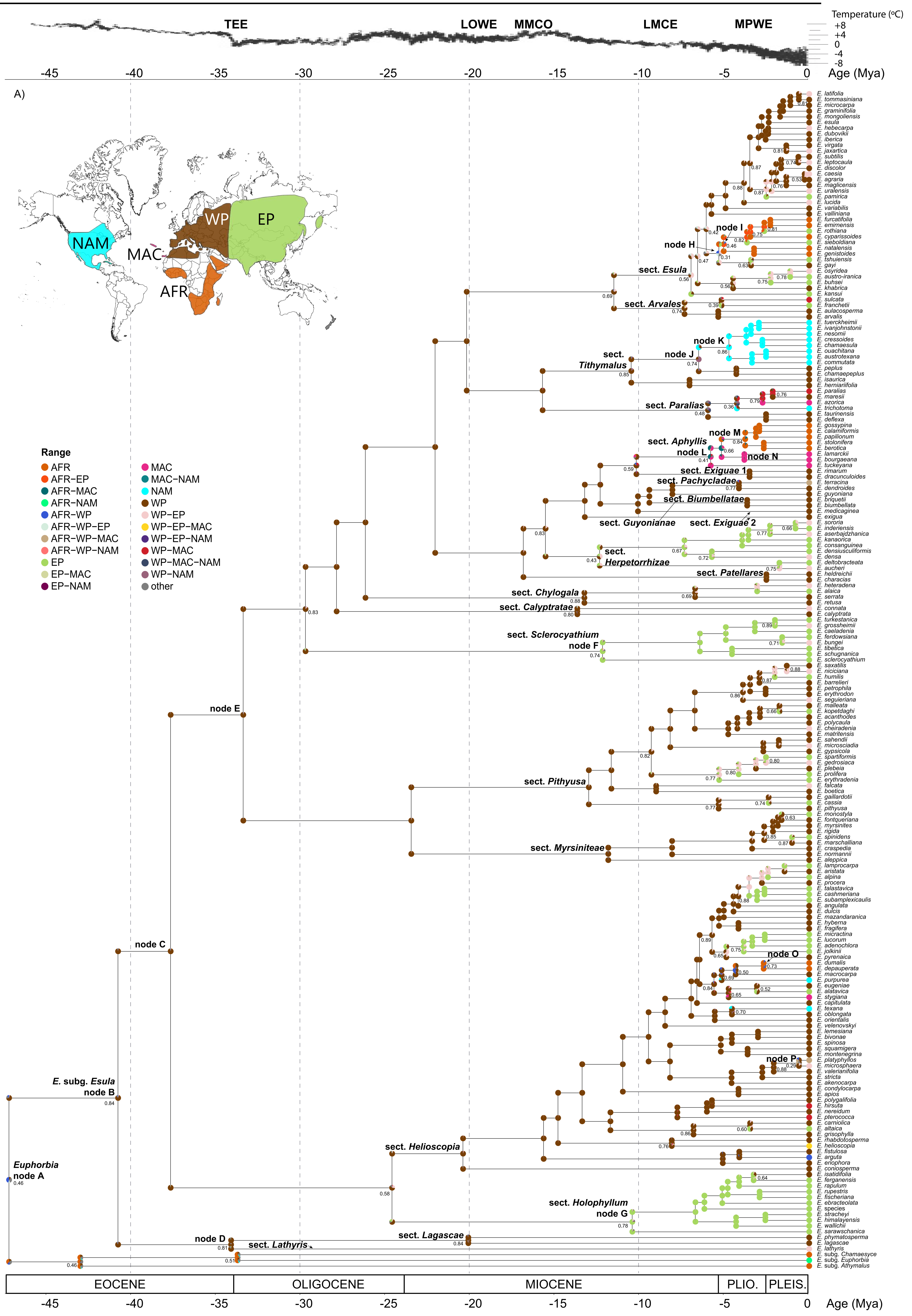

Temperature (ºC)
+8
+4
0
-4
-8
TEE
LOWE
MMCO
LMCE
MPWE
Age (Mya)
A)
NAM
MAC
WP
EP
AFR
Range
AFR
AFR−EP
AFR−MAC
AFR−NAM
AFR−WP
AFR−WP−EP
AFR−WP−MAC
AFR−WP−NAM
EP
EP−MAC
EP−NAM
MAC
MAC−NAM
NAM
WP
WP−EP
WP−EP−MAC
WP−EP−NAM
WP−MAC
WP−MAC−NAM
WP−NAM
other
sect. Esula
sect. Arvales
sect. Tithymalus
sect. Paralias
sect. Aphyllis
sect. Exiguae 1
sect. Pachycladae
sect. Biumbellatae
sect. Guyonianae
sect. Exiguae 2
sect. Herpetorrhizae
sect. Patellares
sect. Chylogala
sect. Calyptratae
sect. Sclerocyathium
sect. Pithyusa
sect. Myrsiniteae
sect. Helioscopia
sect. Holophyllum
sect. Lagascae
sect. Lathyris
node A
node B
node C
node D
node E
node F
node G
node H
node I
node J
node K
node L
node M
node N
node O
node P
Euphorbia
E. subg. Esula
EOCENE
OLIGOCENE
MIOCENE
PLIO.
PLEIS.

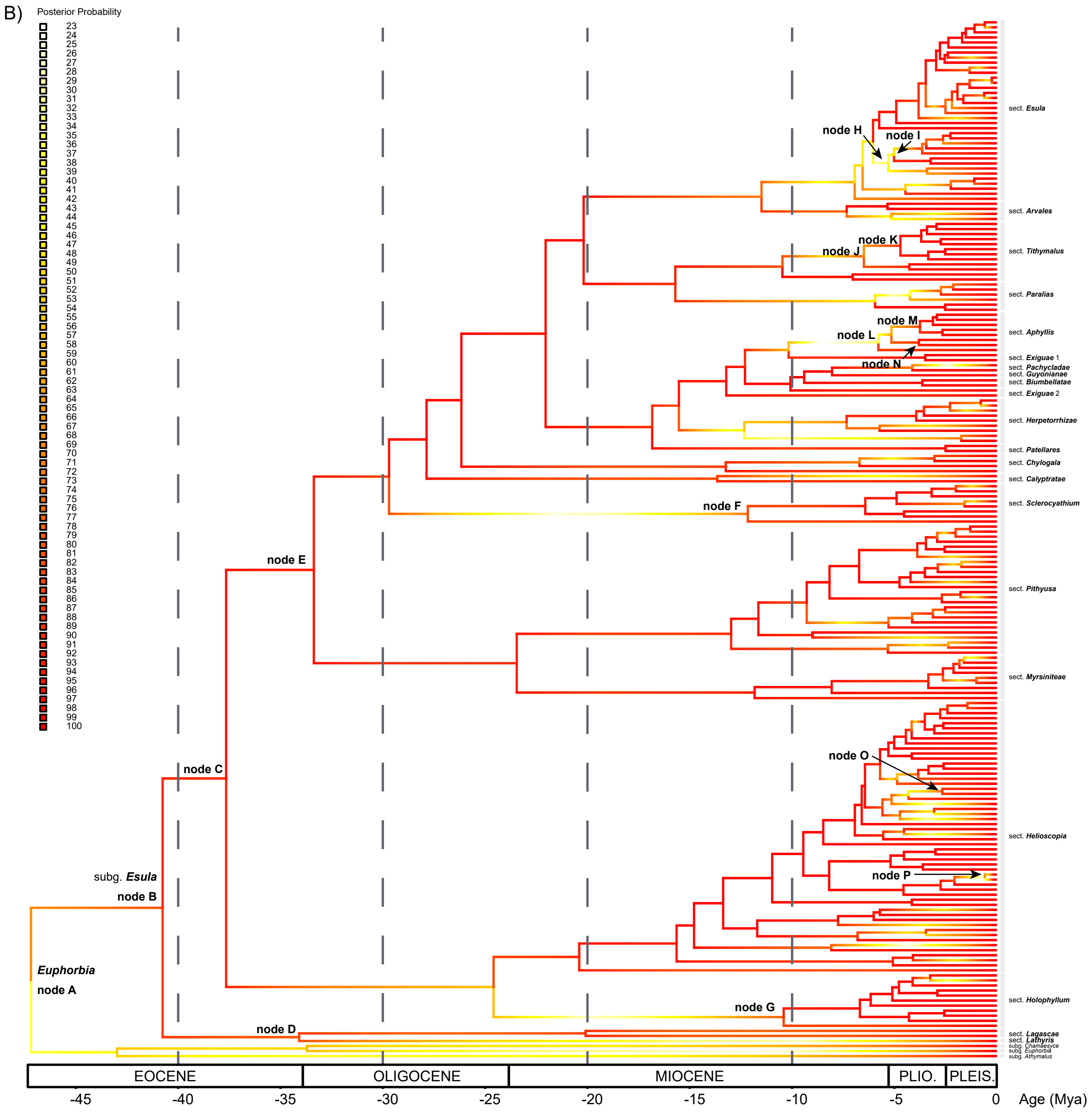
B)
Posterior Probability
sect. Esula
node H
node I
sect. Arvales
node K
node J
sect. Tithymalus
sect. Paralias
node M
node L
sect. Aphyllis
node N
sect. Exiguae 1
sect. Pachycladae
sect. Guyonianae
sect. Biumbellatae
sect. Exiguae 2
sect. Herpetorrhizae
sect. Patellares
sect. Chylogala
sect. Calyptratae
node F
sect. Sclerocyathium
node E
sect. Pithyusa
sect. Myrsiniteae
node C
node O
sect. Helioscopia
subg. Esula
node B
node P
Euphorbia
node A
node G
sect. Holophyllum
node D
sect. Lagascae
sect. Lathyris
subg. Chamaesyce
subg. Euphorbia
subg. Athymalus
EOCENE
OLIGOCENE
MIOCENE
PLIO.
PLEIS.
-45
-40
-35
-30
-25
-20
-15
-10
-5
0
Age (Mya)

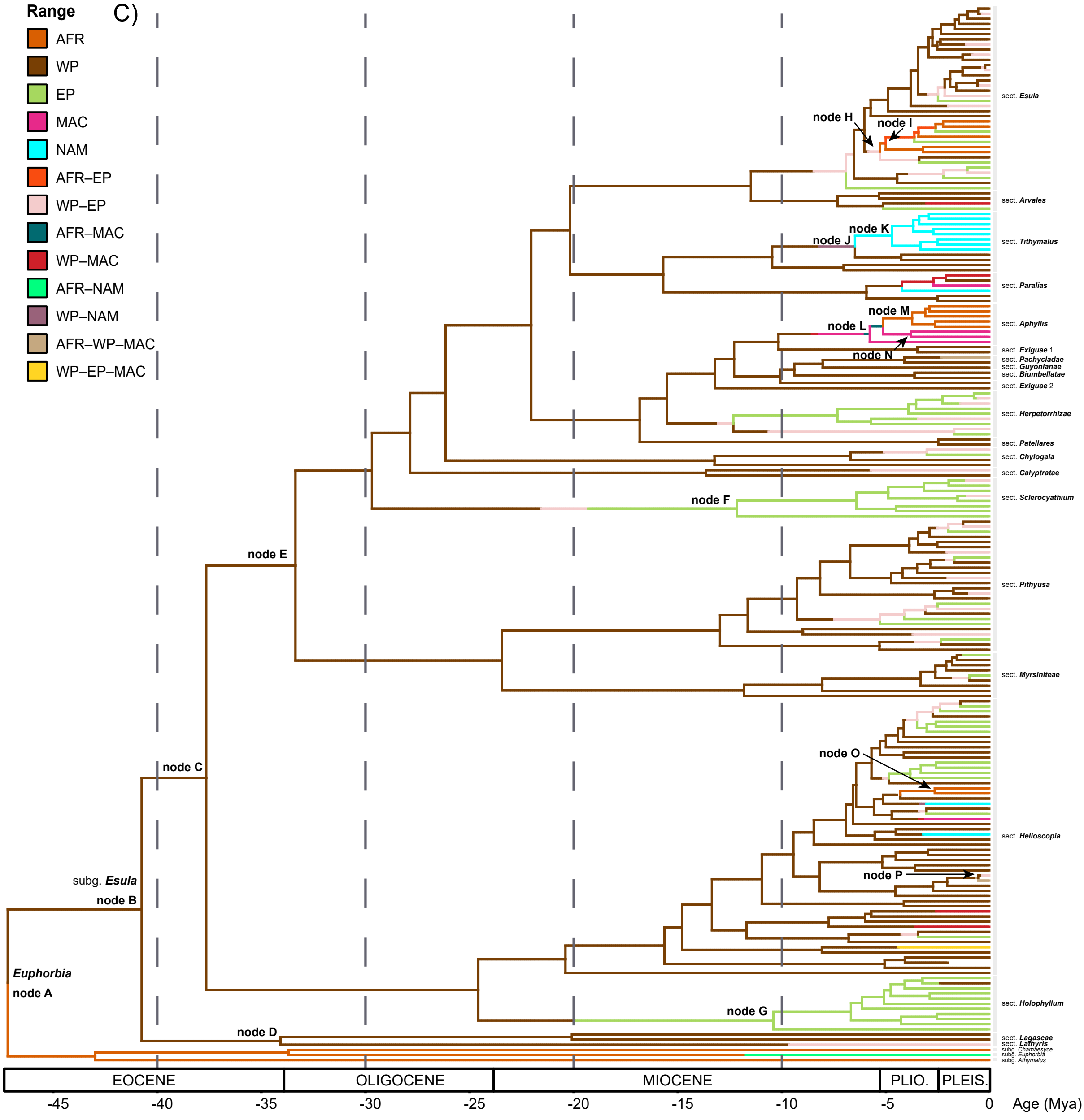

C)
Range
AFR
WP
EP
MAC
NAM
AFR–EP
WP–EP
AFR–MAC
WP–MAC
AFR–NAM
WP–NAM
AFR–WP–MAC
WP–EP–MAC
node A
node B
node C
node D
node E
node F
node G
node H
node I
node J
node K
node L
node M
node N
node O
node P
Euphorbia
subg. Esula
sect. Esula
sect. Arvales
sect. Tithymalus
sect. Paralias
sect. Aphyllis
sect. Exiguae 1
sect. Pachycladae
sect. Guyonianae
sect. Biumbellatae
sect. Exiguae 2
sect. Herpetorrhizae
sect. Patellares
sect. Chylogala
sect. Calyptratae
sect. Sclerocyathium
sect. Pithyusa
sect. Myrsiniteae
sect. Helioscopia
sect. Holophyllum
sect. Lagascae
sect. Lathyris
subg. Chamaesyce
subg. Euphorbia
subg. Athymalus
EOCENE
OLIGOCENE
MIOCENE
PLIO.
PLEIS.
-45
-40
-35
-30
-25
-20
-15
-10
-5
0
Age (Mya)
