## Supplementary Figure 5 for "The conquest and diversification of leafy spurges across the Holarctic and beyond: biogeography and evolution of life-history of *Euphorbia* subgenus *Esula*"

**Fig. S5**. Results under M1 biogeographic model of *Euphorbia* subg. *Esula* and closest allies. The phylogenetic tree is the maximum clade credibility (MCC) tree pruned from BEAST shown in Figs. 2G, 2H. As representatives of the other three *Euphorbia* subgenera, one taxon per subgenus representing its main distributional area. To reduce the size of the tree, those clades within *E.* subg. *Esula* with ages less than 2.5 Mya that had the same distribution were eliminated. **(A)** Maximum a posteriori (MAP) tree showing support values, in the form of marginal posterior probabilities, for the transition events between ancestral ranges reconstructed along branches in Fig. S5B using stochastic mapping; boxes on the left indicate marginal posterior probabilities for the inferences depicted as colours. **(B)** MAP reconstruction of the spatio-temporal evolution in *E.* subg. *Esula* simulated under Bayesian stochastic character mapping implemented in RevBayes; transitions between biogeographic areas are indicated by changes in colour along the branches.

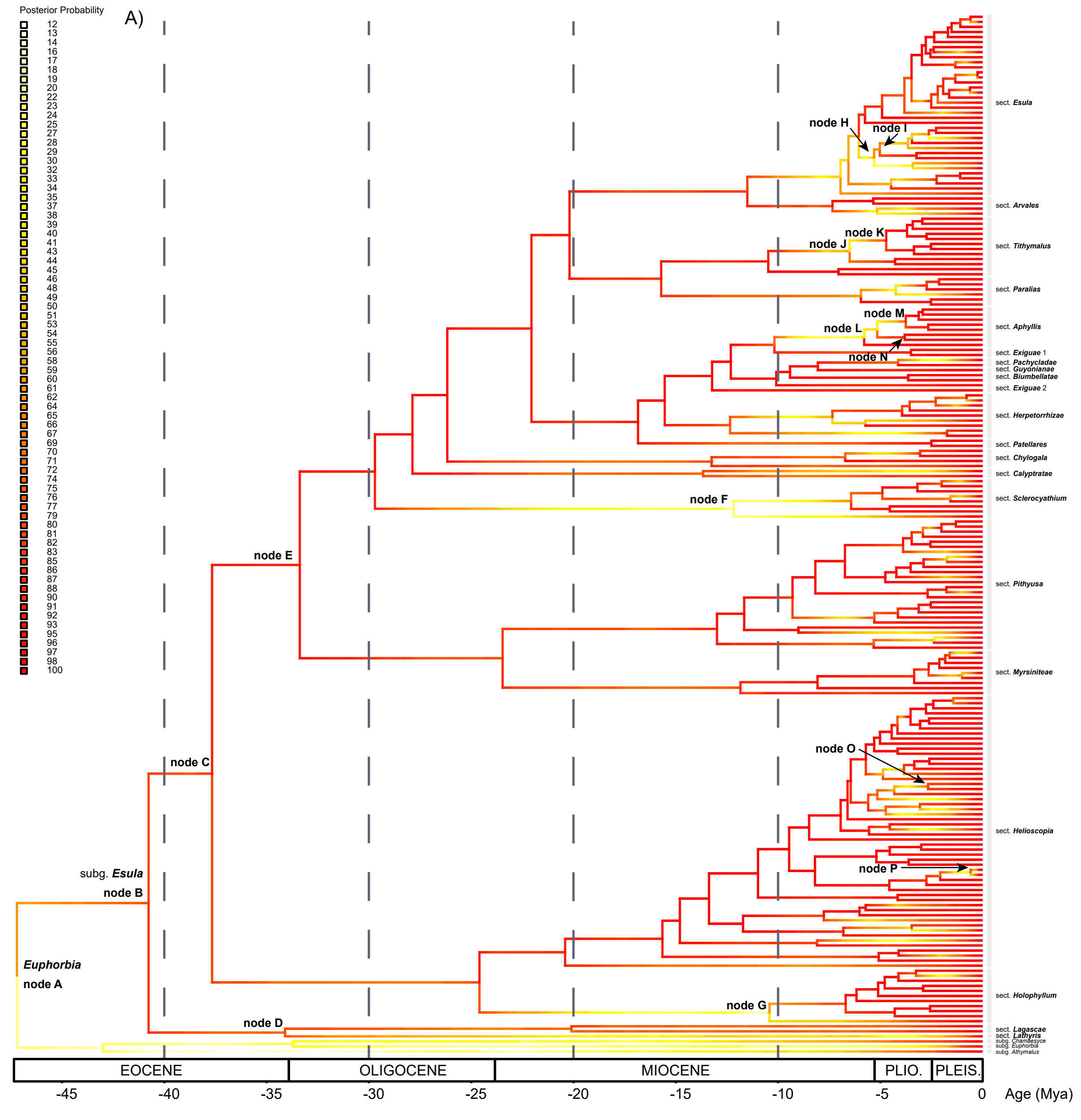

Posterior Probability
12
13
14
16
17
18
19
20
22
23
24
25
27
28
29
30
32
33
34
35
37
38
39
40
41
43
44
45
46
48
49
50
51
53
54
55
56
58
59
60
61
62
64
65
66
67
69
70
71
72
74
75
76
77
79
80
81
82
83
85
86
87
88
90
91
92
93
95
96
97
98
100
A)
node H
node I
node K
node J
node M
node L
node N
node F
node E
node C
node O
node P
subg. Esula
node B
Euphorbia
node A
node G
node D
sect. Esula
sect. Arvales
sect. Tithymalus
sect. Paralias
sect. Aphyllis
sect. Exiguae 1
sect. Pachycladae
sect. Guyonianae
sect. Biumbellatae
sect. Exiguae 2
sect. Herpetorrhizae
sect. Patellares
sect. Chylogala
sect. Calyptratae
sect. Sclerocyathium
sect. Pithyusa
sect. Myrsiniteae
sect. Helioscopia
sect. Holophyllum
sect. Lagascae
sect. Lathyris
subg. Chamaesyce
subg. Euphorbia
subg. Athymalus
EOCENE
OLIGOCENE
MIOCENE
PLIO.
PLEIS.
-45
-40
-35
-30
-25
-20
-15
-10
-5
0
Age (Mya)

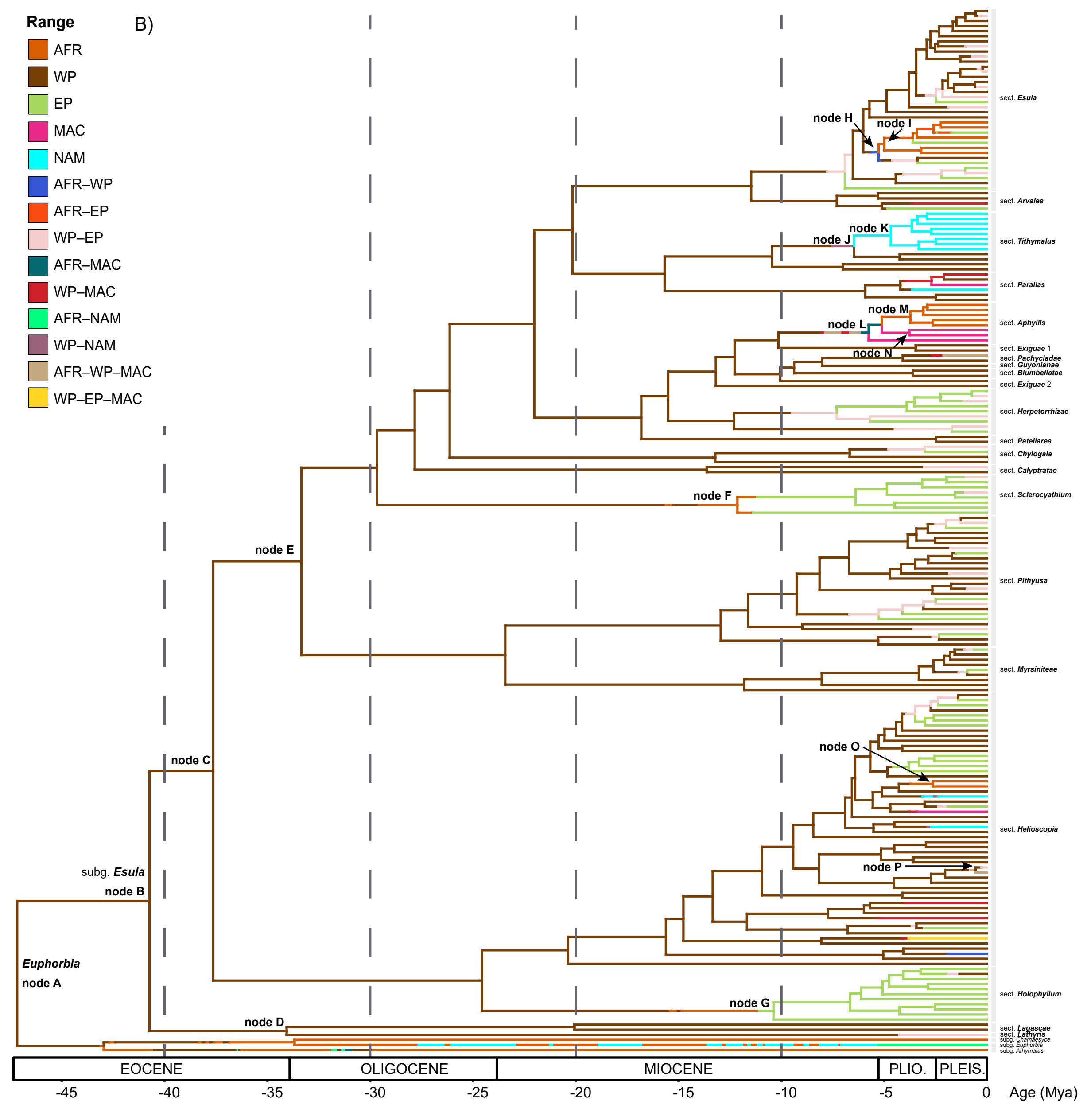

Range
B)
AFR
WP
EP
MAC
NAM
AFR–WP
AFR–EP
WP–EP
AFR–MAC
WP–MAC
AFR–NAM
WP–NAM
AFR–WP–MAC
WP–EP–MAC
node H
node I
node K
node J
node M
node L
node N
node F
node E
node C
node O
node P
subg. Esula
node B
Euphorbia
node A
node G
node D
sect. Esula
sect. Arvales
sect. Tithymalus
sect. Paralias
sect. Aphyllis
sect. Exiguae 1
sect. Pachycladae
sect. Guyonianae
sect. Biumbellatae
sect. Exiguae 2
sect. Herpetorrhizae
sect. Patellares
sect. Chylogala
sect. Calyptratae
sect. Sclerocyathium
sect. Pithyusa
sect. Myrsiniteae
sect. Helioscopia
sect. Holophyllum
sect. Lagascae
sect. Lathyris
subg. Chamaesyce
subg. Euphorbia
subg. Athymalus
EOCENE
OLIGOCENE
MIOCENE
PLIO.
PLEIS.
-45
-40
-35
-30
-25
-20
-15
-10
-5
0
Age (Mya)
