## Supplementary Figure 6 for "The conquest and diversification of leafy spurges across the Holarctic and beyond: biogeography and evolution of life-history of *Euphorbia* subgenus *Esula*"

**Fig. S6**. Estimates of speciation and extinction rates of *Euphorbia* subg. *Esula* using the Horseshoe Markov random field episodic birth-death model implemented in RevBayes.

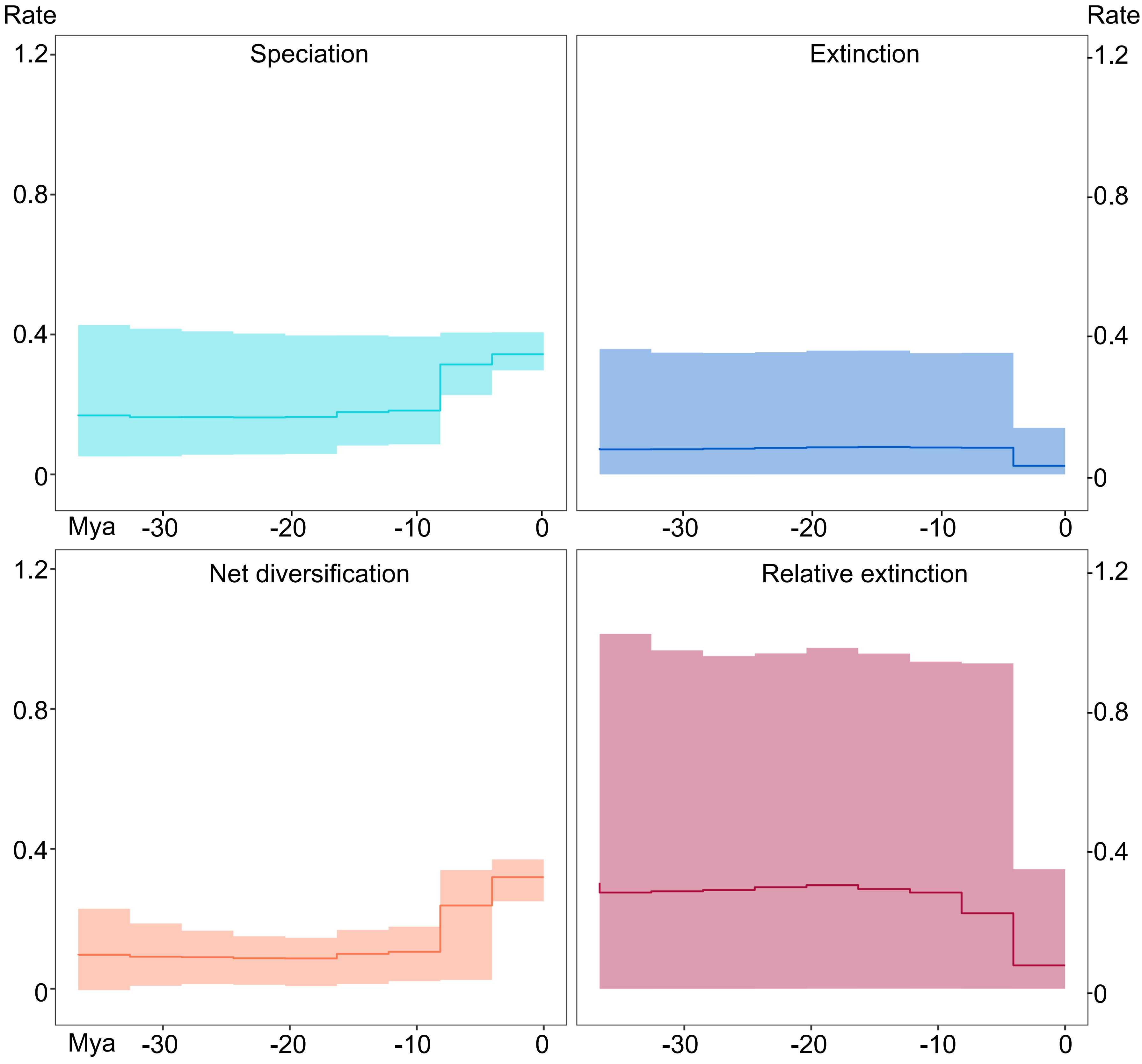
Rate
Rate
Speciation
Extinction
Net diversification
Relative extinction
1.2
0.8
0.4
0
Mya
-30
-20
-10
0
