## Supplementary Figure 7 for "The conquest and diversification of leafy spurges across the Holarctic and beyond: biogeography and evolution of life-history of *Euphorbia* subgenus *Esula*"

**Fig. S7**. Changes in diversification rates over the evolutionary history of *Euphorbia* subg. *Esula* allowing mass extinction events (MEEs) under a uniform sampling strategy, inferred with CoMET. The first two plots show detection of the time of MEEs using Bayes factors (BF) comparisons (BF < 2, negative support; Kass and Raftery, 1995). Results allowing changes at discrete times of diversification, speciation, extinction and relative-extinction rates.

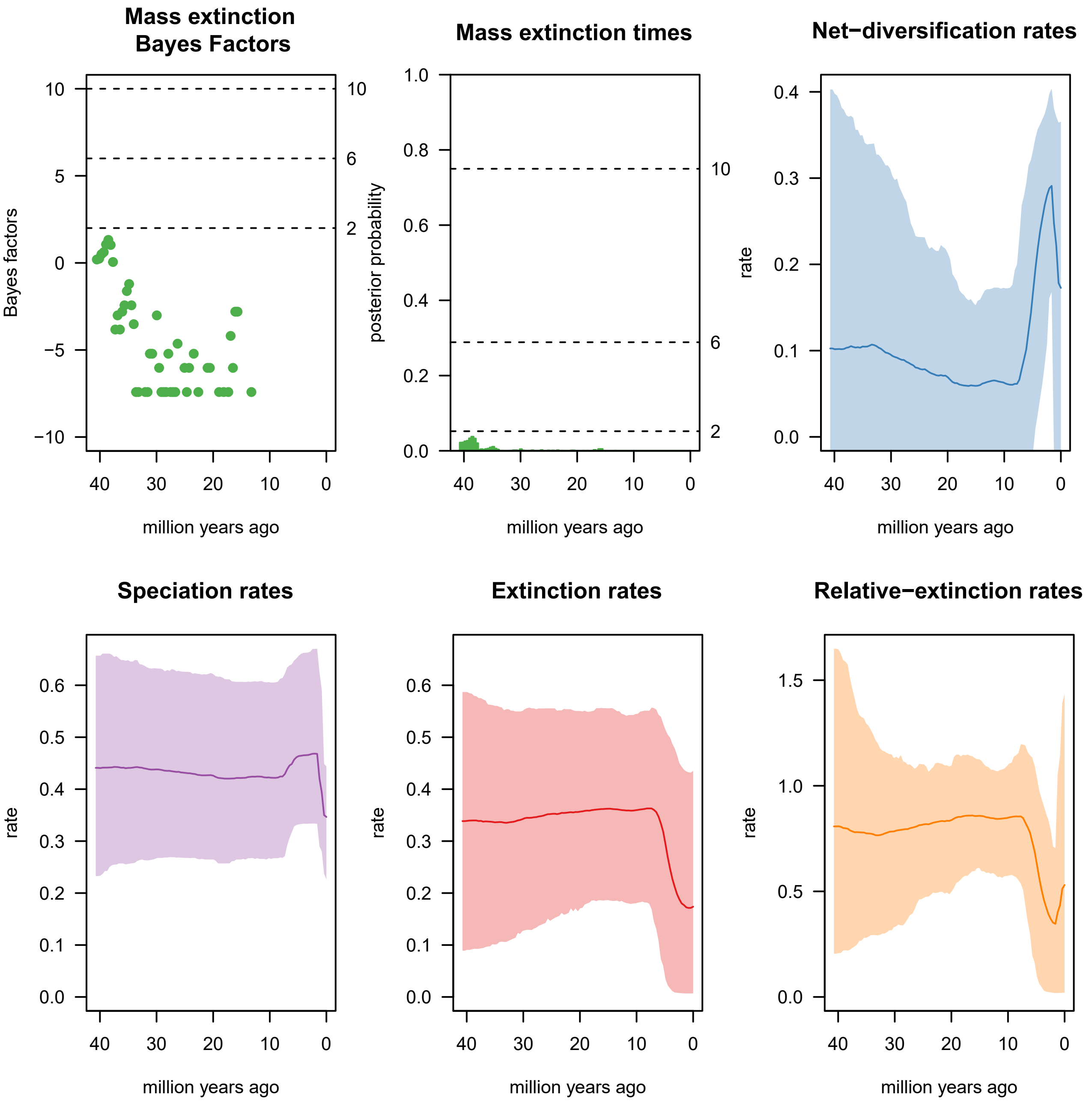

Mass extinction Bayes Factors
Bayes factors
10
5
0
−5
−10
10
6
2
40
30
20
10
0
million years ago
Mass extinction times
posterior probability
1.0
0.8
0.6
0.4
0.2
0.0
10
6
2
40
30
20
10
0
million years ago
Net−diversification rates
rate
0.4
0.3
0.2
0.1
0.0
40
30
20
10
0
million years ago
Speciation rates
rate
0.6
0.5
0.4
0.3
0.2
0.1
0.0
40
30
20
10
0
million years ago
Extinction rates
rate
0.6
0.5
0.4
0.3
0.2
0.1
0.0
40
30
20
10
0
million years ago
Relative−extinction rates
rate
1.5
1.0
0.5
0.0
40
30
20
10
0
million years ago
