## Supplementary Figure 8 for "The conquest and diversification of leafy spurges across the Holarctic and beyond: biogeography and evolution of life-history of *Euphorbia* subgenus *Esula*"

**Fig. S8**. Changes in diversification rates over the evolutionary history of *Euphorbia* subg. *Esula* allowing mass extinction events under an incomplete taxon sampling strategy, inferred with CoMET.

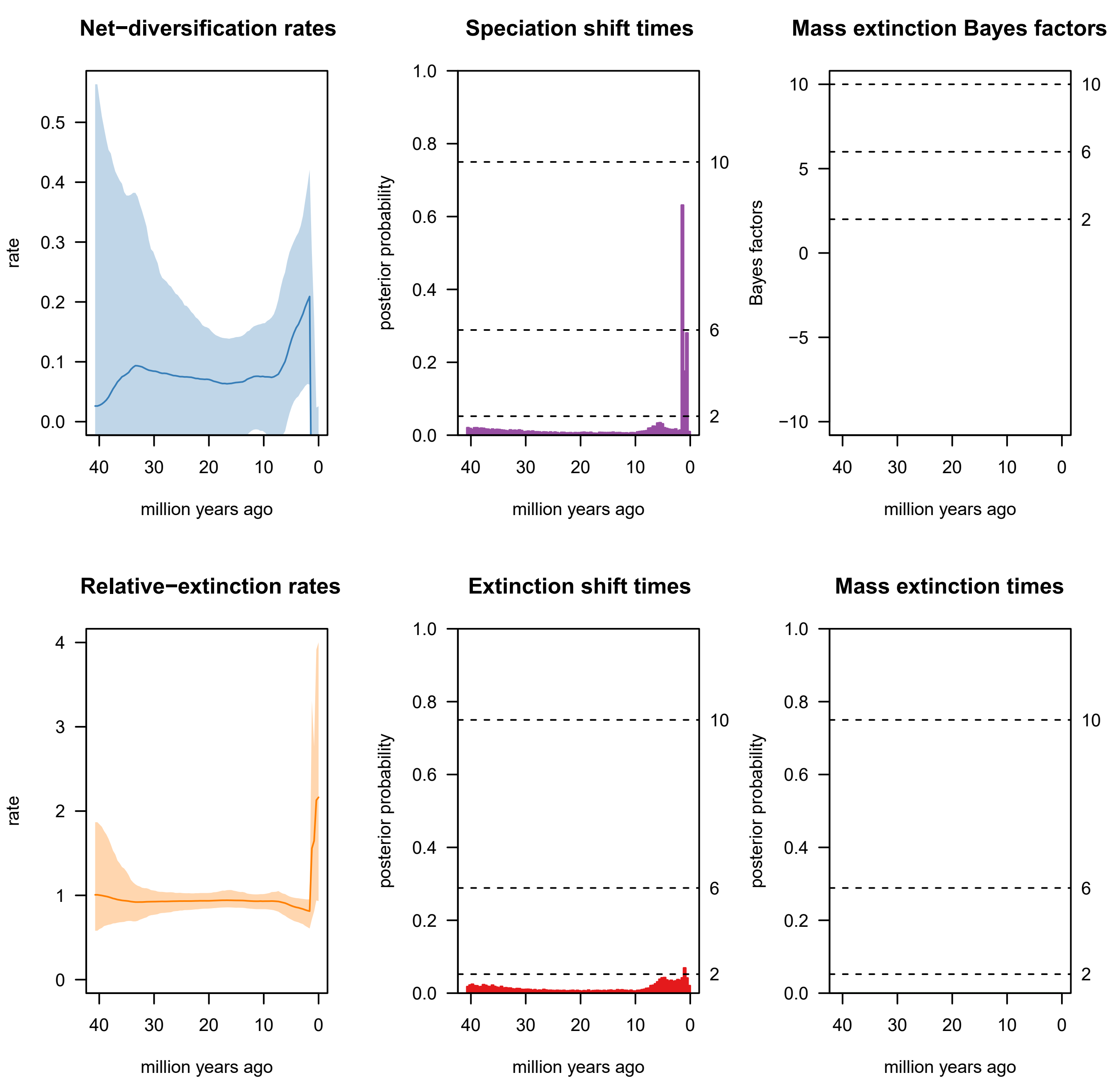
Net−diversification rates
rate
0.0
0.1
0.2
0.3
0.4
0.5
40
30
20
10
0
million years ago
Speciation shift times
posterior probability
0.0
0.2
0.4
0.6
0.8
1.0
2
6
10
million years ago
Mass extinction Bayes factors
Bayes factors
−10
−5
0
5
10
2
6
10
million years ago
Relative−extinction rates
rate
0
1
2
3
4
million years ago
Extinction shift times
posterior probability
million years ago
Mass extinction times
posterior probability
million years ago
