## Supplementary Figure 9 for "The conquest and diversification of leafy spurges across the Holarctic and beyond: biogeography and evolution of life-history of *Euphorbia* subgenus *Esula*"

**Fig. S9**. Effective Sample Size (ESS) of *Euphorbia* subg. *Esula* for the different parameters estimated using the R package *TESS*. ESS > 500 suggests convergence of the chain.

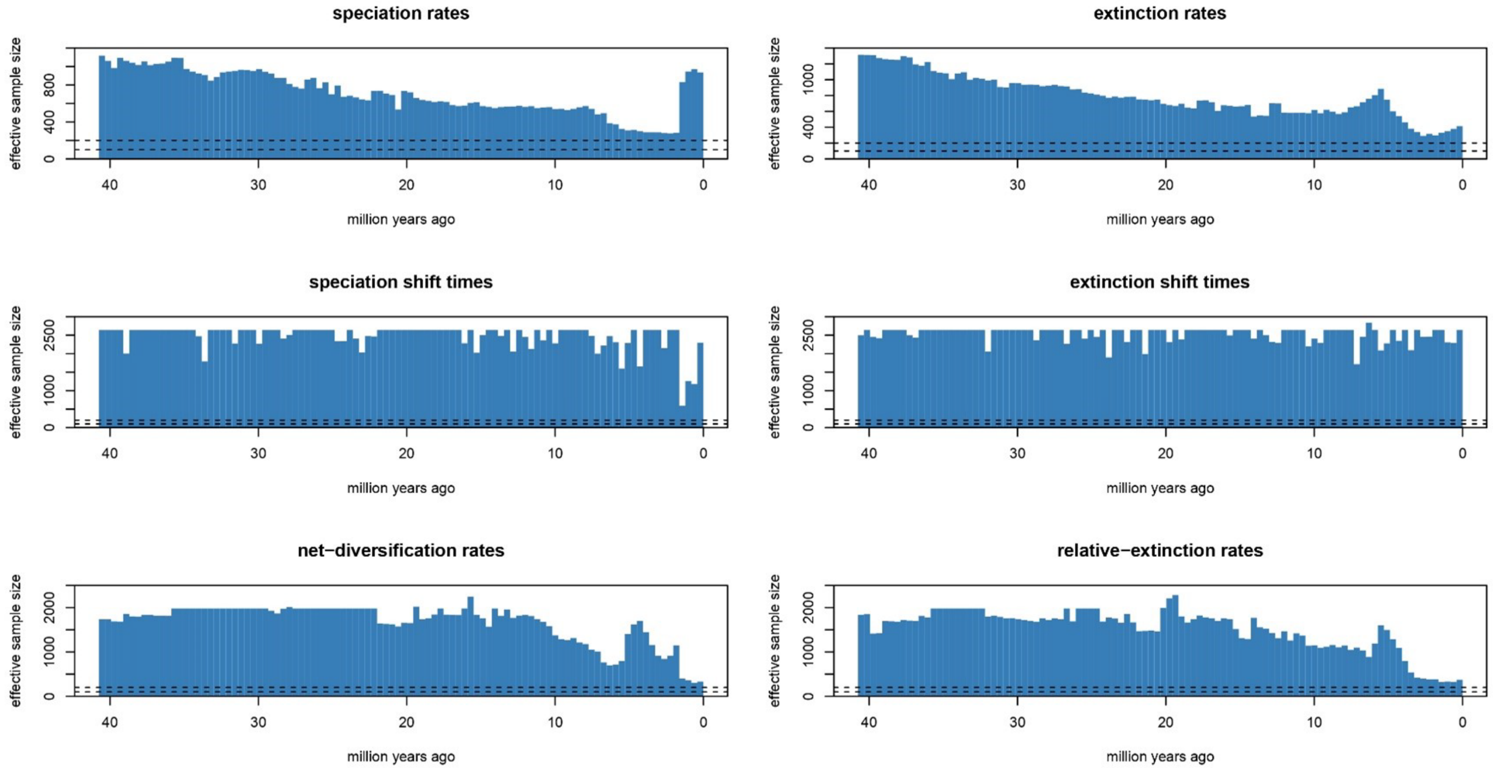

speciation rates
extinction rates
speciation shift times
extinction shift times
net−diversification rates
relative−extinction rates
effective sample size
million years ago
40
30
20
10
0
400
800
1000
2500
2000
