## Supplementary Figure 10 for "The conquest and diversification of leafy spurges across the Holarctic and beyond: biogeography and evolution of life-history of *Euphorbia* subgenus *Esula*"

**Fig. S10**. Lineage-specific birth-death (LSBDS) model representing the extinction rate across the evolution of *Euphorbia* subg. *Esula*. Extinction rate ($\mu$) is high and constant over time: 1 extinction event every 10,000 years.

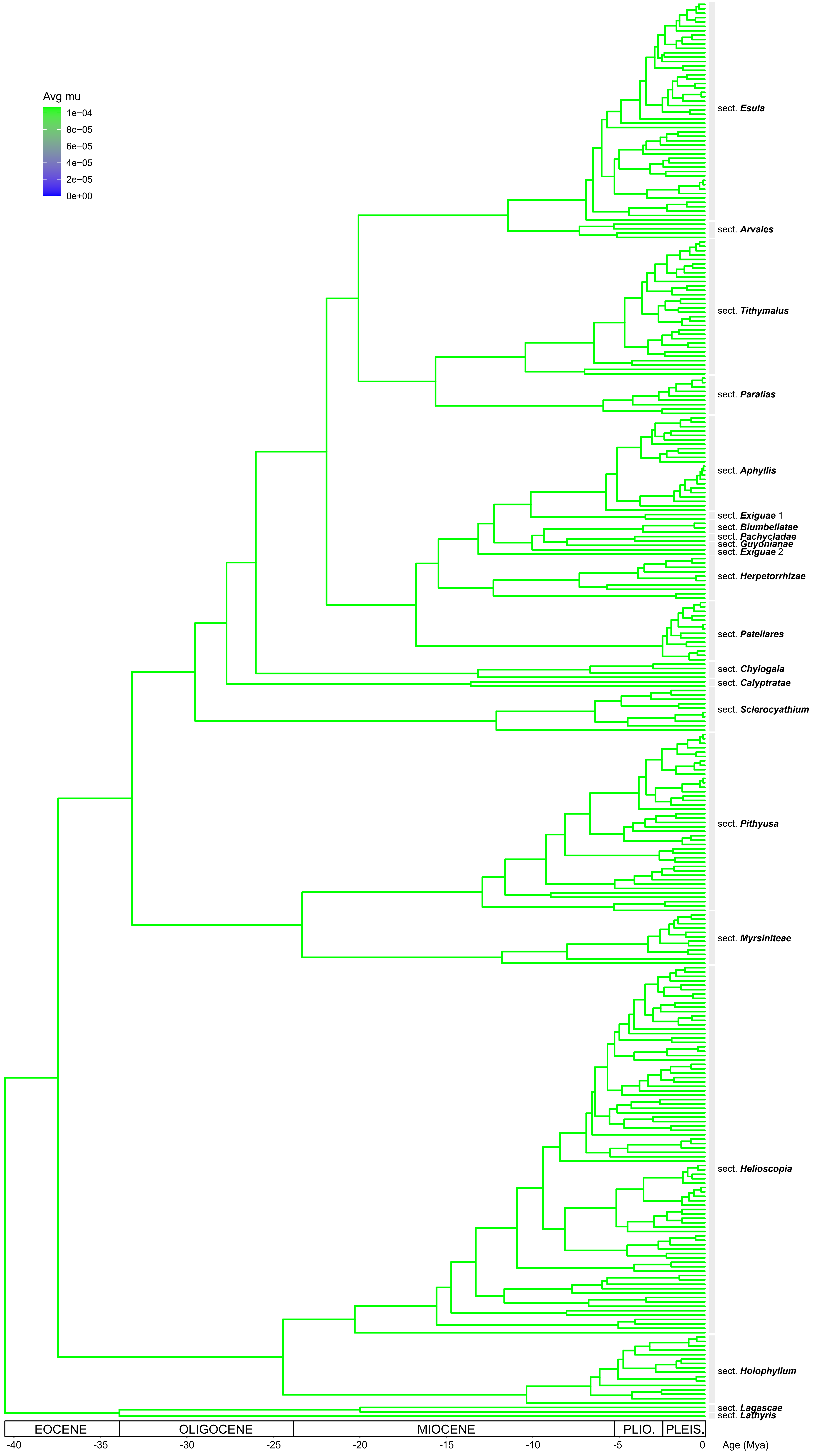
Avg mu
1e−04
8e−05
6e−05
4e−05
2e−05
0e+00
sect. Esula
sect. Arvales
sect. Tithymalus
sect. Paralias
sect. Aphyllis
sect. Exiguae 1
sect. Biumbellatae
sect. Pachycladae
sect. Guyonianae
sect. Exiguae 2
sect. Herpetorrhizae
sect. Patellares
sect. Chylogala
sect. Calyptratae
sect. Sclerocyathium
sect. Pithyusa
sect. Myrsiniteae
sect. Helioscopia
sect. Holophyllum
sect. Lagascae
sect. Lathyris
EOCENE
OLIGOCENE
MIOCENE
PLIO.
PLEIS.
-40
-35
-30
-25
-20
-15
-10
-5
0
Age (Mya)
