## Supplementary Figure 11 for "The conquest and diversification of leafy spurges across the Holarctic and beyond: biogeography and evolution of life-history of *Euphorbia* subgenus *Esula*"

**Fig. S11**. BiSSE diversification parameters of *Euphorbia* subg. *Esula* implemented in RevBayes. Posterior densities of speciation ($\lambda$), extinction ($\mu$), relative extinction ($\mu/\lambda$), and net diversification ($\lambda$–$\mu$) rates. Changes in life-history from state 0 (annual, blue) to 1 (perennial, orange) are associated with diversification rate heterogeneity.

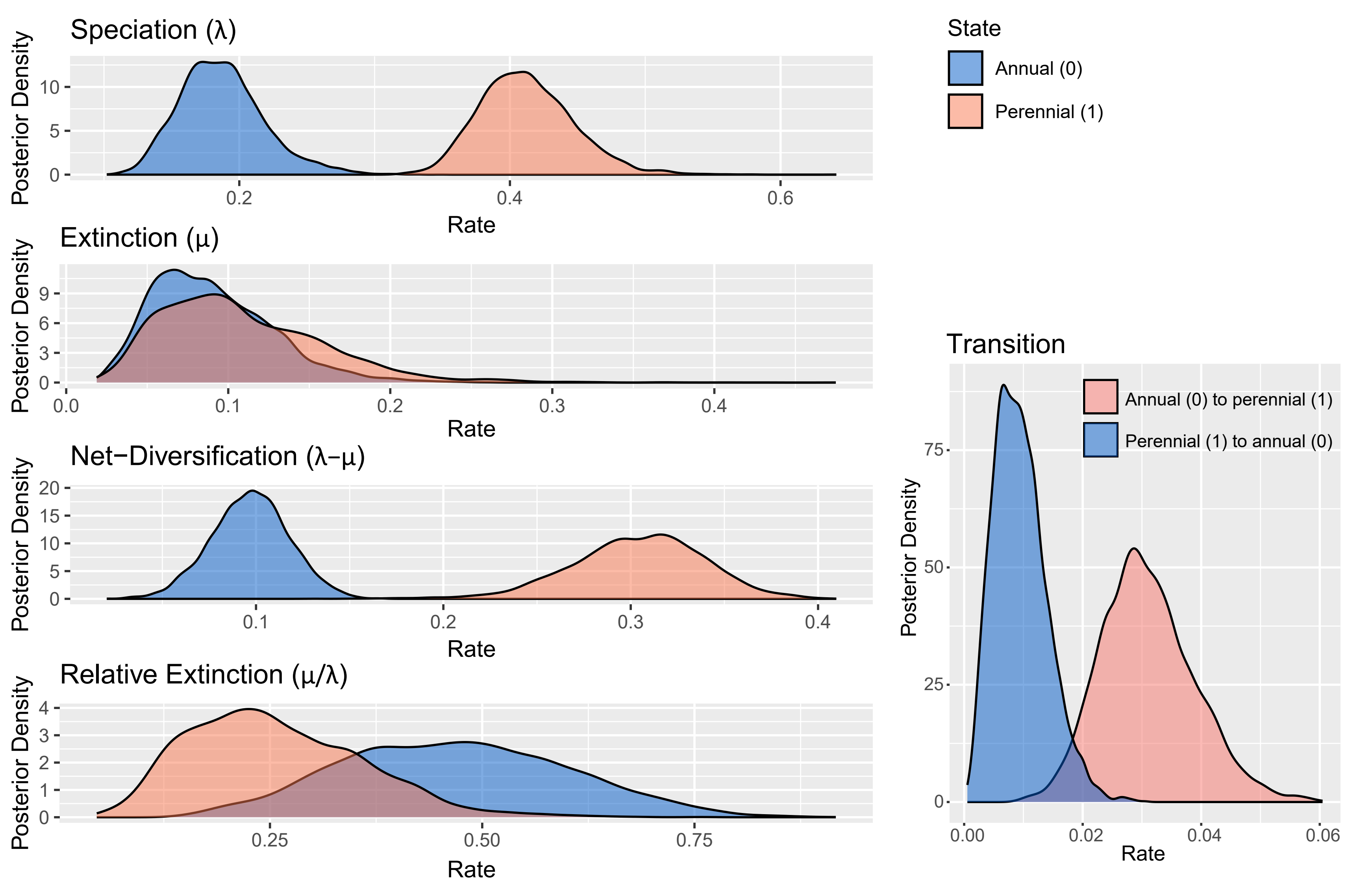
Speciation (λ)
Extinction (μ)
Net−Diversification (λ−μ)
Relative Extinction (μ/λ)
Transition
State
Annual (0)
Perennial (1)
Annual (0) to perennial (1)
Perennial (1) to annual (0)
Posterior Density
Rate
