## Supplementary Figure 12 for "The conquest and diversification of leafy spurges across the Holarctic and beyond: biogeography and evolution of life-history of *Euphorbia* subgenus *Esula*"

**Fig. S12**. Results from the BiSSE model of *Euphorbia* subg. *Esula* implemented in RevBayes. **(A)** Maximum a posteriori (MAP) reconstruction of life-history evolution simulated under Bayesian stochastic character mapping; divergence times in millions of years ago (Mya) are indicated by the axis at the bottom of the tree; transitions between states 0 (annual, blue) and 1 (perennial, orange) are indicated by changes in colour along the branches. **(B)** MAP tree showing support values, in the form of marginal posterior probabilities, for the transition events between states reconstructed along branches in Fig. S12A using stochastic mapping; inset to the left indicates posterior probability for the inferences depicted as colours.

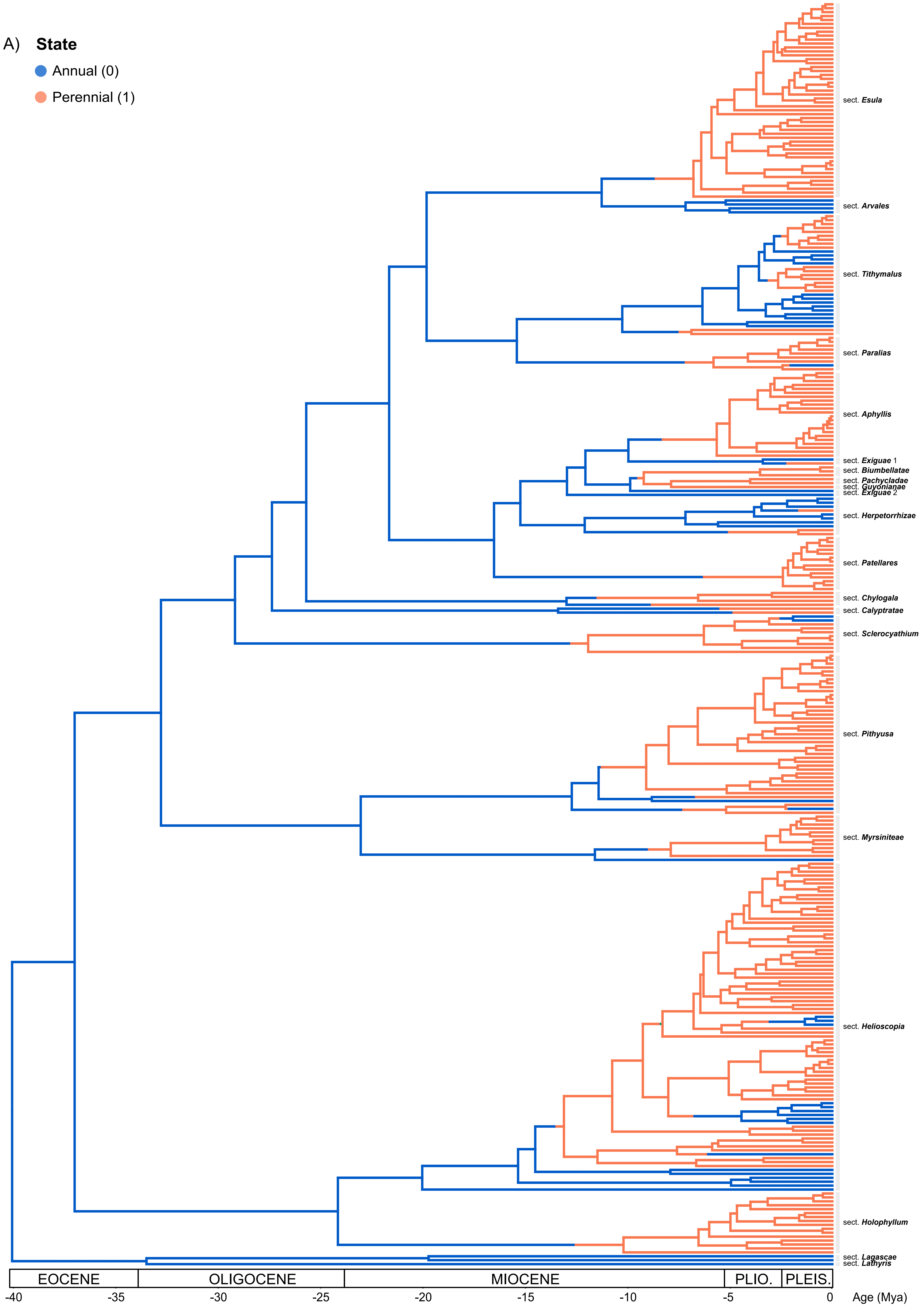
A)
State
Annual (0)
Perennial (1)
sect. Esula
sect. Arvales
sect. Tithymalus
sect. Paralias
sect. Aphyllis
sect. Exiguae 1
sect. Biumbellatae
sect. Pachycladae
sect. Guyonianae
sect. Exiguae 2
sect. Herpetorrhizae
sect. Patellares
sect. Chylogala
sect. Calyptratae
sect. Sclerocyathium
sect. Pithyusa
sect. Myrsiniteae
sect. Helioscopia
sect. Holophyllum
sect. Lagascae
sect. Lathyris
EOCENE
OLIGOCENE
MIOCENE
PLIO.
PLEIS.
-40
-35
-30
-25
-20
-15
-10
-5
0
Age (Mya)

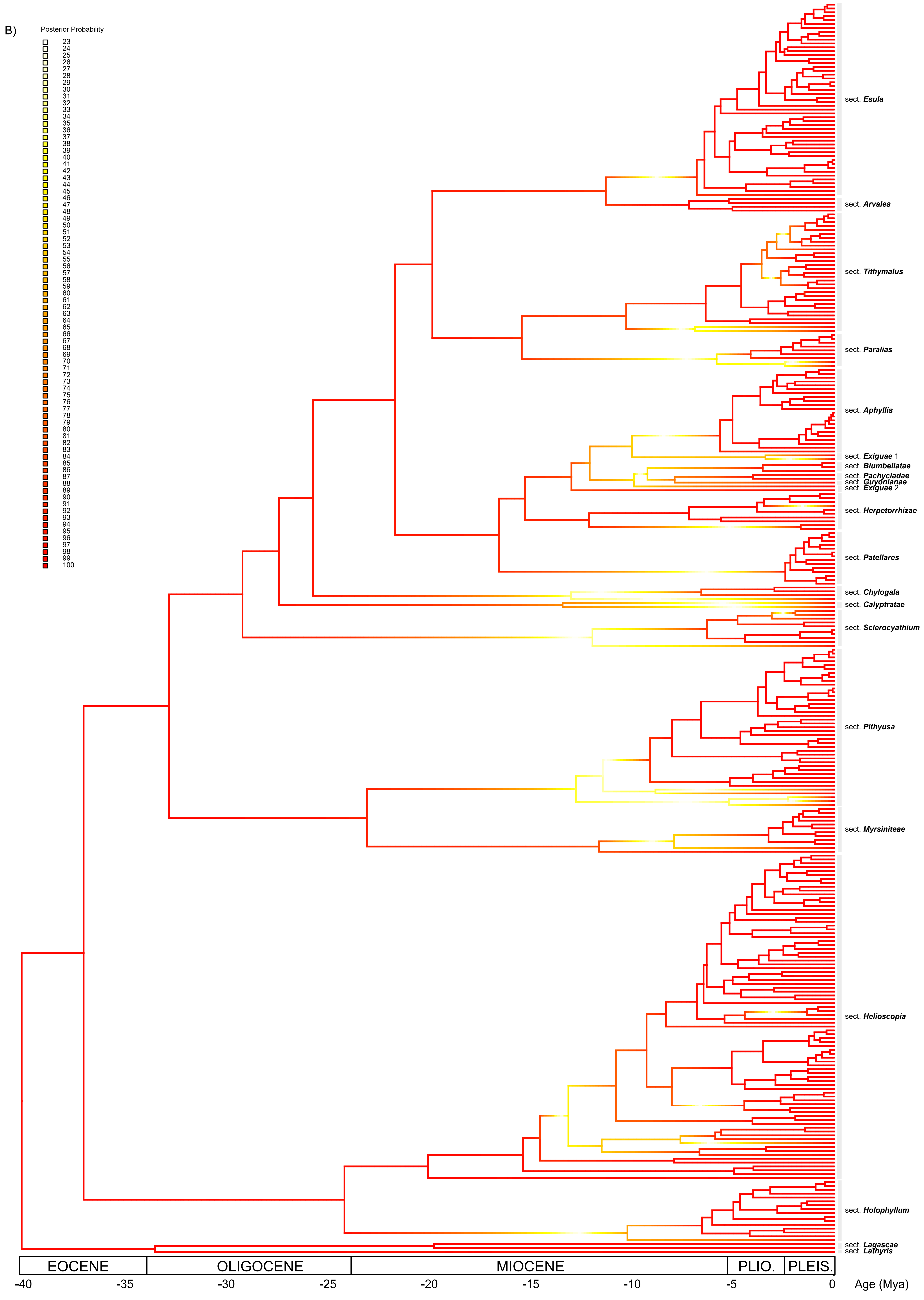
B)
Posterior Probability
sect. Esula
sect. Arvales
sect. Tithymalus
sect. Paralias
sect. Aphyllis
sect. Exiguae 1
sect. Biumbellatae
sect. Pachycladae
sect. Guyonianae
sect. Exiguae 2
sect. Herpetorrhizae
sect. Patellares
sect. Chylogala
sect. Calyptratae
sect. Sclerocyathium
sect. Pithyusa
sect. Myrsiniteae
sect. Helioscopia
sect. Holophyllum
sect. Lagascae
sect. Lathyris
EOCENE
OLIGOCENE
MIOCENE
PLIO.
PLEIS.
-40
-35
-30
-25
-20
-15
-10
-5
0
Age (Mya)
