## Supplementary Figure 13 for "The conquest and diversification of leafy spurges across the Holarctic and beyond: biogeography and evolution of life-history of *Euphorbia* subgenus *Esula*"

**Fig. S13**. HiSSE diversification parameters of *Euphorbia* subg. *Esula* implemented in RevBayes. Posterior densities of speciation (λ), extinction (μ), relative extinction (μ/λ) and net diversification (λ–μ) rates. Colours correspond to the posterior probabilities for annual (0, blue) and perennial (1, green) focal states associated with hidden states (A, B).

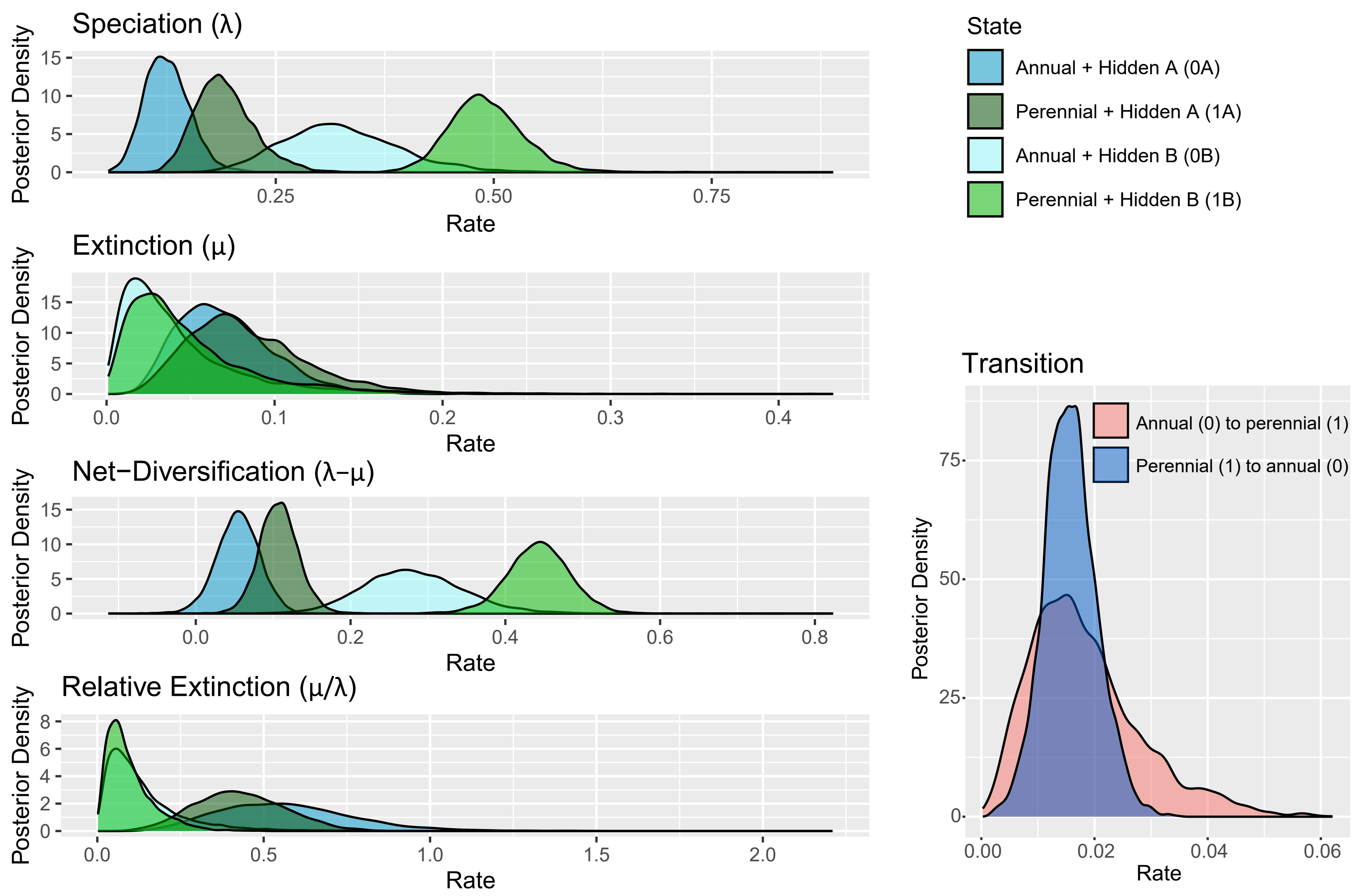

Speciation (λ)
Extinction (μ)
Net−Diversification (λ−μ)
Relative Extinction (μ/λ)
Posterior Density
Rate
State
Annual + Hidden A (0A)
Perennial + Hidden A (1A)
Annual + Hidden B (0B)
Perennial + Hidden B (1B)
Transition
Annual (0) to perennial (1)
Perennial (1) to annual (0)
