## Supplementary Figure 14 for "The conquest and diversification of leafy spurges across the Holarctic and beyond: biogeography and evolution of life-history of *Euphorbia* subgenus *Esula*"

**Fig. S14**. Differences in the HiSSE model between speciation and extinction rates of annual (0) and perennial (1) species of *Euphorbia* subg. *Esula* associated with hidden states (A, B). Differences in the transition rate from annual to perennial (01) and from perennial to annual (10) are also shown.

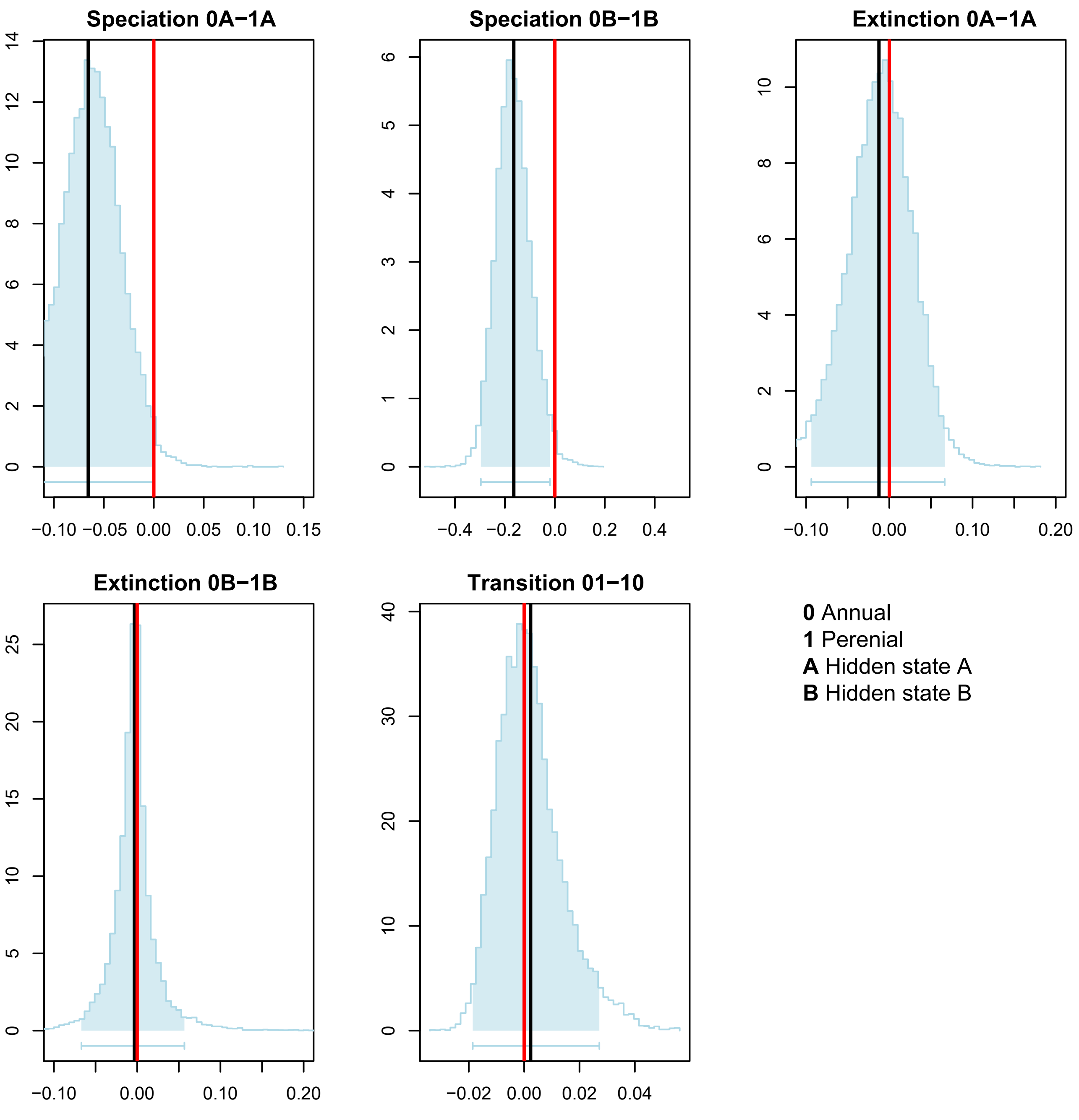
Speciation 0A−1A
Speciation 0B−1B
Extinction 0A−1A
Extinction 0B−1B
Transition 01−10
0 Annual
1 Perenial
A Hidden state A
B Hidden state B
