## Supplementary Figure 15 for "The conquest and diversification of leafy spurges across the Holarctic and beyond: biogeography and evolution of life-history of *Euphorbia* subgenus *Esula*"

**Fig. S15**. Results from the HiSSE model of *Euphorbia* subg. *Esula* implemented in RevBayes. **(A)** Maximum a Posteriori (MAP) tree showing reconstructed ancestral focal states (0, annual, blue; 1, perennial, green) and hidden states (A, B); size of circles represents marginal posterior probabilities. **(B)** Life-history evolution simulated under Bayesian stochastic character mapping; divergence times in millions of years ago (Mya) are shown by the axis at the bottom of the tree; branch colours denote the annual (0, blue) and perennial (1, green) observed states, associated with the hidden states (A or B); transitions between character states are indicated by changes in colour along the branches. **(C)** MAP tree showing support values, in the form of marginal posterior probabilities, for the transition events between states reconstructed along branches in (B) using stochastic mapping; inset to the left indicates posterior probability for the inferences depicted as colours.

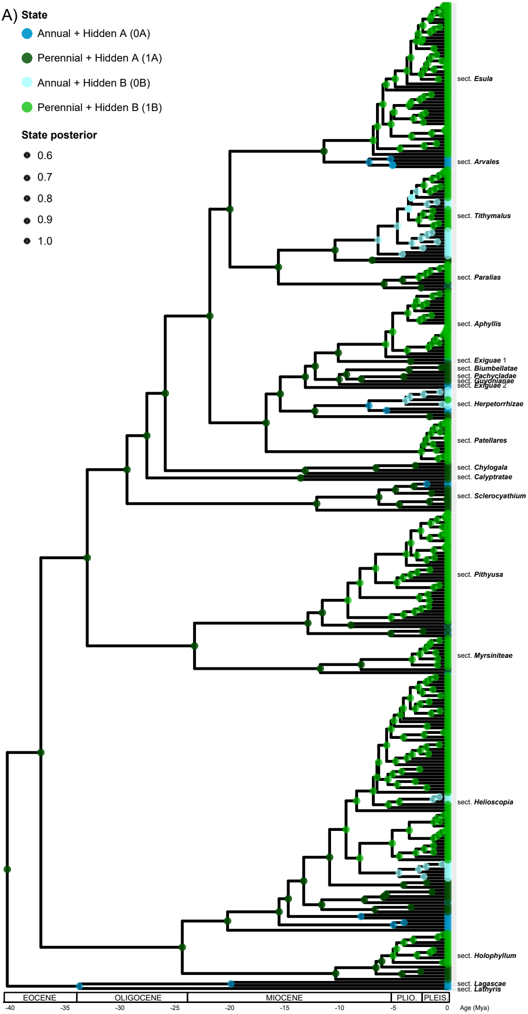

A)
State
Annual + Hidden A (0A)
Perennial + Hidden A (1A)
Annual + Hidden B (0B)
Perennial + Hidden B (1B)
State posterior
0.6
0.7
0.8
0.9
1.0
sect. Esula
sect. Arvales
sect. Tithymalus
sect. Paralias
sect. Aphyllis
sect. Exiguae 1
sect. Biumbellatae
sect. Pachycladae
sect. Guyonianae
sect. Exiguae 2
sect. Herpetorrhizae
sect. Patellares
sect. Chylogala
sect. Calyptratae
sect. Sclerocyathium
sect. Pithyusa
sect. Myrsiniteae
sect. Helioscopia
sect. Holophyllum
sect. Lagascae
sect. Lathyris
EOCENE
OLIGOCENE
MIOCENE
PLIO.
PLEIS.
-40
-35
-30
-25
-20
-15
-10
-5
0
Age (Mya)

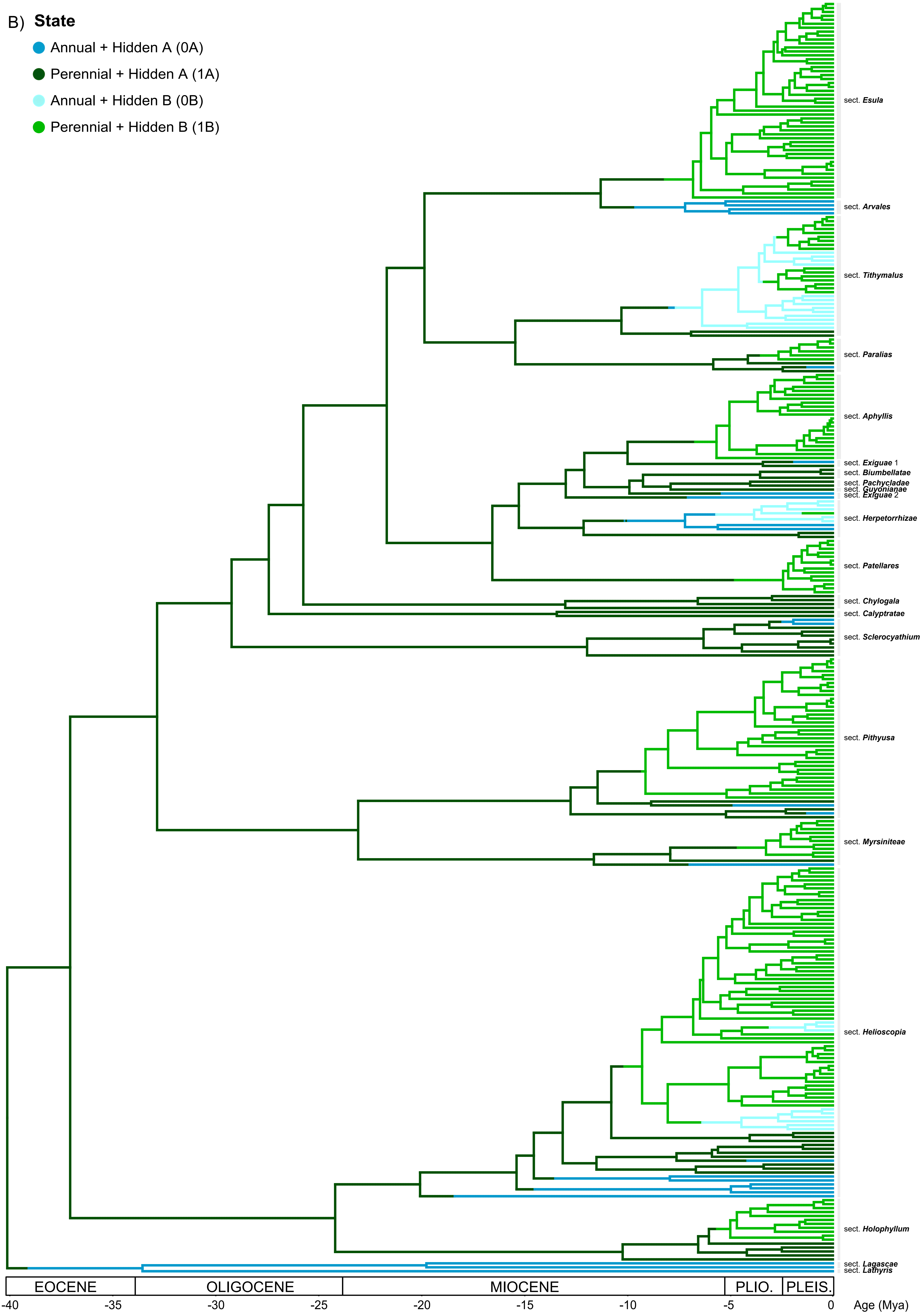

B)
State
Annual + Hidden A (0A)
Perennial + Hidden A (1A)
Annual + Hidden B (0B)
Perennial + Hidden B (1B)
sect. Esula
sect. Arvales
sect. Tithymalus
sect. Paralias
sect. Aphyllis
sect. Exiguae 1
sect. Biumbellatae
sect. Pachycladae
sect. Guyonianae
sect. Exiguae 2
sect. Herpetorrhizae
sect. Patellares
sect. Chylogala
sect. Calyptratae
sect. Sclerocyathium
sect. Pithyusa
sect. Myrsiniteae
sect. Helioscopia
sect. Holophyllum
sect. Lagascae
sect. Lathyris
EOCENE
OLIGOCENE
MIOCENE
PLIO.
PLEIS.
-40
-35
-30
-25
-20
-15
-10
-5
0
Age (Mya)

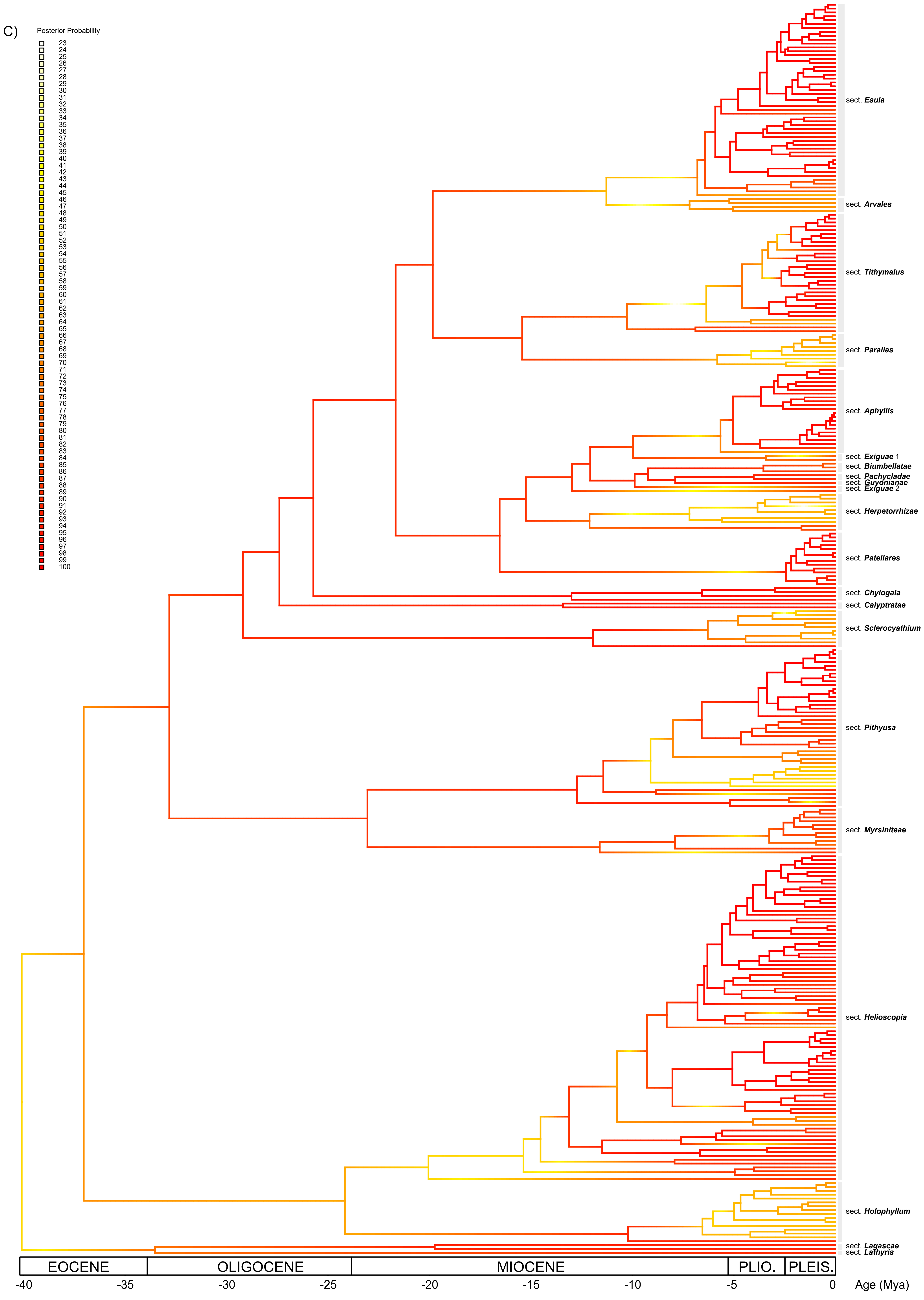

C)
Posterior Probability
sect. Esula
sect. Arvales
sect. Tithymalus
sect. Paralias
sect. Aphyllis
sect. Exiguae 1
sect. Biumbellatae
sect. Pachycladae
sect. Guyonianae
sect. Exiguae 2
sect. Herpetorrhizae
sect. Patellares
sect. Chylogala
sect. Calyptratae
sect. Sclerocyathium
sect. Pithyusa
sect. Myrsiniteae
sect. Helioscopia
sect. Holophyllum
sect. Lagascae
sect. Lathyris
EOCENE
OLIGOCENE
MIOCENE
PLIO.
PLEIS.
-40
-35
-30
-25
-20
-15
-10
-5
0
Age (Mya)
