## Supplementary Figure 17 for "The conquest and diversification of leafy spurges across the Holarctic and beyond: biogeography and evolution of life-history of *Euphorbia* subgenus *Esula*"

**Fig. S17**. HiSSE marginal reconstruction of extinction and relative extinction rates and life-history evolution of *Euphorbia* subg. *Esula* obtained using the R package *hisse*. Extinction rates are represented equally as colour shading along branch edges (blue to red). Life-history is represented as white (annual) or black (perennial) shading inside branches. The insets represent the distribution of extinction and relative extinction rates and character states across the tree.

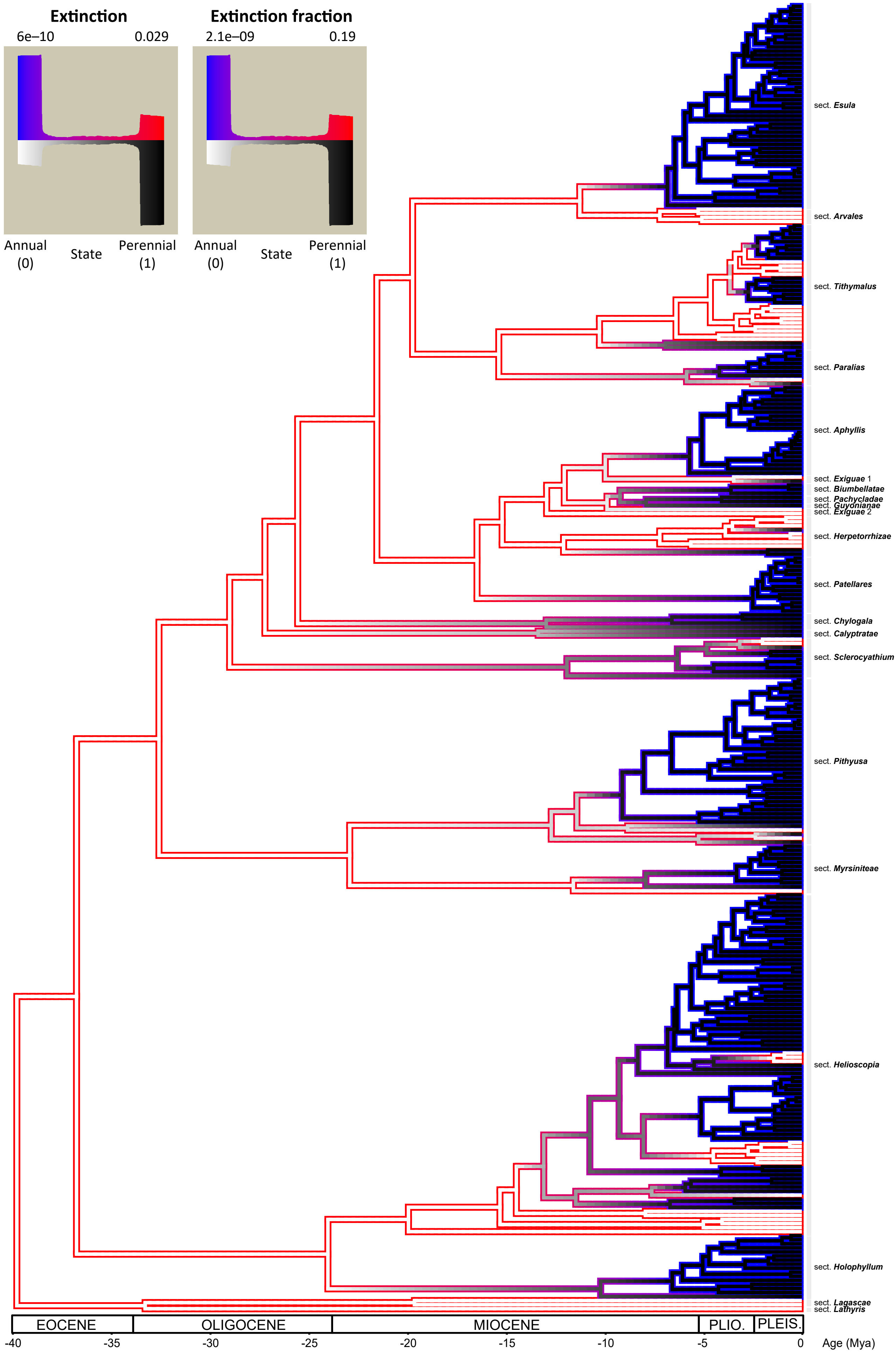

Extinction
6e–10
0.029
Extinction fraction
2.1e–09
0.19
Annual (0)
State
Perennial (1)
Annual (0)
State
Perennial (1)
sect. Esula
sect. Arvales
sect. Tithymalus
sect. Paralias
sect. Aphyllis
sect. Exiguae 1
sect. Biumbellatae
sect. Pachycladae
sect. Guyonianae
sect. Exiguae 2
sect. Herpetorrhizae
sect. Patellares
sect. Chylogala
sect. Calyptratae
sect. Sclerocyathium
sect. Pithyusa
sect. Myrsiniteae
sect. Helioscopia
sect. Holophyllum
sect. Lagascae
sect. Lathyris
EOCENE
OLIGOCENE
MIOCENE
PLIO.
PLEIS.
-40
-35
-30
-25
-20
-15
-10
-5
0
Age (Mya)
