## Supplementary Figure 18 for "The conquest and diversification of leafy spurges across the Holarctic and beyond: biogeography and evolution of life-history of *Euphorbia* subgenus *Esula*"

**Fig. S18**. Graphical proportions at different elevations (low < 1,500 m; high > 1,500 m; mixed, both ranges) of *Euphorbia* subg. *Esula* species, coded as annual or perennial.

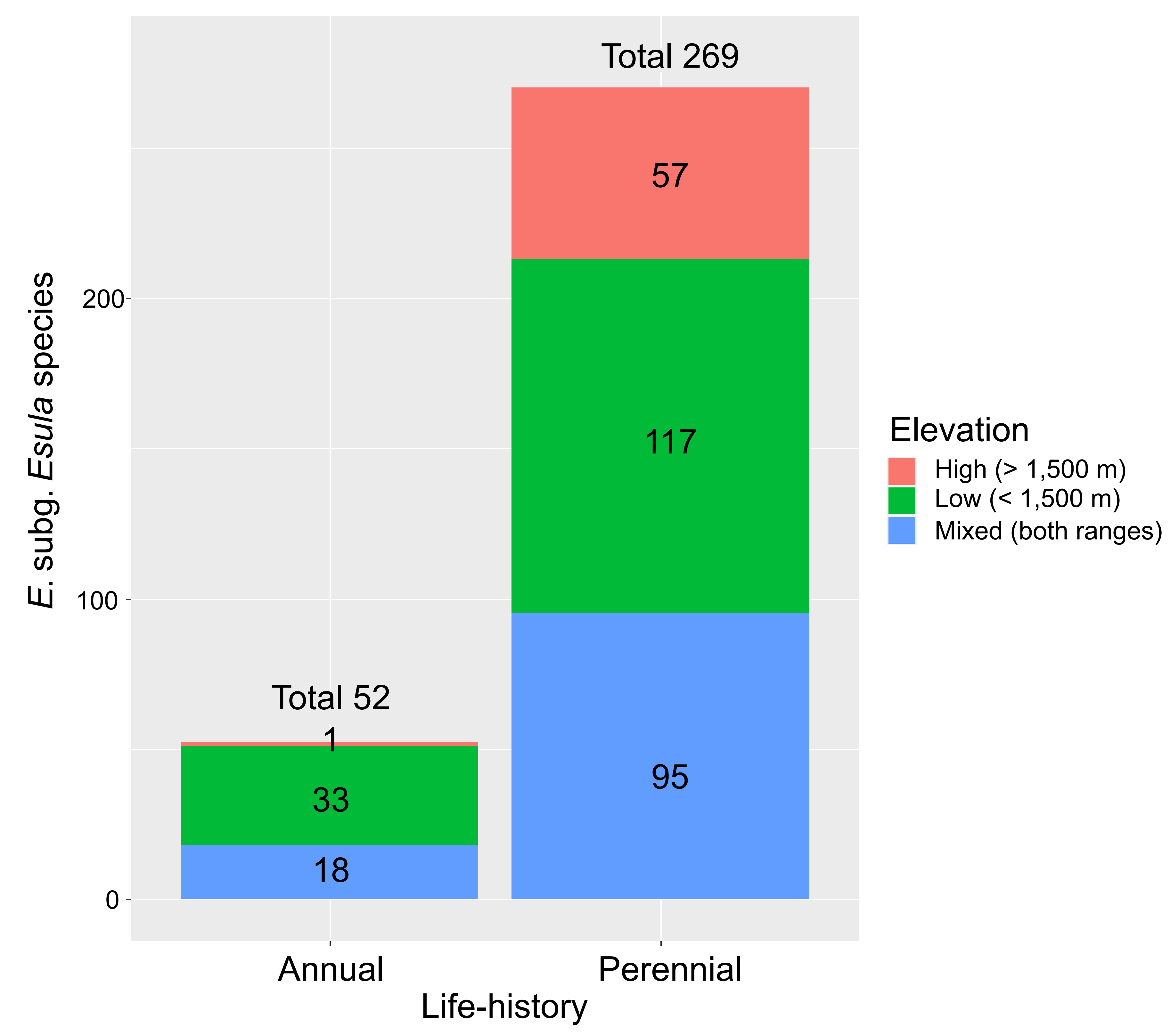
Total 269
57
117
95
Total 52
1
33
18
200
100
0
E. subg. Esula species
Annual
Perennial
Life-history
Elevation
High (> 1,500 m)
Low (< 1,500 m)
Mixed (both ranges)
